# Strong sequence dependence in RNA/DNA hybrid strand displacement kinetics

**DOI:** 10.1101/2023.11.14.567030

**Authors:** Francesca G. Smith, John P. Goertz, Molly M. Stevens, Thomas E. Ouldridge

## Abstract

Strand displacement reactions underlie dynamic nucleic acid nanotechnology. The kinetic and thermodynamic features of DNA-based displacement reactions are well understood and well predicted by current computational models. By contrast, understanding of RNA/DNA hybrid strand displacement kinetics is limited, restricting the design of increasingly complex RNA/DNA hybrid reaction networks with more tightly regulated dynamics. Given the importance of RNA as a diagnostic biomarker, and its critical role in intracellular processes, this shortfall is particularly limiting for the development of strand displacement-based therapeutics and diagnostics. Herein, we characterise 22 RNA/DNA hybrid strand displacement systems, systematically varying several common design parameters including toehold length and branch migration domain length. We observe the differences in stability between RNA-DNA hybrids and DNA-DNA duplexes have large effects on strand displacement rates, with rates for equivalent sequences differing by up to 3 orders of magnitude. Crucially, however, this effect is strongly sequence-dependent, with RNA invaders strongly favoured in a system with RNA strands of high purine content, and disfavoured in a system when the RNA strands have low purine content. These results lay the groundwork for more general design principles, allowing for creation of *de novo* reaction networks with novel complexity while maintaining predictable reaction kinetics.

## INTRODUCTION

Strand displacement reactions form the basis of most dynamic nucleic acid reaction networks and are fundamental to the field of DNA nanotechnology (1–3). Strand displacement is a nucleic acid-based process in which an incumbent strand (*Inc*) is replaced by an invader strand (*Inv*) within a complex (*SInc*) with a substrate strand (*S*). Specifically, toehold-mediated strand displacement (TMSD) reactions are a subset of strand displacement reactions that repeatedly appear in nucleic acid circuits due to their relatively high reaction rates. The presence of an overhang or ‘toehold’ motif on the substrate strand to which the invader can bind, known as the invader toehold *(γ*), provides the thermodynamic drive for forward strand displacement (Figure 1A). Hybridisation of the invader strand to the invader toehold is followed by branch migration, in which displacement of the incumbent strand occurs by a random walk process. Completion of branch migration results in two nucleic acid products: a single-stranded incumbent strand (*Inc*) and fully complementary invader-substrate complex (*SInv*).

**Figure 1.**
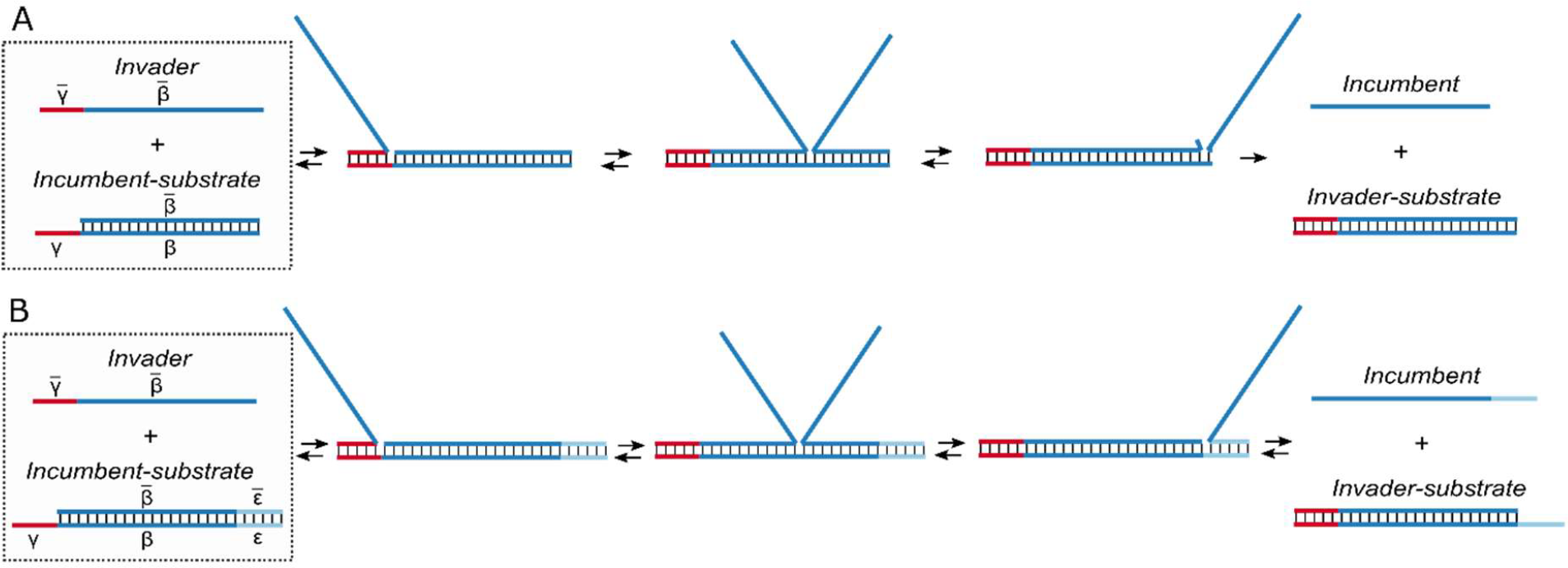
Mechanism of strand displacement reactions. A) Schematic of the mechanism of toehold-mediated strand displacement (TMSD). An invader (*Inv*) strand hybridises to a partially-complementary incumbent-substrate complex (*SInc*) via the invader toehold *(γ*) and displaces the incumbent strand within the displacement domain (*β*). B) Schematic of the mechanism of toehold exchange. The presence of a second toehold (*ε*) allows the incumbent strand to displace the invader from the invader-substrate complex (*SInv*).

The highly efficient and specific nature of TMSD reactions makes this mechanism a critical tool within dynamic nucleic acid reaction circuit designs. Moreover, the kinetic and thermodynamic properties of DNA-based (or DNA>DNA) TMSD reactions have been extensively and systemically characterised (4, 5). Zhang and Winfree (2009) pioneered systematic investigations into the effect of toehold length on the rate of TMSD. This foundational work revealed that the rate of strand displacement increases exponentially with a linear increase in toehold length up to approximately 6-7nt, above which the rate of strand displacement plateaus. The understanding obtained from this seminal work has enabled development of increasingly complex *de novo* reaction networks with predictable reaction kinetics, implemented across broad range of applications including digital nucleic acid computation (1, 2, 6), molecular motors (7), and diagnostic biosensors (3, 8, 9).

Since the foundational work by Zhang and Winfree (2009), further studies have explored the kinetics of DNA>DNA strand displacement in more detail to harness more fine-tuned kinetic control within this reaction motif. The introduction of mismatched base pairs in the branch migration domain of the invader-substrate complex has been found to decrease the rate of strand displacement by up to 4 orders of magnitude (10, 11). On the other hand, elimination of an existing mismatch in the branch migration domain of the incumbent-substrate complex can increase in the rate of strand displacement by up to 2 orders of magnitude (10, 12). Fine-tuned programmable control has equally been realised through introduction of a spacer between the invader toehold and branch migration domain, known as a remote toehold (13). This kinetic variability offers improved system flexibility and has facilitated circuit design with more regulated dynamics.

Building on the experimental work towards understanding DNA>DNA TMSD, a number of predictive models have been developed to capture strand displacement kinetics. Within the second-order limit, TMSD kinetics can be effectively described as an instantaneous, bimolecular reaction, with two products formed from two reactants via a single reaction step. Zhang and Winfree employed a simple, two-intermediate model to describe TMSD reaction kinetics (5). While this phenomenological model was able to successfully explain the exponential increase in rate with toehold length as well as the presence of a rate plateau, it was unable to provide insight into why the rate plateaued for toehold lengths of approximately 6-7nt. Subsequently, Srinivas et al. (2013) developed an ‘Intuitive Energy Landscape’ (IEL) model for TMSD, which provides a nucleotide-level understanding of strand displacement kinetics and offers useful insights from a biophysical perspective. The IEL model has since been expanded to successfully capture the effect of introduction or elimination of mismatches on reaction kinetics, supporting the previous experimental data (10).

While TMSD is frequently employed in nucleic acid reaction networks, it is limited in its use due to its irreversibility. When reversible displacement reactions are desired, the toehold exchange reaction motif is often used as an alternative. Toehold exchange involves a second toehold, known as the incumbent toehold *(ε*), that is initially sequestered in the incumbent-substrate complex but is exposed following incumbent displacement. This toehold facilitates hybridisation of the incumbent strand to the invader-substrate complex and as such allows effective reversibility (5) (Figure 1B). The second toehold enables construction of more flexible, reversible reaction networks by weakening the coupling between the thermodynamics and kinetics of such systems. There are numerous examples in which this property of toehold exchange has been exploited to achieve novel functionality including reversible logic gates (14, 15) and regulated molecular switches (16, 17).

The kinetics of toehold exchange have also been studied in depth. Systematic characterisation showed that the rate of strand displacement is effectively independent of the length of the incumbent toehold within the limit that the incumbent toehold (*ε*) is shorter than the invader toehold *(γ*). However, for *ε* > *γ*, the rate of toehold exchange decays sharply with increasing *ε* (5).

While DNA remains a critical building block in nucleic acid nanotechnology, recent years have seen increased interest in RNA-based or RNA/DNA hybrid nucleic acid nanodevices and reaction circuits (3, 9, 18–21). Despite being less stable and more costly than DNA, the structural and catalytic properties of RNA open up the field of nucleic acid nanotechnology to novel functionalities and design capabilities. Furthermore, RNA has vital roles within the cell, with direct functions in gene regulation, protein-coding, structural scaffolding and catalysis. The fundamental role for RNA in the correct functioning of cells makes it a highly informative biomarker for many diseases and developmental disorders including cancers, heart disease and neurological disorders (22–27). As such, it is unsurprising that many novel nucleic acid reaction schemes have been geared towards RNA biosensing applications in recent years (3, 9, 28, 29). Additionally, RNA-DNA hybrids have been shown to hold important roles in chromosome segregation, telomere regulation and replication regulation (30–32). These observations emphasise the importance of effectively interfacing RNA within hybrid nucleic acid reaction circuits and networks with a view to diagnostic and therapeutic applications. Moreover, *in vivo* studies have also revealed that RNA/DNA strand displacement reactions (in which RNA displaces DNA or equally DNA displaces RNA) appear to play a critical role within numerous intracellular processes including RNA transcription; genome repair, and the operation of the CRISPR/Cas machinery, highlighting the importance of gaining a clear understanding of the kinetics of this reaction in particular (33–35).

At a fundamental level, RNA>DNA (RNA displacement of DNA from a DNA-DNA duplex) strand displacement involves the separation of DNA-DNA complexes and the production of RNA-DNA hybrid complexes and vice versa in the case of DNA>RNA (DNA displacement of RNA from an RNA-DNA hybrid complex) strand displacement. The kinetics of RNA>DNA and DNA>RNA strand displacement are therefore likely to be highly dependent on the stability of RNA-DNA complexes relative to DNA-DNA complexes, a consideration that does not arise in DNA>DNA and RNA>RNA strand displacement. Moreover, it is likely that the properties of the displacement domain will play a much more important role in hybrid strand displacement than the DNA>DNA context, in which branch migration along the displacement domain replace base pairs in one duplex with identical base pairs in another.

Previous hybridisation studies of RNA-DNA hybrids point to a dependence on the purine (G/A) content within the RNA sequence. The consensus suggests that, for low purine content in the RNA strand of the hybrid, RNA-DNA hybrids are less stable than the corresponding DNA-DNA complexes. In contrast, RNA-DNA hybrids show greater stability than the equivalent DNA-DNA complex when there is a high purine content in the RNA sequence of the hybrid (36–39). Notably, many of these studies only tested short oligonucleotides and a relatively limited number of sequences were considered (36, 37). TMSD reactions allow us to effectively probe this purine dependence and quantify its effect on the thermodynamics and reaction kinetics.

Despite the importance of RNA/DNA hybrid (RNA>DNA and DNA>RNA) TMSD, and its potential complexity with respect to the DNA-based analog, understanding of RNA/DNA hybrid strand displacement kinetics remains limited. Some initial studies into RNA/DNA hybrid strand displacement have been performed (40), however thus far there is insufficient experimental data to draw consistent conclusions about the kinetics of these systems and to exploit for further computational modelling. Moreover, no studies have explicitly addressed the role of purine content on the kinetics of RNA/DNA hybrid strand displacement.

In this work we characterise RNA/DNA hybrid strand displacement reactions across a range of common design parameters including invader toehold length and branch migration domain length. We extract rate constants for each system and reveal distinct differences in strand displacement kinetics between DNA>DNA and RNA/DNA hybrid systems. We highlight the importance of the sequence composition and specifically purine content of the branch migration domain in determining the rate of hybrid strand displacement reactions, which lies in contrast to DNA>DNA strand displacement. Finally, we apply this experimental rate data to parameterise a continuous-time Markov chain model of RNA/DNA hybrid strand displacement. This model can describe strand displacement rates for RNA/DNA hybrid systems in terms of toehold length, branch migration domain length and, critically, the relative stability of RNA-DNA and DNA-DNA complexes as dictated by sequence composition.

## MATERIALS AND METHODS

### Reagents and sequence design

All sequences used in the low RNA purine content experiments (DNA_py_>DNA_py_, RNA_py_>DNA_py_, DNA_py_>RNA_py_) were designed in NUPACK (www.nupack.org) (41). Sequences were designed with minimal secondary structure. All sequences used in the high RNA purine content experiments (DNA_pu_>DNA_pu_, RNA_pu_>DNA_pu_, DNA_pu_>RNA_pu_) were adapted from Yao et al. (2015). All DNA and RNA strands were ordered from Integrated DNA technologies (IDT, Coralville, Iowa) with HPLC purification, and normalised to 100 μM in LabReady 1X IDTE buffer (pH 8.0). For labelled strands a TTT spacer was introduced between the fluorophore/quencher and the sequence of interest. All sequences were designed to have between 40% and 60% GC content. All sequences used in this work are given in Table S1 & S2. Table S3 lists all strands required to recreate each figure in this work.

### Annealing complexes

Strands were combined and heated to 95 °C for 5 minutes and then cooled to 20 °C at a constant rate of 1 °C/min to form complexes. For reporter (*FQ*) complexes, *F* and *Q* strands were combined for a final concentration of 300 nM and 360 nM, respectively. For *SInc* complexes, *S* and *Inc* strands were combined for a final concentration of 200 nM and 240 nM, respectively. We used a 20% excess of *Inc* and *Q* strands in order to ensure that all *S* and *F* strands were bound into complexes. All annealing was performed at 1M NaCl in 1X TAE (Tris-Acetate-EDTA) buffer.

### Fluorescence spectroscopy

All fluorescence measurements were performed using the BMG CLARIOstar® microplate reader. Reactions were performed in clear, flat bottom 96-well plates from Greiner Bio-One. All measurements were taken from the bottom of the wells. For all measurements, each read was an average of 20 flashes. Flashes were taken in a spiral configuration in the well with a 4mm diameter, to account for any heterogeneity within each well. For fast (well mode) reactions measurements were not taken in a spiral but at a single point at the centre of each well. Plates were maintained at the reaction temperature (25 °C) for 30 minutes prior to the first measurement to ensure that the reaction mixtures were at the correct temperature. Injection of the relevant trigger strand for each reaction was performed using the automatic injection feature of the CLARIOstar® microplate reader at a pump speed of 430 μL/s. After each injection, samples were shaken for 6 s at a speed of 400 rpm in a double-orbital configuration. The injectors were passivated with 5% BSA for 20 minutes prior to being loaded with DNA or RNA samples to minimise loss of the nucleic acid and maximise reproducibility. Plates were sealed using Thermofisher Adhesive PCR Plate Seals and kept for up to a month in a shaking incubator at the reaction temperature and 70 rpm to allow the reaction to reach equilibrium. *F* was labelled with AlexaFluor488 (excitation: 488 nm; emission: 496 nm) and *Q* was labelled with IowaBlack FQ®. An excitation window of 488-14 and emission window of 535-30 were used. The same focal height (5.9 mm) was used across all experiments to allow for comparison between experiments. The gain was set to the same value as that used for calibration curve measurements to ensure comparability (the gain used for each calibration curve are given in Figure S2).

### General procedure for experimental strand displacement characterisation

Experimental protocols are adapted from (42). All experiments have a final reaction volume of 200 μL. All experiments were composed of multiple measurements that each capture either the reaction kinetics or allow for estimation of reactant or product concentrations. All experiments included a positive control with 15 nM *FQ* and an excess (20 nM) *Inc*; and a buffer-only negative control. All reactions were measured in at least triplicate to obtain an estimate of the error within our measurements. Of note, the concentrations given are intended concentrations of reactants, however for fitting purposes we determined the exact concentrations for each reactant based on fluorescence output after calibration.

#### Fluorescence conversion calibration

Calibration curves were generated to allow for conversion of fluorescence in arbitrary fluorescence units (afu) to concentration of fluorescent product (*FInc*) in nM to quantify the concentration of reacted *FQ* in each assay. Final reaction volumes of 200 μL were used for all calibration experiments. Calibration curves were generated by reacting 15 nM of *FQ* with varying concentrations of *Inc* (2-20 nM for the low RNA purine content system and 4-30 nM for the high RNA purine content system). An additional negative control with 0 nM *Inc* was also included for all experiments. Further details of the calibration experiment protocols are given in supplementary note 6.1 (full protocol in Table S4 & S5). Details of the results of calibration experiments are given in Figure S2 and supplementary note 6.3.

#### Reporter characterisation reactions

Reporter characterisation reactions were used to estimate the reporter rate constant, *k*_rep_, for each *FQ* and *Inc* combination used in this work. Reporter reactions are described as

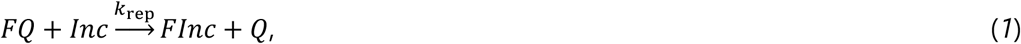

where *FInc* represents the fluorescent product. We can fit a simple ordinary differential equation (ODE) model to the normalised reaction data in order to extract estimates for the reporter rate constant (*k*_rep_). Using conservation laws, the reporter reaction can be reduced to a single ODE

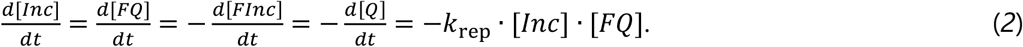

For these experiments we used [*FQ*] = 15 nM and [*Inc*] = 4-12 nM. The majority of *FQ* complexes used herein were designed with a 6nt toehold (RNA>DNA and DNA>DNA strand displacement reactions for the high RNA purine content system used an *FQ* reporter toehold of 8nt). All reactions were saturated with an excess concentration of *Inc* after reaction kinetics were completed to determine [*FQ*] at time t = 0 s. An additional negative control was included with [*FQ*] = 15 nM and [*Inc*] = 0 nM. Further details of experimental protocol for reporter characterisation are given in supplementary note 7.1 (full protocol in Table S6, illustrative fluorescent data in Figure S3). Further details of the normalisation and estimation of *k*_rep_ are given in supplementary note 7.2 and supplementary note 7.3, respectively. All fluorescent traces for reporter characterisation are given in Figure S4, fits of the ODE model to the fluorescent traces are given in Figure S5 & S6 and estimated *k*_rep_ values are given in Table S7 & S8.

#### Characterisation of full toehold exchange reactions

We estimated the effective rate constant, *k*_eff_, for each toehold exchange system that was tested. We model toehold exchange reactions as a second-order reaction

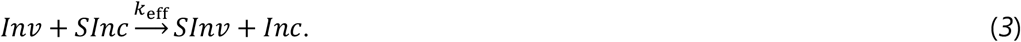

We combine both the toehold exchange and reporter displacement reactions to generate a system of ODEs. Using conservation laws this series of reactions can be well described by 3 ODEs

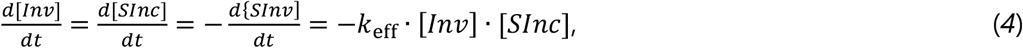

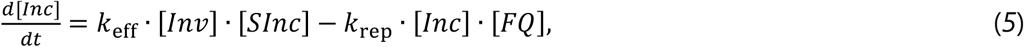

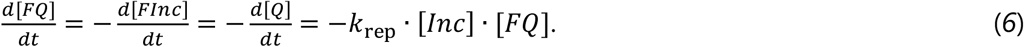

Note that equation (6) is equivalent to equation (2) above.

For each design we estimated *k*_eff_ for DNA>DNA, RNA>DNA and DNA>RNA displacement reactions. These experiments were performed by combining [*FQ*] = 15 nM, [*SInc*] = 10 nM and [*Inv*] = 4-8 nM (for DNA_pu_>RNA_pu_ strand displacement reactions we used [*Inv*] = 40-80 nM). We used an excess of [*FQ*] and [*SInc*], unless specified otherwise, to ensure that all [*Inv*] reacted and we were able to effectively estimate [*Inv*] at time, t = 0, [*Inv*]_0_. For these experiments we used 3 different [*Inv*] concentrations to confirm the robustness of the second-order assumption and each reaction was performed in triplicate. All reactions were saturated with an excess concentration of *Inv* after reactions kinetics were complete to determine [*SInc*] at time t = 0 s. Reactions were next saturated with an excess concentration of *Inc* to determine [*FQ*] at time t = 0 s. An additional negative control was included with [*FQ*] = 15 nM, [*SInc*] = 10 nM and [*Inv*] = 0 nM. Further details of the experimental protocol for toehold exchange characterisation are given in supplementary note 8.1 (full protocol in Table S9 & S10, illustrative fluorescent data in Figure S7 & S8). Further details of the normalisation and estimation of *k*_eff_ are given in supplementary note 8.2 and supplementary note 8.3, respectively. All fluorescent traces for toehold exchange characterisation are given in Figures S10-S12, fits of the ODE model to the fluorescent traces are given in Figures S13-S18 and fitted *k*_eff_ values are given in Table S12 & S13.

#### Characterisation of leak reactions

We observed an undesired leak reaction between *SInc* and *FQ* for DNA_pu_>RNA_pu_ strand displacement reactions. We describe the undesired leak reaction as

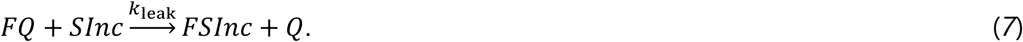

For these strand displacement reactions we adapted the experimental protocol to effectively estimate *k*_eff_ despite the presence of the leak reaction. These experiments were performed by combining [*FQ*] = 15 nM, [*SInc*] = 10 nM and [*Inv*] = 40-80 nM. An excess of *Inc* is also present from formation of the *SInc* complexes. All reactions were performed in triplicate. All reactions were saturated with a further large excess concentration of *Inv* to ensure all *SInc* complexes were dissociated and therefore determine [*SInc*] at time t = 0 s. Reactions were next saturated with an excess concentration of *Inc* to determine [*FQ*] at time t = 0 s. An additional negative control was included with [*FQ*] = 15 nM, [*SInc*] = 10 nM and [*Inv*] = 0 nM. We first estimated the leak reaction rate constant, *k*_leak_, from this negative control reaction using the following ODEs

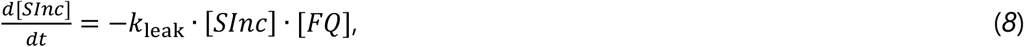

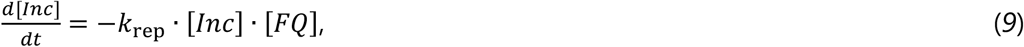

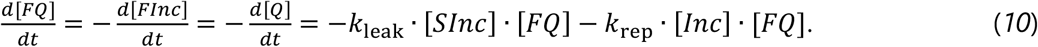

Note that the right hand side of equation (*9*) is equivalent to equations (*2*) and (*6*) above. We include the ODE describing the reporter displacement reaction due to the presence of excess *Inc*.

For the toehold exchange reactions of interest we then estimated *k*_eff_ while keeping the corresponding *k*_leak_ estimate fixed. We adapted the ODE model in equations (*4*) and (*5*) to account for the undesired leak reaction observed for DNA>RNA strand displacement

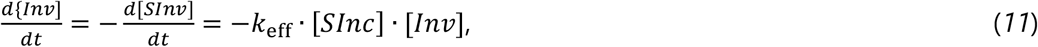

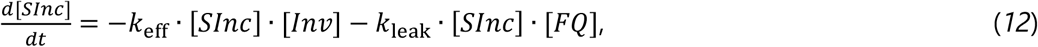

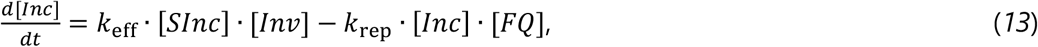

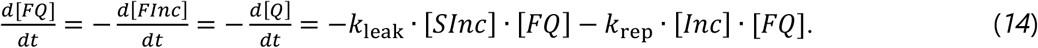

Note that the right hand side of equation (*11*) is equivalent to equation (*4*) above and the right hand side of equation (*14*) is equivalent to equation (*10*) above.

Further details of experimental protocol for toehold exchange characterisation for these leak reactions are given in supplementary note 8.1 (full protocol in Table S11, illustrative fluorescent data in Figure S9). Further details of the normalisation are given in supplementary note 8.3. Further details of *k*_leak_ and *k*_eff_ estimation are given in supplementary note 8.4. All fluorescent traces for toehold exchange characterisation for systems with a leak are given in Figure S12, fits of the ODE model to the fluorescent traces are given in Figures S19 & S20 and fitted *k*_leak_ and *k*_eff_ values are given in Table S14 and Table S15, respectively.

### General fitting procedure

All fitting was performed in Python version 3.8.8. First, kinetic traces for fluorescence in afu were converted to [*FInc*] in nM using the calibration results, then least-squares fitting of the ODE model was used to estimate relevant rate constants. Initial reactant concentration parameters, which may deviate from intended values due to experimental imperfections, were calculated prior to fitting using the calibration results.

For reporter characterisation data we estimated the rate constant *k*_rep_. For each experiment we derived a single, global estimate of *k*_rep_ across all replicates and each initial concentration of *Inc* at time t = 0 s ([*Inc*]_0_). [*Inc*]_0_ and [*FQ*]_0_ were fixed values in the fitting protocol. The *k*_rep_ values estimated in the reporter characterisation experiments were fixed for fitting the corresponding toehold exchange reactions. Similarly, the values of *k*_leak_ obtained from preliminary experiments (see supplementary note 8.4) were fixed for fitting the corresponding DNA_pu_>RNA_pu_ strand displacement reactions.

We estimated *k*_eff_ for each toehold exchange system that we tested. [*Inv*]_0_, [*SInc*]_0_ and [*FQ*]_0_ were fixed values in the fitting protocol. Unless otherwise specified for each experiment we estimated a single, global estimate for *k*_eff_ across all replicates and [*Inv*]_0_.

For DNA_pu_>RNA_pu_ strand displacement reactions we were unable to estimate global *k*_eff_ values. In this case, for each experiment we assumed that [*Inv*]_0_ matched the intended concentration and calculated the individual estimates of *k*_eff_ for each experimental trace. We report the mean of these *k*_eff_ values instead of a global fit of *k*_eff_. Fitting individual estimates of *k*_eff_ for each experimental trace for all other sets of experiments produced results that agreed well with the global fits. Supplementary note 7.3 contains a detailed description of the fitting procedures for reporter characterisation experiments. Supplementary notes 8.3 and 8.4 contain a detailed description of the fitting procedures for full strand displacement experiments.

### Initial rate predictions using IEL model

Initial predictions from the IEL model were based on previous parameters for DNA>DNA strand displacement (Δ*G*_assoc_, Δ*G*_bp_, Δ*G*_p_, Δ*G*_bm_ and *k*_bp_). The values of each free-energy parameter and rate parameter are given in Table 1, reproduced from (10). For initial predictions Δ*G*_rd_, the relative stability of RNA/DNA and DNA/DNA duplexes, is a flexible parameter. For these predictions we used an initial invader concentration of 6 nM, with *γ* = 4nt, *β* = 26nt and *ε* = 4nt (Figure S1). We assumed a reaction temperature of 25 °C (298 K). Full details of the IEL model are given in supplementary notes 1-3. Detailed explanation of these initial predictions are given in supplementary note 4.

**Table 1.** Estimated parameter sets to describe RNA/DNA hybrid strand displacement reactions for low RNA purine content design. Estimated parameters for the IEL model obtained from fits of RNA_py_>DNA_py_, DNA_py_>DNA_py_ and DNA_py_>RNA_py_ toehold exchange experimental data. The bottom two rows are a comparison for the equivalent parameters from previous studies of DNA>DNA strand displacement systems (10, 11). Parameter values in the final row of the table were derived by Irmisch et al. (2020) from data produced by Machinek et al. (2014). For *k*_bp_ we provide lower and upper confidence intervals (*CI*) as this value was estimated as log _0_(*k*_bp_). The available data does not strongly constrain all parameters, with similar fits arising when subsets of parameters were varied together therefore parameters marked with an asterisk (*) were fixed at the values from Irmisch et al. (2020) to allow for appropriate parameter estimation.

| Data set | $\Delta G_{bp}$<br>($k_B T$) | $\Delta G_{assoc}$<br>($k_B T$) | $\Delta G_p$<br>($k_B T$) | $\Delta G_{bm}$<br>($k_B T$) | $k_{bp}$ ($s^{-1}$) | $\Delta G_{rd}$ ($k_B T$) |
| --- | --- | --- | --- | --- | --- | --- |
| This study | -2.52* | 2.5* | 3.5* | $9.3 \pm 1.2$ | $6.4$ (*CI*: 2.3;<br>18.3) $\cdot 10^7$ | $0.16 \pm 0.04$ |
| Irmisch | -2.52 | $2.5 \pm 0.2$ | $3.5 \pm 0.2$ | $7.4 \pm 0.2$ | $(5.9 \pm 1.1) \cdot 10^7$ | - |
| Machinek | -2.51 | $4.0 \pm 0.2$ | $5.4 \pm 0.4$ | $8.5 \pm 0.3$ | $(20.6 \pm 0.6)$<br>$\cdot 10^7$ | - |

### Parameterisation of IEL model for RNA/DNA hybrid strand displacement

Parameterisation of the free-energy landscape model was performed in Python version 3.8.8. For parameterisation based on the low RNA purine content (DNA_py_>DNA_py_, RNA_py_>DNA_py_, DNA_py_>RNA_py_) kinetic data Δ*G*_assoc_, Δ*G*_p_ and Δ*G*_bp_ were fixed at the previously estimated values given in Table 1, while Δ*G*_bm_, *k*_bp_ and Δ*G*_rd_ were estimated. For parameterisation using the high RNA purine content (DNA_pu_>DNA_pu_, RNA_pu_>DNA_pu_, DNA_pu_>RNA_pu_) kinetic data Δ*G*_assoc_, Δ*G*_p_, Δ*G*_bp_ and Δ*G*_bm_ were fixed at the values predicted from the low RNA purine content kinetic data, while *k*_bp_ and Δ*G*_rd_ were flexible parameters. For parameterisation the reaction temperature was set to 25 °C (298 K). Parameterisation was performed by minimising the squared residuals between the model-predicted and the experimentally-derived values of *k*_eff_. The details of the parameterisation protocol are given in supplementary note 9.

## RESULTS

### RNA/DNA hybrid free-energy landscape model

In this work we have developed a continuous-time, Markov chain model to describe the rate of strand displacement for RNA/DNA hybrid systems. This model has been adapted from the Intuitive Energy Landscape (IEL) model developed by Srinivas et al. (2013). The IEL model was initially introduced to explain DNA>DNA strand displacement reaction kinetics (43). More recently it has been expanded to capture the effect of mismatches in strand displacement systems (10). We further expand on the IEL model as described by Irmisch et al. (2020) to allow prediction of RNA>DNA and DNA>RNA strand displacement kinetics (10). For all systems referred to here the substrate strand (S) was composed of DNA.

Our free-energy landscape model is shown in Figure 2A. This model contains a number of distinct states which represent each step in the strand displacement reaction, from state –*γ* through to state *N*. State –*γ* refers to the stage in which the invader is unbound from the incumbent-substrate complex. Completion of toehold hybridisation is defined as state 0, in which in the invader is bound to the incumbent-substrate complex via the toehold but branch migration has not been initiated. Branch migration proceeds through to state *β* in which the incumbent has been successfully displaced by the invader strand. For systems with no incumbent toehold state *β*, is equivalent to state *N*. For systems in which the incumbent toehold length is non-zero (*ε* > 0 as in Figure 1B), state *N* refers to the complete dissociation of the incumbent strand from the incumbent toehold of the invader-substrate complex.

**Figure 2.**
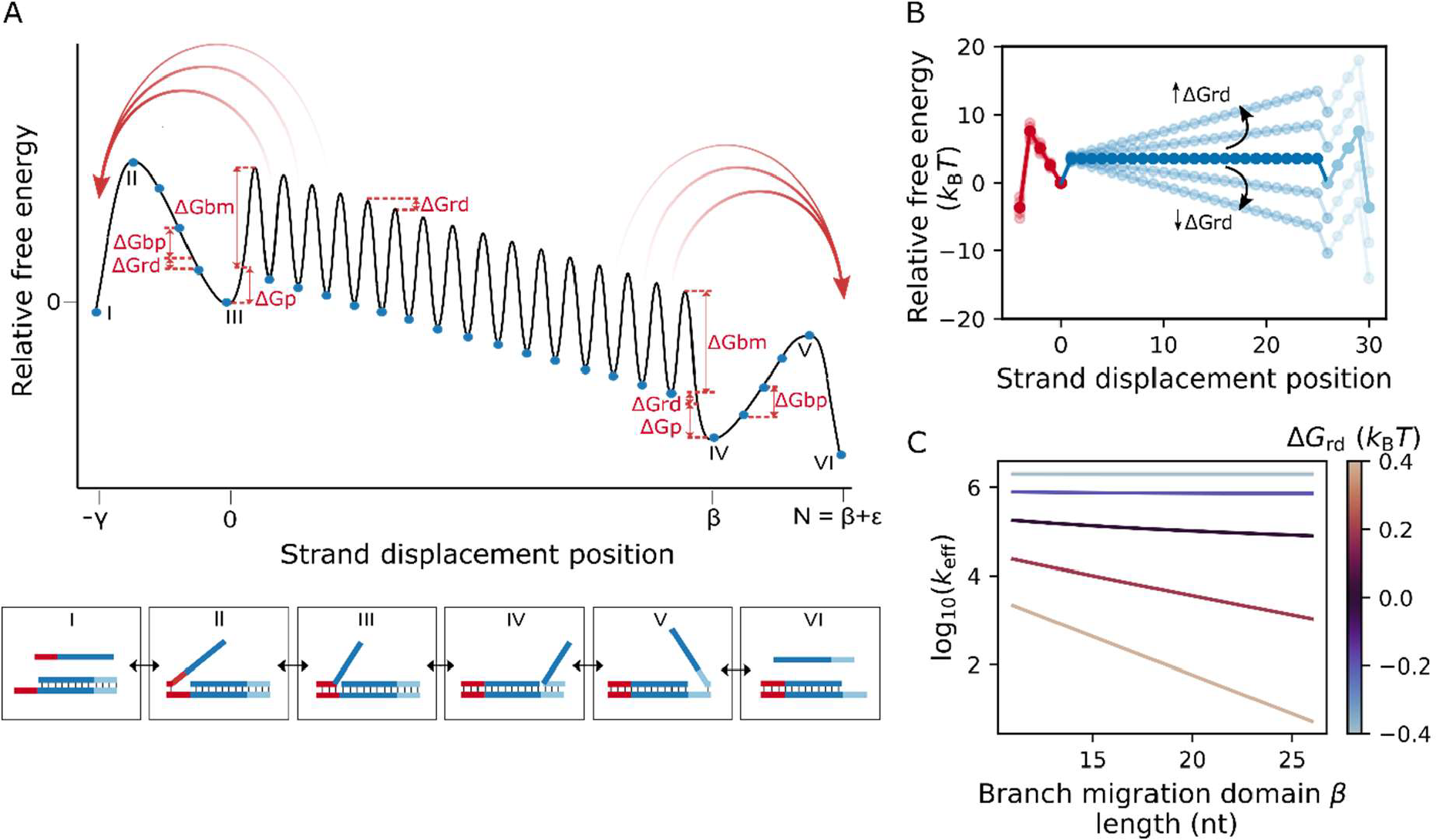
Free-energy landscape model for toehold exchange. A) Schematic of a simplified free-energy landscape for a thermodynamically downhill toehold exchange reaction, based on the IEL model (10). The total length of the substrate strand is denoted as *N* + *γ*, with invader toehold length*γ*, branch migration domain length *β* and incumbent toehold length *ε*. The strand displacement position defines the state of the system; it is the position on the substrate strand at which the leading invader-substrate base pair has formed. Within the diagram, these states are represented by blue dots; the sawtooth represents relatively large barriers inhibiting transitions between these states on the branch migration domain. Specific states are: I) unbound invader and incumbent-substrate complex; II) single invader nucleotide hybridised to the invader toehold; III) invader strand fully hybridised to the invader toehold; IV) invader strand fully displaced incumbent strand in branch migration domain; V) single incumbent nucleotide bound to the incumbent toehold; VI) unbound incumbent and invader-substrate complex. Free energies are given relative to a reference state with free energy of 0 *k*_B_*T* in which the invader is fully bound to the toehold, and a perfectly-matched incumbent strand is bound to the branch migration domain (state III). Curved, red arrows represent spontaneous invader and incumbent dissociation, and straight, double-headed red arrows reflect free-energy changes. The contribution of model parameters to the free-energy landscape is illustrated. B) The value of Δ*G*_rd_ affects gradient of the free-energy landscape within the branch migration domain. For RNA>DNA strand displacement, negative Δ*G*_rd_ values produce an downhill landscape, positive Δ*G*_rd_ values produce an uphill landscape, while Δ*G*_rd_ values of 0 *k*_B_*T* give a flat landscape in the displacement domain (solid coloured landscape). Each state in the free-energy landscape is denoted as a dot. C) For RNA>DNA toehold exchange, we predict the effective rate constant (*k*_eff_) for an increase in *β* = 11nt to *β* = 26nt, across a range of Δ*G*_rd_ = −0.4 *k*_B_*T* to Δ*G*_rd_ = 0.4 *k*_B_*T* for *γ* = 4nt and *ε* = 4nt. *Inc*reasing Δ*G*_rd_ results in increasing rate dependence with branch migration domain length.

The IEL model as defined by Irmisch et al. (2020) fully describes non-mismatched DNA>DNA strand displacement reactions with 4 free-energy parameters and a single rate parameter: Δ*G*_assoc_, Δ*G*_bp_, Δ*G*_p_, Δ*G*_bm_ and *k*_bp_ (10). Δ*G*_assoc_ is the free-energy penalty caused by the initial association of the invader to incumbent-substrate complex within the toehold resulting in reduced orientational and translational freedom of the single strand at the standard 1 M reference concentration. Δ*G*_bp_ corresponds to the free-energy change resulting from formation of a single additional DNA-DNA base pair within a duplex. Therefore, toehold hybridisation can be expressed as a free-energy reduction by Δ*G*_bp_ for each base pair formed in this process. For simplicity, within this model we ignore the individual contribution of the different nucleotides and assume a sequence-average value for Δ*G*_bp_. Δ*G*_p_ defines the free-energy penalty associated with the initiation of branch migration due to the presence of two single-stranded overhangs as the incumbent strand starts to be displaced. Δ*G*_bm_ is the free-energy barrier associated with each branch migration step. This parameter can be thought of as the transition barrier to breaking a single incumbent-substrate base pair and formation of a single invader-substrate base pair, due to the rearrangement of invader and incumbent strands involved in displacement (43). We assume that for DNA>DNA strand displacement reactions there is no net free-energy change between adjacent states during branch migration. These 4 free-energy parameters describe a one-dimensional free-energy landscape for a non-mismatched DNA>DNA strand displacement reaction. Notably, the IEL model as described by Irmisch et al. (2020) also accounts for spontaneous incumbent dissociation before state *N* is reached. The probability of spontaneous detachment of the incumbent strand is assumed to drop off exponentially with an increase in the number of remaining base pairs with the substrate strand. Expanding the one-dimensional landscape model to include spontaneous incumbent dissociation has been shown to improve predictions of experimental strand displacement kinetic data (10, 11).

In this study we introduce an additional free-energy parameter, Δ*G*_rd_, which describes the free-energy difference between DNA-DNA and RNA-DNA base pairs. Using these 5 free-energy parameters the free-energy change between each state was calculated in order to construct complete free-energy landscapes for DNA>DNA, RNA>DNA and DNA>RNA strand displacement reactions.

Using an analytical solution of our model we calculated the predicted rate of reaction from the free-energy landscape for each system (supplementary note 3). We first calculated the first passage time ⟨*t*⟩, which is defined as the time taken to pass from state –*γ* through to state *N* for each system. Within the second order limit the first passage time is related to the effective second-order rate constant (*k*_eff_) according to

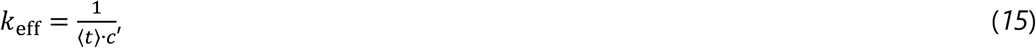

where *c* is the initial free invader concentration.

We made preliminary *k*_eff_ predictions for RNA>DNA and DNA>RNA toehold exchange reactions across a range of toehold lengths and branch migration domain lengths. For initial predictions we made use of free-energy parameter values extracted from previous work of DNA>DNA strand displacement kinetics and allowed Δ*G*_rd_ to remain a flexible parameter (10). The IEL model suggests that for strand displacement reactions with no net free-energy change between states during branch migration (Δ*G*_rd_ = 0 *k*_B_*T*) the rate of strand displacement decreases with an increase in the branch migration domain length, *β*, according to

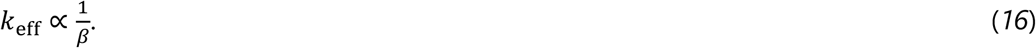

Previous studies of hybridisation within the literature suggest that RNA-DNA hybrids are less stable than the corresponding DNA-DNA complexes for systems with a low purine content in the RNA strand of the hybrid, while a high RNA purine content produces the opposite thermodynamic trend (36–39). Under this assumption, we would expect RNA displacing DNA (RNA>DNA) and DNA displacing RNA (DNA>RNA) to exhibit an overall uphill free-energy landscape for low and high RNA purine content systems, respectively (Figure 2B). Within these limits, our initial predictions suggest an approximately exponential decrease in effective rate constant with a linear in *β* (Figure 2C). By contrast, we anticipate RNA>DNA and DNA>RNA strand displacement reactions to exhibit an overall downhill free-energy landscape for high and low RNA purine content systems, respectively. In this case preliminary predictions indicate a very weak dependence of the effective rate constant on *β* for branch migration domain lengths greater than 10nt. These initial predictions suggest that the branch migration domain length is a critical parameter in effectively extracting Δ*G*_rd_, displaying significantly different trends between uphill and downhill landscapes (Figure S1).

Therefore, we focussed on systems with differing branch migration domain lengths to probe the relative stability of RNA-DNA and DNA-DNA base pairs, and extract a useful estimate for Δ*G*_rd_ for RNA>DNA and DNA>RNA strand displacement reactions across systems of purine content extremes.

### Experimental characterisation of RNA/DNA hybrid strand displacement reactions

We designed a toehold exchange reaction network to probe the reaction rates for RNA>DNA and DNA>RNA strand displacement systems. These toehold exchange reactions were monitored over time by assessing the concentration of released incumbent strand over the course of the reaction. We made use of a distinct fluorescent reporter detection system (Figure 3A) to assess the concentration of released incumbent to avoid directly labelling the strands within the reaction of interest, which may otherwise influence the kinetics of strand displacement (44). The reporter complex was composed of a fluorescently (AlexaFluor488)-labelled strand (*F*) which was partially complementary to a quencher (IowaBlack-FQ)-associated strand (*Q*). The reporter complex was designed such that F had a 5’ overhang of 6nt or 8nt, which acted as the reporter toehold (*δ*). The released incumbent strand (*Inc*) possessed a domain complementary to the toehold *δ*, allowing the incumbent to displace the quencher-associated strand, resulting in a detectable fluorescent product (*FInc*). Across all the systems, we designed the reporter reaction to be effectively instantaneous compared to the toehold exchange reaction of interest such that the reporter acted as an effective readout for the toehold exchange kinetics.

**Figure 3.**
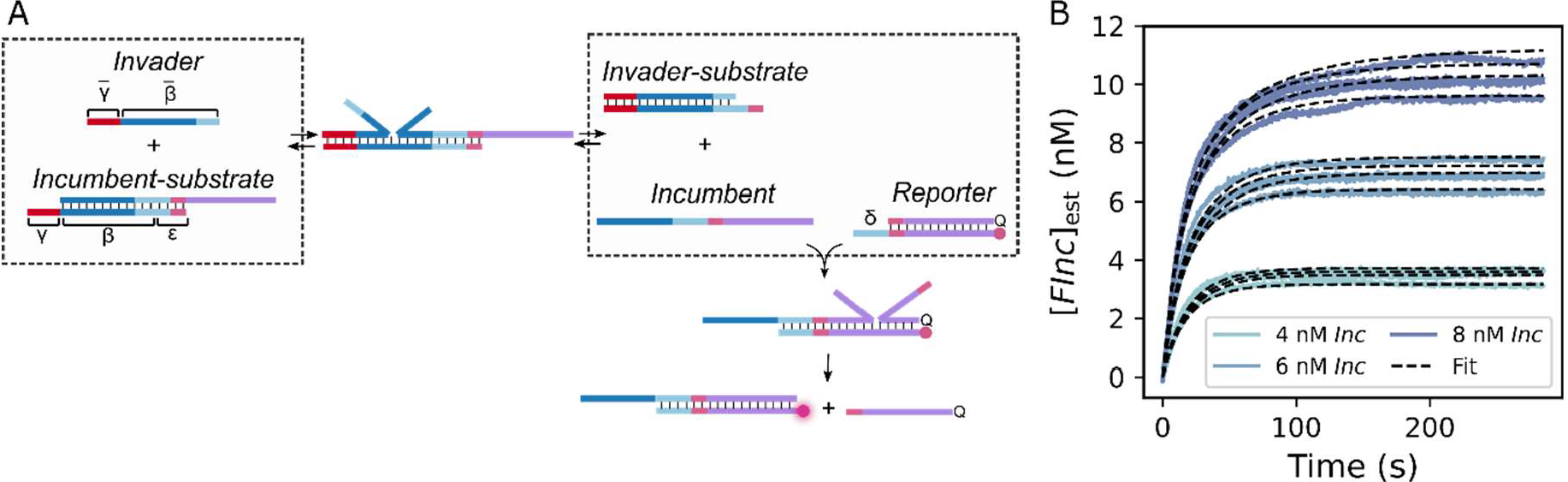
Reporter detection system for experimental characterisation of toehold exchange reactions. A) Domain-level schematic of toehold exchange reaction network, where *γ* represents the invader toehold, *β* represents the branch migration domain, *ε* represents the incumbent toehold and *δ* represents the reporter toehold. Barred letters represent complementary sequences to their unbarred counterparts. Toehold exchange occurs between the invader strand and the incumbent-substrate complex with second-order rate constant *k*_eff_. The released incumbent strand induces fluorescence by displacing the quencher-associated strand (Q) from a complex with a fluorescently-labelled strand (F), with second-order rate constant *k*_rep_. B) Example fluorescent traces of *FInc* production over time across a range of *Inc* concentrations (4-12 nM) to characterise *k*_rep_ for a DNA *Inc* strand with *β* = 26nt (low RNA purine content design). Coloured, solid lines show the normalised fluorescent traces. Dashed, black lines represent output of fitting an ODE model to this fluorescent data.

For each design we assessed RNA and DNA counterparts of invader and incumbent strands, however for all systems herein only DNA substrate strand and DNA-DNA reporter complexes were used. Importantly, all sequences were designed to minimise secondary structure which may otherwise have affected the strand displacement rate, and confounded estimation of *k*_eff_. In this work, we designed one system with low purine content in the invader strand (17% G/A) and designed a second system with high purine content in the invader strand (78% G/A). For these two independently-designed systems we aimed to extract the effective rate constant for DNA>DNA, RNA>DNA and DNA>RNA strand displacement reactions across a range of invader toehold lengths (4nt-10nt) and branch migration domain lengths (11nt-26nt) or (11nt-21nt for the high RNA purine content design). We refer to RNA>DNA strand displacement for the design with high purine content in the RNA invader strand (and by inference high purine content in the incumbent strand) as RNA_pu_>DNA_pu_ and equivalently for DNA>RNA strand displacement as DNA_pu_>RNA_pu_. For systems with low purine content in the invader (and by inference high pyrimidine content), we use RNA_py_>DNA_py_ and DNA_py_>RNA_py_. In order to minimise interaction and limit potential sequestration of the reporter toehold by the invader we ensured that only 4nt of the invader strand overlapped with the reporter toehold. We also introduced a GC-rich clamp in both the incumbent-substrate and reporter complexes to minimise undesired leak reactions. This design resulted in an incumbent toehold (*ε*) of 4nt for all systems.

To minimise cost we designed our system such that only two RNA or DNA invaders were required across all toehold length and branch migration domain length combinations: one for the low RNA purine content design and one for the high RNA purine content design. For the low RNA purine content design an invader with *γ* = 10nt and *β* = 26nt was used, while the toehold length and branch migration domain length of the substrate and incumbent strands were adjusted. For the high RNA purine content design an invader with *γ* = 6nt and *β* = 21nt was used, while the branch migration domain length of the substrate and incumbent strands were adjusted to achieve the desired domain lengths.

We initially performed a series of reporter characterisation reactions to estimate the rate of TMSD for the reporter reaction (*k*_rep_). For these experiments we combined the 15 nM of *FQ* with *Inc* at a range of concentrations (Figure 3B). Across all the reporters that were tested we showed that the reporter strand displacement reaction was sufficiently rapid to act as an effective readout for the reaction of interest, confirming the suitability of these designs. We estimated *k*_rep_ for each reporter reaction by fitting an ODE model to the fluorescence data (supplementary note 7.3 and Figure S6). Final estimates of *k*_rep_ for all designs are given in Table S8.

We next monitored the fluorescence for the complete strand displacement network for the low purine content design (Figure 4A). We initially assessed the effect on *k*_eff_ of changing the invader toehold length (4nt – 10nt) for a fixed branch migration domain length of 26nt.

**Figure 4.**
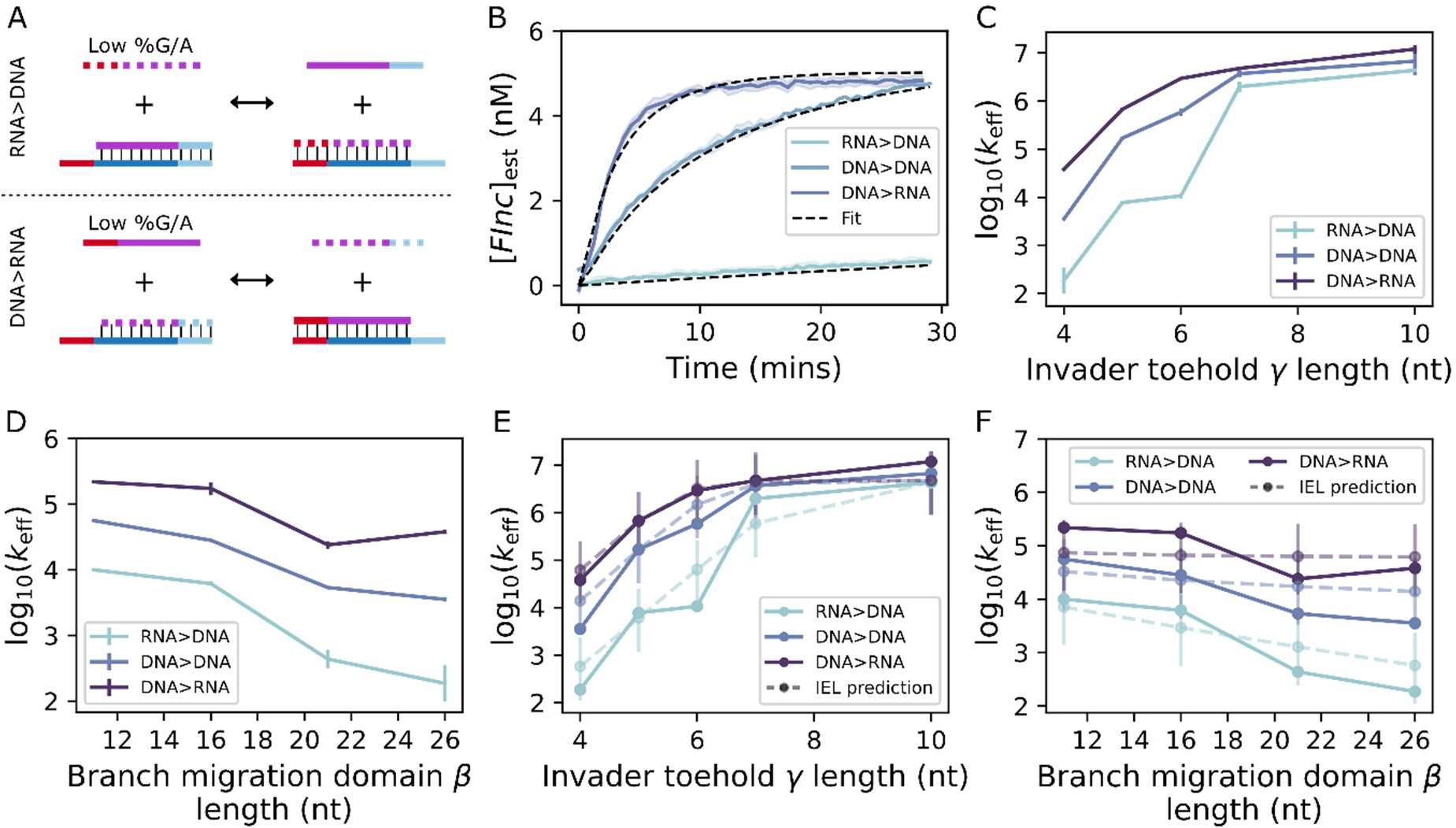
RNA>DNA and DNA>RNA toehold exchange kinetics for low RNA purine content design. A) Domain-level schematic of RNA_py_>DNA_py_ and DNA_py_>RNA_py_ toehold exchange. B) Example fluorescent traces for RNA_py_>DNA_py_, DNA_py_>DNA_py_ and DNA_py_>RNA_py_ toehold exchange system with *γ* = 5nt, *β* = 26nt and *ε* = 4nt for an *Inv* concentration of 6 nM, across 3 replicates. Solid, coloured lines represent normalised fluorescent curves. Dashed, black lines represent output of fitting an ODE model to this kinetic data. C) Summary of experimentally-derived *k*_eff_ values for RNA_py_>DNA_py_, DNA_py_>DNA_py_ and DNA_py_>RNA_py_ toehold exchange reactions across a range of invader toehold lengths (4-10nt) for a fixed *β* = 26nt and *ε* = 4nt. Error bars show 95% confidence intervals based on the standard error of *k*_eff_ estimates from individual curves. D) Summary of experimentally-derived *k*_eff_ values for RNA_py_>DNA_py_, DNA_py_>DNA_py_ and DNA_py_>RNA_py_ toehold exchange reactions across a range of branch migration domain lengths (11-26nt) for a fixed *γ* = 4nt and *ε* = 4nt. Error bars show 95% confidence intervals based on the standard error of *k*_eff_ estimates from individual curves. N-3. E) Predicted *k*_eff_ from parameterised IEL model (dashed lines) compared to experimentally-derived values (solid lines) across invader toehold lengths (4-10nt). Error bars show 95% confidence intervals of model-predicted *k*_eff_ values. N=3. F) Predicted *k*_eff_ from parameterised IEL model (dashed lines) compared to experimentally-derived values (solid lines) across branch migration domain lengths (11-26nt). Error bars show 95% confidence intervals of model-predicted *k*_eff_ values. N=3.

Reactions were initiated by combining 4 nM, 6 nM or 8 nM of *Inv* with 10 nM of *SInc* and 15 nM of *FQ*. We then estimated *k*_eff_ by fitting a simple ODE model to the fluorescent traces using a non-linear least-squares approach (supplementary note 8.3, Figures S16-18) (Figure 4B). We found that the estimated *k*_eff_ values (Figure 4C) for DNA_py_>DNA_py_ toehold exchange were in good agreement with experimental data from previous studies, with the rate appearing to plateau at an invader toehold length between 6nt and 7nt (5, 10). Both DNA_py_>RNA_py_ and RNA_py_>DNA_py_ toehold exchange systems appeared to follow the same general trend as DNA_py_>DNA_py_ toehold exchange. However, RNA_py_>DNA_py_ and DNA_py_>RNA_py_ strand displacement rates plateaued at toehold lengths closer to 7nt and 6nt, respectively. These results also reveal that RNA_py_>DNA_py_ strand displacement is up to an order of magnitude slower than the corresponding DNA_py_>DNA_py_ strand displacement reaction. In contrast, DNA_py_>RNA_py_ strand displacement is up to an order of magnitude faster than DNA_py_>DNA_py_ strand displacement and up to two orders of magnitude faster than RNA_py_>DNA_py_ strand displacement. Numerical estimates of *k*_eff_ for RNA_py_>DNA_py_, DNA_py_>DNA_py_ and DNA_py_>RNA_py_ strand displacement reactions are given in Table S13.

Subsequently, we investigated the effect of branch migration domain length on the rate of strand displacement. A 4nt invader toehold was used and a total of 4 branch migration domain lengths (11nt, 16nt, 21nt and 26nt) were explored (Figure 4D). A fixed invader toehold of 4nt was selected as our initial model predictions suggested this toehold length would provide the most obvious rate differences between systems with uphill and downhill free-energy landscapes (supplementary note 4 and Figure S1). We found that for DNA_py_>DNA_py_ strand displacement reactions, *k*_eff_ decreased by a factor of 15 for an approximate 2.5-fold increase in branch migration domain length. This fold-change is slightly larger than we would predict by equation (16), although this is likely explained by variability between sequences. More importantly, *k*_eff_ for DNA_py_>RNA_py_ strand displacement only decreased by a factor of 5.7 between 11nt and 26nt, suggesting that *k*_eff_ is less dependent on the length of the branch migration domain as compared to DNA_py_>DNA_py_ strand displacement. Finally, the rate of RNA_py_>DNA_py_ strand displacement decreased by a factor of 54 for an increase in the branch migration domain length between 11nt and 26nt, which is significantly larger than for DNA_py_>DNA_py_ strand displacement.

Using the experimental data for RNA_py_>DNA_py_, DNA_py_>DNA_py_ and DNA_py_>RNA_py_ strand displacement systems we were able to parameterise our adapted IEL model and estimate Δ*G*_rd_. The available kinetic data did not strongly constrain all IEL parameters (Table S16), as such we performed a 3-parameter fit to 24 rate constants. The predicted *k*_eff_ values obtained from the fully parameterised model showed good fit to the experimental kinetic data across both toehold length and branch migration domain length (Figure 4E and 4F). The estimated free-energy parameters and rate parameter are given in Table 1. These parameters are in good agreement with previous values of DNA>DNA strand displacement reactions (10, 11). For the low RNA purine content system we extracted a Δ*G*_rd_ value of 0.16 *k*_B_*T*. This value of Δ*G*_rd_ for RNA_py_>DNA_py_ strand displacement predicts an overall uphill (although notably shallow) free-energy landscape during branch migration. In contrast, our model predicts an overall downhill free-energy landscape for DNA_py_>RNA_py_ branch migration with a Δ*G*_rd_ value of the same magnitude.

Next we probed the rate of strand displacement for an independent design with alternative sequence composition. Specifically, we investigated the effect of high purine content (78%) within the RNA invader strand (Figure 5A). We studied 3 different branch migration domain lengths (11nt, 16nt and 21nt) for a fixed invader toehold length of 4nt. As above, we estimated *k*_eff_ for each system by fitting an ODE model to the fluorescence traces (Figure 5B). In complete contrast to the low RNA purine content design, we found that RNA_pu_>DNA_pu_ strand displacement was up to an order of magnitude faster than DNA_pu_>DNA_pu_ strand displacement, while DNA_pu_>RNA_pu_ strand displacement was over an order of magnitude slower than DNA_pu_>DNA_pu_ strand displacement (Figure 5C). For DNA_pu_>DNA_pu_ strand displacement we see a reduction in *k*_eff_ by a factor of 1.7 between branch migration domain lengths of 11nt and 21nt, in accordance with equation (16). For an approximate doubling in the branch migration domain length, we observe no decrease or even a slight increase in *k*_eff_ for RNA_pu_>DNA_pu_ strand displacement (*k*_eff_ = 9.670 · 10^4^ M^-1^ s^-1^ for *β* = 11nt and *k*_eff_ = 1.640 · 10^5^ M^-1^ s^-1^ for *β* = 21nt), indicating that *k*_eff_ is effectively independent of branch migration domain length across this range. Finally, we observe an reduction in *k*_eff_ by a factor of 4.8 for DNA_pu_>RNA_pu_ strand displacement for an increase in *β* between 11nt and 16nt. Notably, we were unable to explicitly estimate *k*_eff_ for *β* = 21nt because the reaction speed was slow enough for an undesired leak reaction to obfuscate the reaction kinetics of interest. Numerical estimates of *k*_eff_ for RNA_pu_>DNA_pu_, DNA_pu_>DNA_pu_ and DNA_pu_>RNA_pu_ strand displacement reactions are given in Table S13 and Table S15.

**Figure 5.**
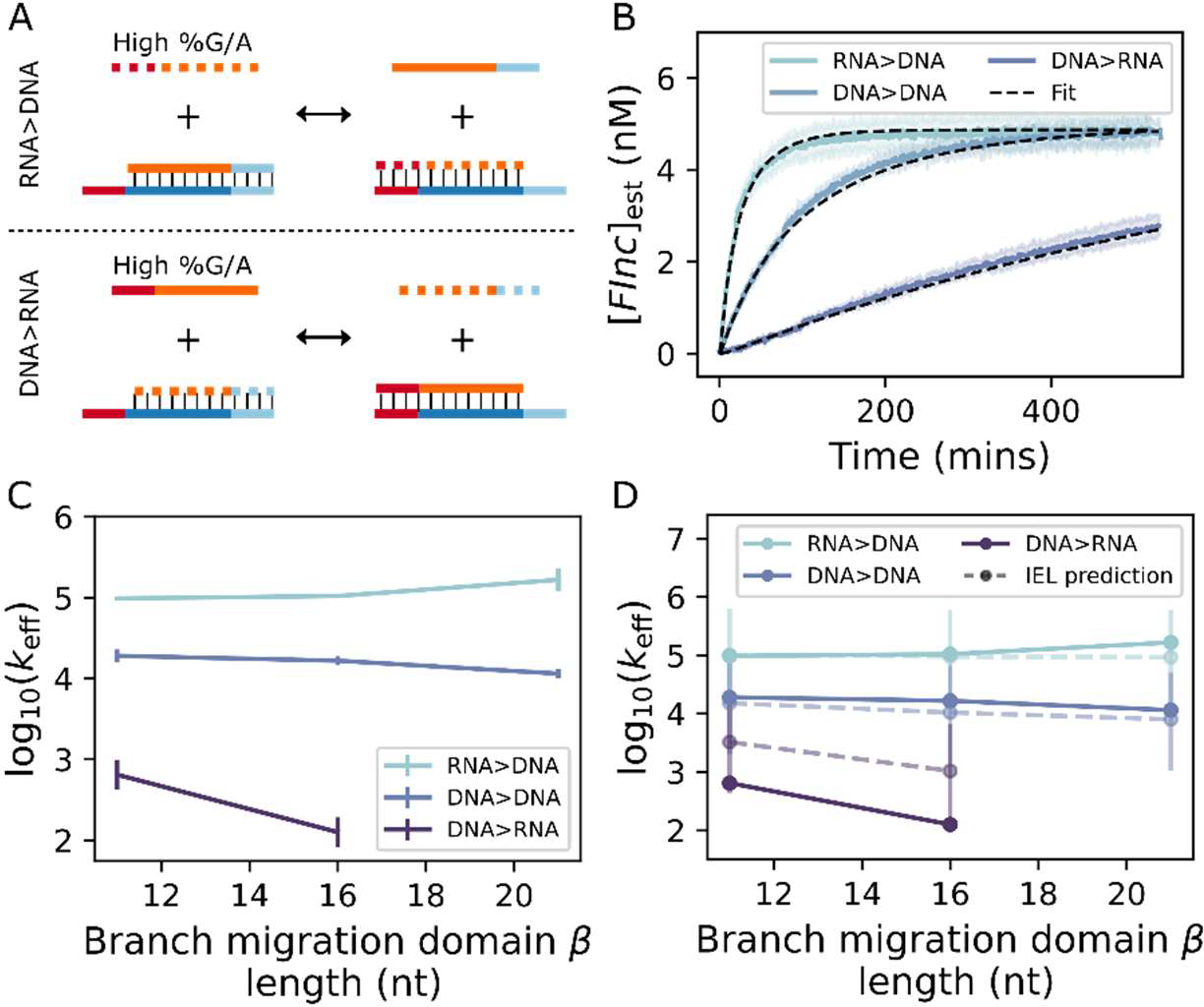
RNA>DNA and DNA>RNA toehold exchange kinetics for high RNA purine content design. A) Schematic of RNA_pu_>DNA_pu_ and DNA_pu_>RNA_pu_ toehold exchange. B) Example fluorescent traces for RNA_pu_>DNA_pu_, DNA_pu_>DNA_pu_, DNA_pu_>RNA_pu_ toehold exchange system with *γ* = 4nt, *β* = 11nt and *ε* = 4nt for an *Inv* concentration of 6 nM (for RNA_pu_>DNA_pu_, DNA_pu_>DNA_pu_) or 60 nM (DNA_pu_>RNA_pu_) across 3 replicates. Solid, coloured lines represent normalised fluorescent curves. Dashed, black lines represent output of fitting an ODE model to this kinetic data. C) Summary of experimentally-derived *k*_eff_ values for RNA_pu_>DNA_pu_, DNA_pu_>DNA_pu_ and DNA_pu_>RNA_pu_ toehold exchange reactions across a range of branch migration domain lengths (11-21nt) for fixed *γ* = 4nt and *ε* = 4nt. For the DNA_pu_>RNA_pu_ toehold exchange reaction with *β* = 21nt, *k*_eff_ could not be reliably estimated due to an undesired leak reaction. Error bars show 95% confidence intervals based on the standard error of *k*_eff_ estimates from individual curves. N=3. D) Predicted *k*_eff_ from parameterised IEL model (dashed lines) compared to experimentally-derived values (solid lines) across branch migration domain lengths (11-21nt). Error bars show 95% confidence intervals of model predicted *k*_eff_ values. N=3.

Using the experimental data from the high purine content design, we reparameterised the IEL model using the same approach as above. We fixed values for Δ*G*_bp_, Δ*G*_assoc_ and Δ*G*_p_ as above. We also fixed Δ*G*_bm_ at the value predicted from experimental rate data of the low RNA purine content design (Table 1). We allowed both *k*_bp_ and Δ*G*_rd_ to be flexible parameters and extracted estimates of 3.0 (*CI*: 1.9; 4.5) · 10^7^ s^-1^ and −0.25 ± 0.07 *k*_B_*T* for these parameters, respectively. This parameterisation provided *k*_eff_ predictions that were in line with the experimental kinetic data for RNA_pu_>DNA_pu_ and DNA_pu_>DNA_pu_ strand displacement (Figure 5D). However, for DNA_pu_>RNA_pu_ strand displacement the model predicts higher *k*_eff_ values, although the trend across branch migration domain length is correctly predicted. Notably, the estimated value of |Δ*G*_rd_| is slightly larger than that predicted from the low RNA purine content experimental data, but in this case Δ*G*_rd_ is negative. We identified alternative fits and Δ*G*_rd_ estimates by fixing different free-energy parameters but importantly Δ*G*_rd_ is estimated to be negative across all parameterisations (Figure S21, Table S17). In contrast to the low RNA purine content design, these estimates of Δ*G*_rd_ predict an overall downhill (although again notably shallow) free-energy landscape during branch migration for RNA_pu_>DNA_pu_ strand displacement. Meanwhile, this model parameterisation predicts an overall uphill free-energy landscape for DNA_pu_>RNA_pu_ branch migration.

## DISCUSSION

In this work we experimentally characterised RNA>DNA and DNA>RNA strand displacement systems, which revealed notable differences in kinetics between DNA>DNA and RNA/DNA hybrid strand displacement systems, with important implications for the rational design of hybrid reaction networks. We have developed a parameterised free-energy landscape model which accurately predicts strand displacement kinetics for the test hybrid systems employed herein, given a fixed branch migration domain purine content. Most importantly, we demonstrate a strong sequence-dependence for the RNA>DNA and DNA>RNA strand displacement reactions that we tested. We highlight the importance of purine content within the RNA strand in determining the relative stability of RNA-DNA hybrids compared to DNA-DNA duplexes. We found that between high (78%) and low (17%) purine content within the RNA strand, estimated Δ*G*_rd_ values shifted from −0.25 *k*_B_*T* to 0.16 *k*_B_*T*. These results support conclusions from previous thermodynamic studies into RNA-DNA hybridisation, which suggest that a high purine content in the RNA strand of an RNA-DNA complex results in increased stability compared to DNA-DNA duplexes while low RNA purine content results in reduced stability of RNA-DNA complexes (36–39).

Analysis of the RNA-DNA hybrid nearest neighbour (NN) parameters derived by Banerjee et al. (2020) predicts Δ*G*_rd_ values of +0.25 *k*_B_*T* for 17% purine content in the RNA strand and −0.49 *k*_B_*T* for 78% purine content in the RNA strand (36). These values are larger than those predicted in this work and would therefore predict more extreme kinetic behaviour for hybrid systems. It is worth noting that only a limited number of data points were used to derive these hybrid NN parameters and as such extreme behaviour is likely not well accounted for. Importantly, these NN parameters indicate that high purine content within the RNA strand should give a larger effect than low purine content in the RNA strand (36), which is also predicted from our experiments. Notably, there is some suggestion that the relative stability of RNA-DNA hybrids as compared to DNA-DNA complexes is also dependent on the G/C content of the displacement domain, particularly for sequences with high G/C content (39). Our study was limited to systems with approximately 50% GC content within the branch migration domain so future work should investigate how these sequence composition factors interact to determine the kinetics of strand displacement reactions.

Although the estimated Δ*G*_rd_ values of −0.25 *k*_B_*T* and 0.16 *k*_B_*T* are relatively small, our results suggest these free-energy differences per base pair can compound over the whole landscape giving orders of magnitude differences in the rate of RNA>DNA strand displacement compared to DNA>RNA strand displacement. Moreover, changing the sequence composition of the RNA from 17% purine content to 78% purine content resulted in a difference in rate of up to 3 orders of magnitude, suggesting that this factor could be a critical parameter in the rational design of RNA/DNA hybrid strand displacement systems and facilitate tuneable reaction kinetics. Indeed, requesting minimal secondary structure within NUPACK typically yields sequence designs excluding G or C nucleotides (41), and therefore by default these sequences exhibit extremes in purine content. As such our work is particularly relevant to synthetic designs within the low secondary structure constraint.

With knowledge of the underlying free-energy landscapes of hybrid strand displacement, other tools, including use of mismatches, could be exploited for further fine-tuned kinetic control (11, 12). Given that the predicted Δ*G*_rd_ values are relatively small, elimination of mismatches within the incumbent-substrate complex could be employed to counter the observed uphill free-energy landscapes. We also suggest that future studies could explicitly investigate the effect of mismatches within hybrid systems to gain a better understanding how this might differ from DNA>DNA strand displacement reactions.

The vital role of RNA in the correct functioning of intracellular processes makes RNA a critical target for both diagnostics and therapeutics. While real clinically relevant RNA sequences may not fall into these purine content extremes, we propose that our work may provide informative limits on the possible behaviours of these systems. Future work should focus on exploring the role of purine content in more detail across multiple sequence designs to confirm the general applicability of the trends revealed herein and confirm the limits of the behaviour of hybrid systems. Additional studies could equally assess sequences with a more average purine content, although secondary structure will likely be a confounding factor in estimating the reaction kinetics. A potentially more informative study would investigate the effect of purine residue distribution within sequences, which may facilitate more fine-tuned regulation of kinetics and could also go some way to inform the mechanism behind the observed sequence-dependent effect.

At present, our free-energy landscape model is limited to a sequence-average parameterisation, ignoring the specific contributions of each nucleotide. Future iterations of the model could potentially look at introducing a free-energy parameter that explicitly captures the effect of purine residues. This parameterisation would require experimental data from systems with differing purine content distributions. This fully-parameterised model would have the capacity to inform and aid the rational design of RNA/DNA hybrid nucleic acid devices with predictable kinetics, by offering general design principles for such systems.

## DATA AVAILABILITY STATEMENT

All raw data, fitting and parameterisation code are freely available at https://doi.org/10.5281/zenodo.10090783

## ACKNOWLEDGEMENTS

The authors would like to acknowledge Dr Rakesh Mukherjee for his assistance in checking the data and code associated with this work.

