## Supplementary Information for "Strong sequence dependence in RNA/DNA hybrid strand displacement kinetics"

**CONTENTS TABLE**

Table S6. Detailed experimental protocol for reporter characterisation reactions to estimate $k_{\mathrm{rep}}$ ... 30

Figure S3. Example of raw fluorescent traces for reporter characterisation experiment ... 32

*Supplementary note 7.2. Normalisation and data-processing of reporter characterisation experiments for $k_{\mathrm{rep}}$ estimation* ... 33

Figure S4. Processed fluorescent traces for all reporter characterisation reactions ... 35

*Supplementary note 7.3. Fitting ordinary differential equations (ODEs) to reporter characterisation experiments for $k_{\mathrm{rep}}$ estimation* ... 36

Table S7. $k_{\mathrm{rep}}$ estimates from averaging over individual reporter characterisation reactions ... 37

Figure S5. Individual fits to fluorescent traces in reporter characterisation reactions ... 38

Table S8. Global $k_{\mathrm{rep}}$ estimates across reporter characterisation reactions ... 40

Figure S6. Global fits to fluorescent traces in reporter characterisation reactions ... 41

**Supplementary note 8. Full strand displacement reactions** ... 42

*Supplementary note 8.1. Experimental set-up for full strand displacement reactions* ... 42

Table S9. Detailed experimental protocol for DNA>DNA and RNA>DNA strand displacement reactions to estimate $k_{\mathrm{eff}}$ ... 43

Figure S7. Example of raw fluorescent traces for DNA>DNA and RNA>DNA strand displacement experiment ... 45

Table S10. Detailed experimental protocol for $DNA_{py}$>$RNA_{py}$ strand displacement reactions to estimate $k_{\mathrm{eff}}$ ... 46

Figure S8. Example of raw fluorescent traces for $DNA_{py}$>$RNA_{py}$ strand displacement experiment ... 49

Table S11. Detailed experimental protocol for $DNA_{pu}$>$RNA_{pu}$ strand displacement reactions to estimate $k_{\mathrm{eff}}$ ... 50

Figure S9. Example of raw fluorescent traces for $DNA_{pu}$>$RNA_{pu}$ strand displacement experiment ... 53

*Supplementary note 8.2. Normalisation and data-processing of-full strand displacement experiments for $k_{\mathrm{eff}}$ estimation* ... 53

Figure S10. Processed fluorescent traces for full DNA>DNA strand displacement reactions for $k_{\mathrm{eff}}$ estimation ... 56

Figure S11. Processed fluorescent traces for full RNA>DNA strand displacement reactions for $k_{\mathrm{eff}}$ estimation ... 58

Figure S12. Processed fluorescent traces for full DNA>RNA strand displacement reactions for $k_{\mathrm{eff}}$ estimation ... 60

*Supplementary note 8.3. Fitting ODEs to full strand displacement experiments for $k_{\mathrm{eff}}$ estimation* ... 61

### Supplementary note 1. State space and free energies of the Intuitive Energy Landscape model

*Supplementary note 1.1. State nomenclature*

We initially define the lengths of the domains involved in strand displacement, with $\gamma$ representing the length of the invader toehold; $\beta$ the length of the branch migration domain and $\varepsilon$ the length of the incumbent toehold. Therefore, $\varepsilon = 0$ for TMSD systems and $\varepsilon > 0$ for toehold exchange systems. The total length of the substrate strand is specified by $S = \gamma + \beta + \varepsilon$. We define the nucleotides of the substrate strand as starting at position $-\gamma + 1$, with the final nucleotide being defined as position $N$ $(\beta + \varepsilon)$. We model toehold exchange as a one-dimensional free-energy landscape with $S + 1$ states, where state $-\gamma$ (Figure 2AI) refers to the system where invader strand is unbound from the incumbent-substrate complex, and state $N$ (state $\beta + \varepsilon$) describes when the invader has fully displaced the incumbent strand from the incumbent-substrate complex, and the incumbent strand has been released. All states between positions $-\gamma$ and $N$ represent a single step in the strand displacement reaction. State $-\gamma + 1$ (Figure 2AII) denotes when the first toehold nucleotide of the invader strand binds to the toehold domain of the substrate strand. Binding of each subsequent invader nucleotide to toehold comprises a subsequent state (up to state 0 (Figure 2AIII)). During branch migration, each state corresponds to each completed branch migration step, in which an invader nucleotide successfully displaces an incumbent nucleotide (up to state $\beta$ (Figure 2AIV)). For $\varepsilon > 0$, each step between state $\beta$ and state $\beta + \varepsilon - 1$, represents the unbinding of a single incumbent nucleotide from the incumbent toehold. The transition from state $\beta + \varepsilon - 1$ (i.e., state $N - 1$ (Figure 2AV)) to state $\beta + \varepsilon$ (i.e., state $N$ (Figure 2AVI)) describes the final strand displacement step in which the final nucleotide of the incumbent strand dissociates from the substrate strand.

*Supplementary note 1.2. Thermodynamic free energy changes for DNA>DNA strand displacement*

We initially define the reference state as a system in which the invader is fully bound to the invader toehold and a perfectly-matched (i.e., mismatch-free) incumbent is fully bound to the branch migration domain and incumbent toehold of the substrate strand. In the case of DNA>DNA strand displacement, the ground free energy refers to DNA-DNA base pairing within the toeholds and DNA-DNA base pairing within the branch migration domain. For RNA>DNA strand displacement the reference state has RNA-DNA base pairing within the invader toehold and DNA-DNA base pairing within the branch migration domain and incumbent toehold. Finally, for DNA>RNA strand displacement the reference state has DNA-DNA base pairing within the invader toehold and RNA-DNA base pairing within the branch migration domain and incumbent toehold. We assign a free energy $G_{\mathrm{R}} = 0\ k_{\mathrm{B}}T$ to this reference state, and all other states are defined in relation to this state.

Breaking of a base pair is associated with a free-energy penalty of $-\Delta G_{\mathrm{bp}} > 0$. Therefore, dissociation of each invader nucleotide from the toehold domain is associated with an increase in the free energy by $-\Delta G_{\mathrm{bp}}$, relative to the reference state. The free-energy change to reach the state $-\gamma + 1$ is

$$G_{-\gamma+1} - G_{\mathrm{R}} = -\Delta G_{\mathrm{bp}} \cdot (\gamma - 1). \quad (S1)$$

For $\varepsilon > 0$, the same free-energy penalty applies for dissociation of incumbent nucleotides from the incumbent toehold. Dissociation of the final nucleotide from the toehold is also associated with a penalty of $-\Delta G_{\mathrm{bp}}$. However, initial binding of the invader strand to the invader toehold is associated with an additional free-energy ($\Delta G_{\mathrm{assoc}}$, defined at a standard concentration of $c_0 = 1$ M) penalty due to a reduction in the translational and orientational freedom upon binding of the toehold, conferring a decrease in entropy. We introduce the term $\ln\left(\frac{c}{c_0}\right)$ in order to adjust $\Delta G_{\mathrm{assoc}}$ for the actual initial concentration of unbound invader strand ($c$). Therefore, the overall free energy change which accompanies dissociation of the final nucleotide from the toehold is given by

$$G_{-\gamma} - G_{-\gamma+1} = -\Delta G_{\mathrm{assoc}} + k_{\mathrm{B}}T \cdot \ln\left(\frac{c}{c_0}\right) - \Delta G_{\mathrm{bp}}, \quad (S2)$$

where $c$ is the initial concentration of unbound invader (in Molar); $k_{\mathrm{B}}$ is the Boltzmann constant; and $T$ is the temperature (in K). The same free energy change applies for dissociation of the final incumbent nucleotide, be it from the incumbent toehold (as in toehold exchange) or the branch migration domain (as in TMSD).

Progression through the branch migration domain involves displacement of incumbent nucleotides by invader nucleotides, thus the number of base pairs formed with the substrate strand is unchanged compared to the reference state. As such, for DNA>DNA strand displacement, there is no net free energy change as branch migration proceeds. Nevertheless, each branch migration step is associated with a thermodynamic barrier which must be overcome ($\Delta G_{\mathrm{bm}}$) for successful replacement of base pairs. The exception to this rule is the initiation of branch migration, associated with an additional free-energy cost $\Delta G_{\mathrm{p}}$ resulting from the presence of two single-stranded overhangs during displacement, compared to the reference state for which only one single-stranded overhang is present. When considering toehold exchange, there is a reduction in free energy by $\Delta G_{\mathrm{p}}$ for states after position $\beta - 1$, as branch migration is completed thus only one single-stranded overhang is present (Figure 2A).

*Supplementary note 1.3. Thermodynamic free energy changes for RNA>DNA and DNA>RNA strand displacement*

To capture the difference in the thermodynamic stability of RNA-DNA hybrids compared to DNA-DNA duplexes, we introduce the parameter $\Delta G_{\mathrm{rd}}$. $\Delta G_{\mathrm{rd}}$ describes the free energy difference between dissociation of a DNA-DNA base pair compared to RNA-DNA base pair dissociation. Each RNA-DNA base pair which is unbound compared to the reference state is associated with a free-energy penalty of $-\Delta G_{\mathrm{bp}} - \Delta G_{\mathrm{rd}}$. Therefore, the free energy change for breaking of RNA-DNA base pairs within the invader toehold domain to reach state $-\gamma + 1$ from the reference state (as in RNA>DNA strand displacement) is defined as

$$G_{-\gamma+1} - G_{\mathrm{R}} = -(\Delta G_{\mathrm{bp}} + \Delta G_{\mathrm{rd}}) \cdot (\gamma - 1). \qquad (S3)$$

By extension, the free energy change for breaking of RNA-DNA base pairs within the incumbent toehold to reach state $N$ – 1 from state $N$ - $\varepsilon$ (as in DNA>RNA strand displacement) is described as

$$G_{N-1} - G_{N-\varepsilon} = -(\Delta G_{\mathrm{bp}} + \Delta G_{\mathrm{rd}}) \cdot (\varepsilon - 1). \qquad (S4)$$

The free energy change which accompanies the dissociation of the final nucleotide from the toehold is also altered by $-\Delta G_{\mathrm{rd}}$, so for RNA>DNA strand displacement $G_{-\gamma} - G_{-\gamma+1}$, and for DNA>RNA strand displacement $G_N - G_{N-1}$, are calculated as

$$G_{-\gamma} - G_{-\gamma+1} \equiv G_N - G_{N-1} = -\Delta G_{\mathrm{assoc}} + k_{\mathrm{B}}T \cdot \ln\left(\frac{c}{c_0}\right) - \Delta G_{\mathrm{bp}} - \Delta G_{\mathrm{rd}}. \qquad (S5)$$

Unlike DNA>DNA branch migration, RNA displacing DNA and DNA displacing RNA are not thermodynamically symmetrical processes. We introduce this thermodynamic asymmetry by adjusting the free energy by $\Delta G_{\mathrm{rd}}$ for each DNA-DNA base pair that is replaced by an RNA-DNA base pair through branch migration, compared to the reference state. Thus, under the condition where RNA-DNA base pairs are more stable than DNA-DNA base pairs ($\Delta G_{\mathrm{rd}} < 0$) there is an overall reduction in the free energy as branch migration progresses towards completion. In contrast, in the limit where RNA-DNA base pairs are less stable than DNA-DNA base pairs ($\Delta G_{\mathrm{rd}} > 0$) there is an increase in the free energy as branch migration progresses towards completion. The free-energy change for the branch migration process is assumed to be

$$G_{\beta} - G_{\mathrm{R}} = \Delta G_{\mathrm{rd}} \cdot \beta. \qquad (S6)$$

### Supplementary note 2. Kinetics of intuitive energy landscape models

*Supplementary note 2.1. Microscopic transition rates*

We assume that the transitions between the states within our system follow a continuous-time Markov process, in which each transition is independent of the previous transition (1). Therefore, from our free-energy landscapes we can define the rate of forward and reverse transitions between any two adjacent states, $n$ and $n + 1$. To comply with the principle of detailed balance, in which there is no net flux between any two adjacent states at equilibrium (2), the rate of the forward ($k_n^+$) and backward ($k_{n+1}^-$) reactions between any two adjacent states must be related to the free energy change of the transition by

$$\frac{k_n^+}{k_{n+1}^-} = e^{-\frac{(G_{n+1}-G_n)}{k_{\mathrm{B}}T}}. \quad (S7)$$

Therefore, we need only specify either the forward or reverse transition rate between two states to determine the other. We assume the rate of all base pairing events, in which a base pair forms at the end of a double-stranded region, is constant ($k_{\mathrm{bp}}$). Thus the rate of base pair dissociation within the invader toehold ($k_n^-$ for $-\gamma + 1 < n \leq 0$) and incumbent toehold ($k_n^+$ for $N - \varepsilon \leq n < N$ - 1), are assumed to be

$$k_n^- \equiv k_n^+ = k_{\mathrm{bp}} \cdot e^{\frac{\Delta G_{\mathrm{bp}}}{k_{\mathrm{B}}T}}. \quad (S8)$$

We also assume that the rate of RNA-DNA base pair formation is $k_{\mathrm{bp}}$. As such, when we consider RNA-DNA base pair dissociation, e.g., RNA dissociating from the DNA invader toehold in RNA>DNA strand displacement ($k_n^-$ for $-\gamma + 1 < n \leq 0$) or RNA dissociating from the DNA incumbent toehold in DNA>RNA strand displacement ($k_n^+$ for $N - \varepsilon \leq n < N$ - 1), we alter this equation accordingly

$$k_n^- \equiv k_n^+ = k_{\mathrm{bp}} \cdot e^{\frac{\Delta G_{\mathrm{bp}} + \Delta G_{\mathrm{rd}}}{k_{\mathrm{B}}T}}. \quad (S9)$$

We assume the rate of dissociation of the first toehold base pair is equivalent to equation (*S8*) (or equation (*S9*) for RNA-DNA base pairs). Therefore, the rate of invader binding ($k_{-\gamma}^+$), can be written as

$$k_{-\gamma}^+ = k_{\mathrm{bp}} \cdot e^{-\frac{\Delta G_{\mathrm{assoc}}}{k_{\mathrm{B}}T}} \cdot \frac{c}{c_0}. \quad (S10)$$

Similarly, we can determine the rate of each branch migration step ($k_{\mathrm{bm}}$) using the free-energy barrier to branch migration

$$k_{\mathrm{bm}} = k_{\mathrm{bp}} \cdot e^{-\frac{\Delta G_{\mathrm{bm}}}{k_{\mathrm{B}}T}}. \quad (S11)$$

This rate applies to both forward and reverse branch migration steps, for DNA>DNA strand displacement. However, when we consider branch migration steps involving RNA-DNA hybrids, the ratio between the forward and the reverse rate is altered such that it is no longer unity. It is the difference in the forward and the reverse rates which reflects the thermodynamic favourability or unfavourability of RNA>DNA branch migration and DNA>RNA branch migration, respectively.

We assume that the rate of a branch migration step involving the unbinding of a DNA-DNA base pair and its replacement by an RNA-DNA base pair is unchanged compared to DNA>DNA branch migration; i.e., $k_{\mathrm{bm}}$ in equation (*S11*). This rate captures a forward branch migration step in RNA>DNA strand displacement and a backward branch migration step in DNA>RNA strand displacement. By contrast, a branch migration step which involves the unbinding of an RNA-DNA base pair (and subsequent replacement by a DNA-DNA base pair) exhibits an altered rate, related to $\Delta G_{\mathrm{rd}}$. Thus, a forward branch migration step for DNA>RNA ($k_i^+$) strand displacement and a backward branch migration step for RNA>DNA strand displacement ($k_j^-$) have the rate

$$k_j^- \equiv k_i^+ = k_{\mathrm{bp}} \cdot e^{-\left(\frac{\Delta G_{\mathrm{bm}} - \Delta G_{\mathrm{rd}}}{k_{\mathrm{B}}T}\right)}. \quad (S12)$$

*Supplementary note 2.2. Modelling spontaneous incumbent dissociation*

We expand our one-dimensional Markov chain model to include spontaneous incumbent dissociation (3). Spontaneous incumbent dissociation represents an alternative or 'shortcut' pathway for completion of the strand displacement process in which the final few base pairs of a duplex break without needing to be displaced (4). We define $k_n^{\mathrm{off}}$ as the rate of the spontaneous incumbent dissociation, at position $n$. The state $n = N - 1$ can transition directly to $n = N$ without a shortcut, by breaking the final incumbent-substrate base pair. This transition, already present in the model, has a rate (for a DNA incumbent) of

$$k_{N-1}^{\mathrm{off}} = k_{\mathrm{bp}} \cdot e^{-\left(\frac{-\Delta G_{\mathrm{bp}}}{k_{\mathrm{B}}T}\right)}. \quad (S13)$$

We include spontaneous incumbent to model all systems described herein. Notably, we do not explicitly model spontaneous incumbent dissociation from either invader or incumbent toeholds but rather only from within the branch migration domain (states 0 through to $\beta - 1$).

We can define this as the transition state (with known rate of dissociation). We assume that spontaneous dissociation from a state $n \neq N - 1$ corresponds to breaking $N - 1 - n$ base pairs to reach a state with only 1 incumbent-substrate base pair, and then breaking the final base pair. We estimate the overall rate for this process ($k_n^{\mathrm{off}}$) using simple transition state theory as

$$k_n^{\mathrm{off}} = k_{N-1}^{\mathrm{off}} \cdot p_{N-1|n}, \quad (S14)$$

$$p_{N-1|n} = e^{-\frac{(G_{N-1|n} - G_n)}{k_{\mathrm{B}}T}}, \quad (S15)$$

where $p_{N-1|n}$ is the approximate probability of finding the system with only 1 incumbent-substrate base pair given the progress of displacement has reached state $n$.

For DNA>DNA and RNA>DNA strand displacement system, we describe the free energy difference between the transition state and the reaction coordinate as follows, given reaching the transition state requires breaking of $N-n-1$ base pairs:

$$G_{N-1|n} - G_n = -(N-n-1) \cdot \Delta G_{\mathrm{bp}}, \tag{S16}$$

$$p_{N-1|n} = e^{-\left(\frac{-(N-n-1)\cdot\Delta G_{\mathrm{bp}}}{k_{\mathrm{B}}T}\right)}, \tag{S17}$$

$$k_n^{\mathrm{off}} = k_{\mathrm{bp}} \cdot e^{-\left(\frac{-\Delta G_{\mathrm{bp}}}{k_{\mathrm{B}}T}\right)} \cdot e^{-\left(\frac{-(N-n-1)\cdot\Delta G_{\mathrm{bp}}}{k_{\mathrm{B}}T}\right)} = k_{\mathrm{bp}} \cdot e^{\frac{(N-n)\cdot\Delta G_{\mathrm{bp}}}{k_{\mathrm{B}}T}}. \tag{S18}$$

For DNA>RNA strand displacement we have to adjust this equation slightly as incumbent dissociation involves breaking of RNA-DNA base pairs

$$k_n^{\mathrm{off}} = k_{\mathrm{bp}} \cdot e^{\frac{(N-n)\cdot(\Delta G_{\mathrm{rd}}+\Delta G_{\mathrm{bp}})}{k_{\mathrm{B}}T}}. \tag{S19}$$

### Supplementary note 3. Analysis of the free-energy landscape model

To determine the second-order rate constant ($k_{\text{eff}}$) for a strand displacement reaction, we calculate the expected first passage time $\langle t \rangle$, defined as the average time for a system to progress from state $-\gamma$ to state $N$. $\langle t \rangle$ is related to the effective rate constant, assuming low reactant concentrations, as described in equation (*15*).

We apply an analytical solution to assess the expected first passage time for a particular free-energy landscape.

*Supplementary note 3.1. Analytical solution to first passage time*

To extract the expected first passage time, we set transmission boundary conditions on our one-dimensional landscape (3). Namely, any system that reaches state $N$ is placed immediately back at state $-\gamma$ to restart the process. Assuming steady state of this modified system, the flux of the system with transmission boundary conditions allows us to calculate the expected first passage time (3). Ignoring spontaneous dissociation, the flux between any two adjacent states ($n$ and $n$ + 1) is calculated as the difference between the probability of being in state $n$ ($p_n$) multiplied by the forward rate constant from state $n$ to $n$ + 1 ($k_n^+$) and the probability of being in state $n$ + 1 multiplied by the backward rate constant out of state $n$ + 1 to $n$ ($k_{n+1}^-$):

$$j_n = k_n^+ \cdot p_n - k_{n+1}^- \cdot p_{n+1}. \qquad \textit{(S20)}$$

In steady state, there is no net flux into or out of any state $n$, hence $j_{n-1} = j_n = j_{-\gamma}$. By rearrangement of equation (*S20*) we derive

$$\frac{p_n}{j_{-\gamma}} = \frac{1}{k_n^+} + \frac{k_{n+1}^-}{k_n^+} \cdot \frac{p_{n+1}}{j_{-\gamma}}. \qquad \textit{(S21)}$$

To determine the overall flux from state $-\gamma$ to state $N$, we calculate $\frac{p_n}{j_{-\gamma}}$ recursively over all states, with the initial condition that $p_N$ = 0, due to the presence of the transmission boundary at state $N$. $j_{-\gamma}$ represents the flux from state $-\gamma$ to state $-\gamma$ + 1, i.e., the total flux into the system. Therefore, the expected first passage time, $\langle t \rangle$, is calculated as the reciprocal of $j_{-\gamma}$

$$\langle t \rangle = \frac{1}{j_{-\gamma}} = \frac{1}{j_{-\gamma}} \sum_{n=-\gamma}^{N} p_n = \sum_{n=-\gamma}^{N} \frac{p_n}{j_{-\gamma}}. \qquad \textit{(S22)}$$

Introduction of spontaneous incumbent dissociation means that the net flux is no longer constant for all states $n$. However, at steady state, the incoming flux still equals the outgoing flux, at any particular state $n$ + 1. The outgoing flux is now equal to the outgoing branch migration step and the spontaneous incumbent dissociation 'shortcut' pathway

$$j_n = j_{n+1} + k_{n+1}^{\text{off}} \cdot p_{n+1}. \qquad \textit{(S23)}$$

Therefore, we adapt the iterative formula accordingly

$$P_N = 0,$$

$$\frac{j_{N-1}}{j_{N-1}} = 1, \qquad \frac{p_{N-1}}{j_{N-1}} = \frac{1}{k_{N-1}^{+}},$$

$$\frac{j_{N-2}}{j_{N-1}} = 1 + k_{N-1}^{\text{off}} \cdot \frac{p_{N-1}}{j_{N-1}}, \qquad \frac{p_{N-2}}{j_{N-1}} = \frac{1}{k_{N-2}^{+}} \cdot \frac{j_{N-2}}{j_{N-1}} + \frac{k_{N-1}^{-}}{k_{N-2}^{+}} \cdot \frac{p_{N-1}}{j_{N-1}},$$

$$\frac{j_{N-3}}{j_{N-1}} = \frac{j_{N-2}}{j_{N-1}} + k_{N-2}^{\text{off}} \cdot \frac{p_{N-2}}{j_{N-1}}, \qquad \cdots$$

$$\cdots \qquad \frac{p_n}{j_{N-1}} = \frac{1}{k_n^{+}} \cdot \frac{j_n}{j_{N-1}} + \frac{k_{n+1}^{-}}{k_n^{+}} \cdot \frac{p_{n+1}}{j_{N-1}}, \tag{S24}$$

$$\frac{j_{n-1}}{j_{N-1}} = \frac{j_n}{j_{N-1}} + k_n^{\text{off}} \cdot \frac{p_n}{j_{N-1}}, \qquad \cdots \tag{S25}$$

The expected first passage time can then be calculated as

$$\langle t \rangle = \frac{1}{j_{-\gamma}} = \frac{1}{j_{-\gamma}} \cdot \frac{j_{N-1}}{j_{N-1}} \sum_{n=-\gamma}^{N} p_n = \frac{j_{N-1}}{j_{-\gamma}} \sum_{n=-\gamma}^{N} \frac{p_n}{j_{N-1}}. \tag{S26}$$

*Supplementary note 3.2. Calculating the second-order limit*

The relationship between the first passage time and the effective rate constant, defined in equation (*15*) is dependent on the system obeying second-order kinetics. Within this limit the time spent in the three-stranded complex is short relative to the time spent in unbound state $-\gamma$. Therefore, calculating the probability of being in state $-\gamma$ over the probability of being in any state $n$ ($p_{\text{unbound}}$) gives the proportion of time spent in the unbound state compared to the three-stranded complex. We assume that a $p_{\text{unbound}}$ < 0.5 violates the second-order limit. Ignoring spontaneous dissociation $p_{\text{unbound}}$ can be calculated from $\frac{p_{-\gamma}}{j_{-\gamma}}$ and $\frac{p_n}{j_{-\gamma}}$, that are generated in equation (*S21*), as

$$p_{\text{unbound}} = \frac{p_{-\gamma}}{\sum_{n=-\gamma}^{N} p_n} = \frac{\left(\frac{p_{-\gamma}}{j_{-\gamma}}\right)}{\sum_{n=-\gamma}^{N} \frac{p_n}{j_{-\gamma}}}. \tag{S27}$$

In the case of the model including spontaneous incumbent dissociation we use $\frac{p_{-\gamma}}{j_{N-1}}$ and $\frac{p_n}{j_{N-1}}$ that are generated recursively from equation (*S24*) to calculate $p_{\text{unbound}}$:

$$p_{\text{unbound}} = \frac{p_{-\gamma}}{\sum_{n=-\gamma}^{N} p_n} = \frac{\left(\frac{p_{-\gamma}}{j_{N-1}}\right)}{\sum_{n=-\gamma}^{N} \frac{p_n}{j_{N-1}}}. \tag{S28}$$

### Supplementary note 4. Initial predictions from free-energy landscape

We performed some initial predictions using free-energy and rate parameters extracted from Irmisch et al. (2020) to gain insights for experimental design (Table 1). We predicted that for RNA>DNA positive values of $\Delta G_{\mathrm{rd}}$ (uphill displacement landscape) resulted in strong dependence on branch migration domain length (Figure S1A). While for DNA>RNA strand displacement negative values of $\Delta G_{\mathrm{rd}}$ (uphill displacement landscape) yielded a strong dependence on branch migration domain length (Figure S1B). We showed that this difference was most evident for $\gamma$ = 4nt.

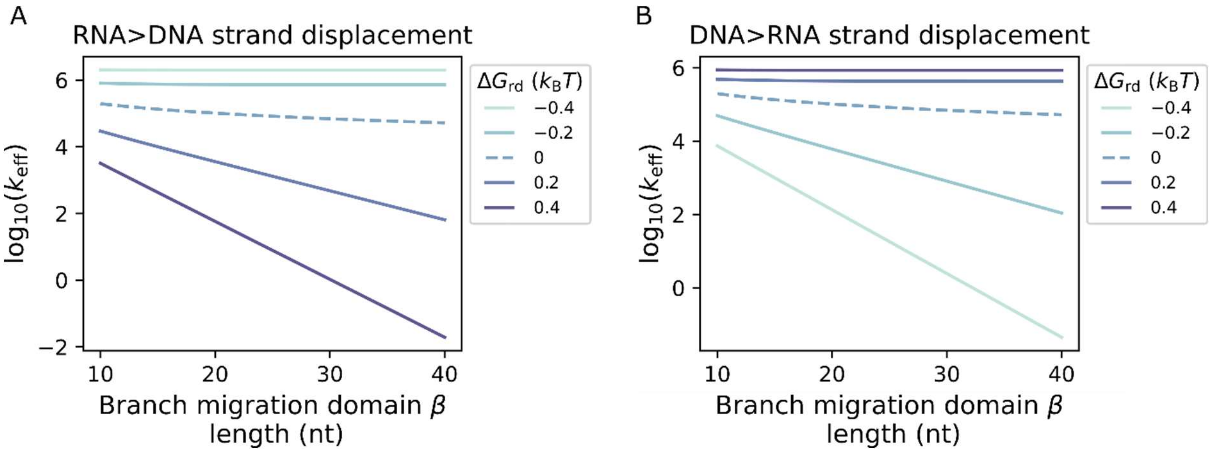

**Figure S1. Initial model predicted $k_{\mathrm{eff}}$ values across branch migration domain length.** A) Predicted $k_{\mathrm{eff}}$ values for RNA>DNA toehold exchange for $\gamma$ = 4nt, $\varepsilon$ = 4nt across a range of branch migration domain lengths ($\beta$ = 10nt – 40nt) for varying $\Delta G_{\mathrm{rd}}$ values ($-0.4\ k_{\mathrm{B}}T$ - $0.4\ k_{\mathrm{B}}T$). B) Predicted $k_{\mathrm{eff}}$ values for DNA>RNA toehold exchange for $\gamma$ = 4nt, $\varepsilon$ = 4nt across a range of branch migration domain lengths ($\beta$ = 10nt to 40nt) for varying $\Delta G_{\mathrm{rd}}$ values ($-0.4\ k_{\mathrm{B}}T$ to $0.4\ k_{\mathrm{B}}T$).

**Supplementary note 5. Sequences**

The strands used in our experiments are listed in Tables S1 and S2. Table S3 lists the particular strands used to generate the results in each figure. We use L and H to indicate strands that were used in low and high RNA purine content experiments, respectively. y, b, d, and z refer to the lengths of $\gamma$, $\beta$, $\delta$ and $\zeta$ domains (or corresponding complementary domain). For *Inc* and *Inv* we add D or R to DNA or RNA versions of each strand, respectively. To refer to specific nucleic acid complexes we combine the strands names as defined in Table S1 and Table S2. *FQ* complexes are given as e.g. d06z16 Rep-FQ, where d06 refers to the toehold domain of *F* and z16 refers to the branch migration domain of both *F* and *Q* strands. Similarly, we define specific *SInc* complexes in the same way e.g. y04b26 SIncD, where y04 refers to the toehold domain of *S* and b26 refers to the branch migration domain of both *S* and *Inc* strands.

| **Strand name** | **Sequence (5′ → 3′)** |
|---|---|
| L-y00b26 IncD | CTC ACC TCA CCT CAC TCC ACC TCC ACT TCA CTC ACC ACT CCT CTC C |
| L-y00b21 IncD | CTC ACC TCA CCT CAC TCC ACC TCC ACT TCA CTC ACC ACT CC |
| L-y00b16 IncD | CTC ACC TCA CCT CAC TCC ACC TCC ACT TCA CTC ACC |
| L-y00b11 IncD | CTC ACC TCA CCT CAC TCC ACC TCC ACT TCA C |
| L-y10b26 Sub | GTT GAG GAG AGG AGA GGA GTG GTG AGT GAA GTG GAG GTG G |
| L-y07b26 Sub | GAG GAG AGG AGA GGA GTG GTG AGT GAA GTG GAG GTG G |
| L-y06b26 Sub | AGG AGA GGA GAG GAG TGG TGA GTG AAG TGG AGG TGG |
| L-y05b26 Sub | GGA GAG GAG AGG AGT GGT GAG TGA AGT GGA GGT GG |
| L-y04b26 Sub | GAG AGG AGA GGA GTG GTG AGT GAA GTG GAG GTG G |
| L-y04b21 Sub | GAG AGG AGT GGT GAG TGA AGT GGA GGT GG |
| L-y04b16 Sub | GAG TGG TGA GTG AAG TGG AGG TGG |
| L-y04b11 Sub | GTG AGT GAA GTG GAG GTG G |
| L-y10b26 InvD | CTC CAC TTC ACT CAC CAC TCC TCT CCT CTC CTC AAC |
| L-d06z16 Rep-F | GGA GGT GGA GTG AGG TGA GGT GAG TTT-AlexaFluor 488 |
| L-d00z16 Rep-Q | IowaBlack FQ-TTT CTC ACC TCA CCT CAC TCC |

**Table S1. Sequences of *Inc*, *S*, *Inv*, *F* and *Q* strands for the low RNA purine content design.** Sequences are given 5′ to 3′. Nucleotides in each strand are coloured to correspond to each domain as in Figure 3A. Sequences are given as DNA, for the equivalent RNA sequences for *Inv* and *Inc* strands replace T with U. For the equivalent RNA sequences names for InvD and IncD sequences become InvR and IncR, respectively.

| Strand name | Sequence (5′ → 3′) |
| --- | --- |
| H-y00b21 IncD | GGA GAG AAG GTG TGG TGC GAG TAG GTA GTG AGA TGA AGG TG<br>GGA GAG AAG GTG TGG TGC GAG TAG GTA GTG AGA TGA AGG TG |
| H-y00b16 IncD | GGA GAG AAG GTG TGG TGC GAG TAG GTA GTG AGA TGA<br>GGA GAG AAG GTG TGG TGC GAG TAG GTA GTG AGA TGA |
| H-y00b11 IncD | GGA GAG AAG GTG TGG TGC GAG TAG GTA GTG A<br>GGA GAG AAG GTG TGG TGC GAG TAG GTA GTG A |
| H-y04b21 Sub | CTC ACA CCT TCA TCT CAC TAC CTA CTC GC<br>CTC ACA CCT TCA TCT CAC TAC CTA CTC GC |
| H-y04b16 Sub | ACC TTC ATC TCA CTA CCT ACT CGC<br>ACC TTC ATC TCA CTA CCT ACT CGC |
| H-y04b11 Sub | CAT CTC ACT ACC TAC TCG C<br>CAT CTC ACT ACC TAC TCG C |
| H-y06b21 InvD | GTA GGT AGT GAG ATG AAG GTG TGA GGG<br>GTA GGT AGT GAG ATG AAG GTG TGA GGG |
| H-d06z16 Rep-F | CTA CTC GCA CCA CAC CTT CTC TCC TTT-AlexaFluor 488 |
| H-d08z16 Rep-F | ACC TAC TCG CAC CAC ACC TTC TCT CCT TT-AlexaFluor 488 |
| H-d00z16 Rep-Q | IowaBlack FQ-TTT GGA GAG AAG GTG TGG TGC |

**Table S2. Sequences of *Inc*, *S*, *Inv*, *F* and *Q* strands for the high RNA purine content design.** Sequences are given 5′ to 3′. Nucleotides in each strand are coloured to correspond to each domain as in Figure 3A. Sequences are given as DNA, for the equivalent RNA sequences for *Inv* and *Inc* strands replace T with U. For the equivalent RNA sequences names for InvD and IncD sequences become InvR and IncR, respectively. For *S*, *Inv* and *Inc* strands domains are shown for reactions with H-d06z16 Rep-F (top) and H-d08z16 Rep-F (bottom), respectively.

| Figure | Subfigure | Strands |
|---|---|---|
| Figure 3 | B | L-y00b26 IncD, L-d06z16 Rep-F, L-d00z16 Rep-Q |
| Figure 4 | B | L-y10b26 InvD, L-y10b26 InvR, L-y05b26 Sub, L-y00b26 IncD, L-y00b26 IncR, L-d06z16 Rep-F, L-d00z16 Rep-Q |
| | C, E | L-y10b26 InvD, L-y10b26 InvR, L-y10b26 Sub, L-y07b26 Sub, L-y06b26 Sub, L-y05b26 Sub, L-y04b26 Sub, L-y00b26 IncD, L-y00b26 IncR, L-d06z16 Rep-F, L-d00z16 Rep-Q |
| | D, F | L-y10b26 InvD, L-y10b26 InvR, L-y04b26 Sub, L-y04b21 Sub, L-y04b16 Sub, L-y04b11 Sub, L-y00b26 IncD, L-y00b26 IncR, L-y00b21 IncD, L-y00b21 IncR, L-y00b16 IncD, L-y00b16 IncR, L-y00b11 IncD, L-y00b11 IncR, L-d06z16 Rep-F, L-d00z16 Rep-Q |
| Figure 5 | B | H-y06b21 InvD, H-y06b21 InvR, H-y04b11 Sub, H-y00b11 IncD, H-y00b11 IncR, H-d06z16 Rep-F, H-d08z16 Rep-F, H-d00z16 Rep-Q |
| | C, D | H-y06b21 InvD, H-y06b21 InvR, H-y04b21 Sub, H-y04b16 Sub, H-y04b11 Sub, H-y00b21 IncD, H-y00b21 IncR, H-y00b16 IncD, H-y00b16 IncR, H-y00b11 IncD, H-y00b11 IncR, H-d06z16 Rep-F, H-d08z16 Rep-F, H-d00z16 Rep-Q |
| Figure S2 | A | L-y00b26 IncD, L-d06z16 Rep-F, L-d00z16 Rep-Q |
| | B | L-y00b26 IncR, L-d06z16 Rep-F, L-d00z16 Rep-Q |
| | C | H-y00b21 IncD, H-d08z16 Rep-F, H-d00z16 Rep-Q |
| | D | H-y00b21 IncR, H-d06z16 Rep-F, H-d00z16 Rep-Q |
| Figure S3 | | L-y00b26 IncR, L_d06z16 Rep-F, L-d00z16 Rep-Q |
| Figure S4 | A | L-y00b26 IncD, L-d06z16 Rep-F, L-d00z16 Rep-Q |
| | B | L-y00b26 IncR, L-d06z16 Rep-F, L-d00z16 Rep-Q |
| | C | H-y00b11 IncD, H-d08z16 Rep-F, H-d00z16 Rep-Q |
| | D | H-y00b11 IncR, H-d06z16 Rep-F, H-d00z16 Rep-Q |
| | E | H-y00b16 IncD, H-d08z16 Rep-F, H-d00z16 Rep-Q |
| | F | H-y00b16 IncR, H-d06z16 Rep-F, H-d00z16 Rep-Q |
| | G | H-y00b21 IncD, H-d08z16 Rep-F, H-d00z16 Rep-Q |
| | H | H-y00b21 IncR, H-d06z16 Rep-F, H-d00z16 Rep-Q |
| Figure S5 | A | L-y00b26 IncD, L-d06z16 Rep-F, L-d00z16 Rep-Q |
| | B | L-y00b26 IncR, L-d06z16 Rep-F, L-d00z16 Rep-Q |
| | C | H-y00b11 IncD, H-d08z16 Rep-F, H-d00z16 Rep-Q |
| | D | H-y00b11 IncR, H-d06z16 Rep-F, H-d00z16 Rep-Q |
| | E | H-y00b16 IncD, H-d08z16 Rep-F, H-d00z16 Rep-Q |
| | F | H-y00b16 IncR, H-d06z16 Rep-F, H-d00z16 Rep-Q |
| | G | H-y00b21 IncD, H-d08z16 Rep-F, H-d00z16 Rep-Q |
| | H | H-y00b21 IncR, H-d06z16 Rep-F, H-d00z16 Rep-Q |
| Figure S6 | A | L-y00b26 IncD, L-d06z16 Rep-F, L-d00z16 Rep-Q |
| | B | L-y00b26 IncR, L-d06z16 Rep-F, L-d00z16 Rep-Q |
| | C | H-y00b11 IncD, H-d08z16 Rep-F, H-d00z16 Rep-Q |
| | D | H-y00b11 IncR, H-d06z16 Rep-F, H-d00z16 Rep-Q |
| | E | H-y00b16 IncD, H-d08z16 Rep-F, H-d00z16 Rep-Q |
| | F | H-y00b16 IncR, H-d06z16 Rep-F, H-d00z16 Rep-Q |
| | G | H-y00b21 IncD, H-d08z16 Rep-F, H-d00z16 Rep-Q |

| | | |
|---|---|---|
| | H | H-y00b21 IncR, H-d06z16 Rep-F, H-d00z16 Rep-Q |
| Figure S7 | | L-y10b26 InvD, L-y04b11 Sub, L-y00b11 IncD, L-d06z16 Rep-F, L-d00z16 Rep-Q |
| Figure S8 | | L-y10b26 InvD, L-y04b11 Sub, L-y00b11 IncR, L-d06z16 Rep-F, L-d00z16 Rep-Q |
| Figure S9 | | H-y06b21 InvD, H-y04b11 Sub, H-y00b11 IncR, H-d06z16 Rep-F, H-d00z16 Rep-Q |
| Figure S10 | A | L-y10b26 InvD, L-y04b11 Sub, L-y00b11 IncD, L-d06z16 Rep-F, L-d00z16 Rep-Q |
| | B | L-y10b26 InvD, L-y04b16 Sub, L-y00b16 IncD, L-d06z16 Rep-F, L-d00z16 Rep-Q |
| | C | L-y10b26 InvD, L-y04b21 Sub, L-y00b21 IncD, L-d06z16 Rep-F, L-d00z16 Rep-Q |
| | D | L-y10b26 InvD, L-y04b26 Sub, L-y00b26 IncD, L-d06z16 Rep-F, L-d00z16 Rep-Q |
| | E | L-y10b26 InvD, L-y05b26 Sub, L-y00b26 IncD, L-d06z16 Rep-F, L-d00z16 Rep-Q |
| | F | L-y10b26 InvD, L-y06b26 Sub, L-y00b26 IncD, L-d06z16 Rep-F, L-d00z16 Rep-Q |
| | G | L-y10b26 InvD, L-y07b26 Sub, L-y07b26 IncD, L-d06z16 Rep-F, L-d00z16 Rep-Q |
| | H | L-y10b26 InvD, L-y10b26 Sub, L-y00b26 IncD, L-d06z16 Rep-F, L-d00z16 Rep-Q |
| | I | H-y06b21 InvD, H-y04b11 Sub, H-y00b11 IncD, H-d08z16 Rep-F, H-d00z16 Rep-Q |
| | J | H-y06b21 InvD, H-y04b16 Sub, H-y00b16 IncD, H-d08z16 Rep-F, H-d00z16 Rep-Q |
| | K | H-y06b21 InvD, H-y04b21 Sub, H-y00b21 IncD, H-d08z16 Rep-F, H-d00z16 Rep-Q |
| Figure S11 | A | L-y10b26 InvR, L-y04b11 Sub, L-y00b11 IncD, L-d06z16 Rep-F, L-d00z16 Rep-Q |
| | B | L-y10b26 InvR, L-y04b16 Sub, L-y00b16 IncD, L-d06z16 Rep-F, L-d00z16 Rep-Q |
| | C | L-y10b26 InvR, L-y04b21 Sub, L-y00b21 IncD, L-d06z16 Rep-F, L-d00z16 Rep-Q |
| | D | L-y10b26 InvR, L-y04b26 Sub, L-y00b26 IncD, L-d06z16 Rep-F, L-d00z16 Rep-Q |
| | E | L-y10b26 InvR, L-y05b26 Sub, L-y00b26 IncD, L-d06z16 Rep-F, L-d00z16 Rep-Q |
| | F | L-y10b26 InvR, L-y06b26 Sub, L-y00b26 IncD, L-d06z16 Rep-F, L-d00z16 Rep-Q |
| | G | L-y10b26 InvR, L-y07b26 Sub, L-y07b26 IncD, L-d06z16 Rep-F, L-d00z16 Rep-Q |
| | H | L-y10b26 InvR, L-y10b26 Sub, L-y00b26 IncD, L-d06z16 Rep-F, L-d00z16 Rep-Q |

| | | |
|---|---|---|
| | I | H-y06b21 InvR, H-y04b11 Sub, H-y00b11 IncD, H-d08z16 Rep-F, H-d00z16 Rep-Q |
| | J | H-y06b21 InvR, H-y04b16 Sub, H-y00b16 IncD, H-d08z16 Rep-F, H-d00z16 Rep-Q |
| | K | H-y06b21 InvR, H-y04b21 Sub, H-y00b21 IncD, H-d08z16 Rep-F, H-d00z16 Rep-Q |
| Figure S12 | A | L-y10b26 InvD, L-y04b11 Sub, L-y00b11 IncR, L-d06z16 Rep-F, L-d00z16 Rep-Q |
| | B | L-y10b26 InvD, L-y04b16 Sub, L-y00b16 IncR, L-d06z16 Rep-F, L-d00z16 Rep-Q |
| | C | L-y10b26 InvD, L-y04b21 Sub, L-y00b21 IncR, L-d06z16 Rep-F, L-d00z16 Rep-Q |
| | D | L-y10b26 InvD, L-y04b26 Sub, L-y00b26 IncR, L-d06z16 Rep-F, L-d00z16 Rep-Q |
| | E | L-y10b26 InvD, L-y05b26 Sub, L-y00b26 IncR, L-d06z16 Rep-F, L-d00z16 Rep-Q |
| | F | L-y10b26 InvD, L-y06b26 Sub, L-y00b26 IncR, L-d06z16 Rep-F, L-d00z16 Rep-Q |
| | G | L-y10b26 InvD, L-y07b26 Sub, L-y07b26 IncR, L-d06z16 Rep-F, L-d00z16 Rep-Q |
| | H | L-y10b26 InvD, L-y10b26 Sub, L-y00b26 IncR, L-d06z16 Rep-F, L-d00z16 Rep-Q |
| | I | H-y06b21 InvD, H-y04b11 Sub, H-y00b11 IncR, H-d06z16 Rep-F, H-d00z16 Rep-Q |
| | J | H-y06b21 InvD, H-y04b16 Sub, H-y00b16 IncR, H-d06z16 Rep-F, H-d00z16 Rep-Q |
| | K | H-y06b21 InvD, H-y04b21 Sub, H-y00b21 IncR, H-d06z16 Rep-F, H-d00z16 Rep-Q |
| Figure S13 | A | L-y10b26 InvD, L-y04b11 Sub, L-y00b11 IncD, L-d06z16 Rep-F, L-d00z16 Rep-Q |
| | B | L-y10b26 InvD, L-y04b16 Sub, L-y00b16 IncD, L-d06z16 Rep-F, L-d00z16 Rep-Q |
| | C | L-y10b26 InvD, L-y04b21 Sub, L-y00b21 IncD, L-d06z16 Rep-F, L-d00z16 Rep-Q |
| | D | L-y10b26 InvD, L-y04b26 Sub, L-y00b26 IncD, L-d06z16 Rep-F, L-d00z16 Rep-Q |
| | E | L-y10b26 InvD, L-y05b26 Sub, L-y00b26 IncD, L-d06z16 Rep-F, L-d00z16 Rep-Q |
| | F | L-y10b26 InvD, L-y06b26 Sub, L-y00b26 IncD, L-d06z16 Rep-F, L-d00z16 Rep-Q |
| | G | L-y10b26 InvD, L-y07b26 Sub, L-y00b26 IncD, L-d06z16 Rep-F, L-d00z16 Rep-Q |
| | H | L-y10b26 InvD, L-y10b26 Sub, L-y00b26 IncD, L-d06z16 Rep-F, L-d00z16 Rep-Q |
| | I | H-y06b21 InvD, H-y04b11 Sub, H-y00b11 IncD, H-d08z16 Rep-F, H-d00z16 Rep-Q |

| | | |
|---|---|---|
| | J | H-y06b21 InvD, H-y04b16 Sub, H-y00b16 IncD, H-d08z16 Rep-F, H-d00z16 Rep-Q |
| | K | H-y06b21 InvD, H-y04b21 Sub, H-y00b21 IncD, H-d08z16 Rep-F, H-d00z16 Rep-Q |
| Figure S14 | A | L-y10b26 InvR, L-y04b11 Sub, L-y00b11 IncD, L-d06z16 Rep-F, L-d00z16 Rep-Q |
| | B | L-y10b26 InvR, L-y04b16 Sub, L-y00b16 IncD, L-d06z16 Rep-F, L-d00z16 Rep-Q |
| | C | L-y10b26 InvR, L-y04b21 Sub, L-y00b21 IncD, L-d06z16 Rep-F, L-d00z16 Rep-Q |
| | D | L-y10b26 InvR, L-y04b26 Sub, L-y00b26 IncD, L-d06z16 Rep-F, L-d00z16 Rep-Q |
| | E | L-y10b26 InvR, L-y05b26 Sub, L-y00b26 IncD, L-d06z16 Rep-F, L-d00z16 Rep-Q |
| | F | L-y10b26 InvR, L-y06b26 Sub, L-y00b26 IncD, L-d06z16 Rep-F, L-d00z16 Rep-Q |
| | G | L-y10b26 InvR, L-y07b26 Sub, L-y00b26 IncD, L-d06z16 Rep-F, L-d00z16 Rep-Q |
| | H | L-y10b26 InvR, L-y10b26 Sub, L-y00b26 IncD, L-d06z16 Rep-F, L-d00z16 Rep-Q |
| | I | H-y06b21 InvR, H-y04b11 Sub, H-y00b11 IncD, H-d08z16 Rep-F, H-d00z16 Rep-Q |
| | J | H-y06b21 InvR, H-y04b16 Sub, H-y00b16 IncD, H-d08z16 Rep-F, H-d00z16 Rep-Q |
| | K | H-y06b21 InvR, H-y04b21 Sub, H-y00b21 IncD, H-d08z16 Rep-F, H-d00z16 Rep-Q |
| Figure S15 | A | L-y10b26 InvD, L-y04b11 Sub, L-y00b11 IncR, L-d06z16 Rep-F, L-d00z16 Rep-Q |
| | B | L-y10b26 InvD, L-y04b16 Sub, L-y00b16 IncR, L-d06z16 Rep-F, L-d00z16 Rep-Q |
| | C | L-y10b26 InvD, L-y04b21 Sub, L-y00b21 IncR, L-d06z16 Rep-F, L-d00z16 Rep-Q |
| | D | L-y10b26 InvD, L-y04b26 Sub, L-y00b26 IncR, L-d06z16 Rep-F, L-d00z16 Rep-Q |
| | E | L-y10b26 InvD, L-y05b26 Sub, L-y00b26 IncR, L-d06z16 Rep-F, L-d00z16 Rep-Q |
| | F | L-y10b26 InvD, L-y06b26 Sub, L-y00b26 IncR, L-d06z16 Rep-F, L-d00z16 Rep-Q |
| | G | L-y10b26 InvD, L-y07b26 Sub, L-y00b26 IncR, L-d06z16 Rep-F, L-d00z16 Rep-Q |
| | H | L-y10b26 InvD, L-y10b26 Sub, L-y00b26 IncR, L-d06z16 Rep-F, L-d00z16 Rep-Q |
| Figure S16 | A | L-y10b26 InvD, L-y04b11 Sub, L-y00b11 IncD, L-d06z16 Rep-F, L-d00z16 Rep-Q |
| | B | L-y10b26 InvD, L-y04b16 Sub, L-y00b16 IncD, L-d06z16 Rep-F, L-d00z16 Rep-Q |

| | | |
|---|---|---|
| | C | L-y10b26 InvD, L-y04b21 Sub, L-y00b21 IncD, L-d06z16 Rep-F, L-d00z16 Rep-Q |
| | D | L-y10b26 InvD, L-y04b26 Sub, L-y00b26 IncD, L-d06z16 Rep-F, L-d00z16 Rep-Q |
| | E | L-y10b26 InvD, L-y05b26 Sub, L-y00b26 IncD, L-d06z16 Rep-F, L-d00z16 Rep-Q |
| | F | L-y10b26 InvD, L-y06b26 Sub, L-y00b26 IncD, L-d06z16 Rep-F, L-d00z16 Rep-Q |
| | G | L-y10b26 InvD, L-y07b26 Sub, L-y00b26 IncD, L-d06z16 Rep-F, L-d00z16 Rep-Q |
| | H | L-y10b26 InvD, L-y10b26 Sub, L-y00b26 IncD, L-d06z16 Rep-F, L-d00z16 Rep-Q |
| | I | H-y06b21 InvD, H-y04b11 Sub, H-y00b11 IncD, H-d08z16 Rep-F, H-d00z16 Rep-Q |
| | J | H-y06b21 InvD, H-y04b16 Sub, H-y00b16 IncD, H-d08z16 Rep-F, H-d00z16 Rep-Q |
| | K | H-y06b21 InvD, H-y04b21 Sub, H-y00b21 IncD, H-d08z16 Rep-F, H-d00z16 Rep-Q |
| Figure S17 | A | L-y10b26 InvR, L-y04b11 Sub, L-y00b11 IncD, L-d06z16 Rep-F, L-d00z16 Rep-Q |
| | B | L-y10b26 InvR, L-y04b16 Sub, L-y00b16 IncD, L-d06z16 Rep-F, L-d00z16 Rep-Q |
| | C | L-y10b26 InvR, L-y04b21 Sub, L-y00b21 IncD, L-d06z16 Rep-F, L-d00z16 Rep-Q |
| | D | L-y10b26 InvR, L-y04b26 Sub, L-y00b26 IncD, L-d06z16 Rep-F, L-d00z16 Rep-Q |
| | E | L-y10b26 InvR, L-y05b26 Sub, L-y00b26 IncD, L-d06z16 Rep-F, L-d00z16 Rep-Q |
| | F | L-y10b26 InvR, L-y06b26 Sub, L-y00b26 IncD, L-d06z16 Rep-F, L-d00z16 Rep-Q |
| | G | L-y10b26 InvR, L-y07b26 Sub, L-y00b26 IncD, L-d06z16 Rep-F, L-d00z16 Rep-Q |
| | H | L-y10b26 InvR, L-y10b26 Sub, L-y00b26 IncD, L-d06z16 Rep-F, L-d00z16 Rep-Q |
| | I | H-y06b21 InvR, H-y04b11 Sub, H-y00b11 IncD, H-d08z16 Rep-F, H-d00z16 Rep-Q |
| | J | H-y06b21 InvR, H-y04b16 Sub, H-y00b16 IncD, H-d08z16 Rep-F, H-d00z16 Rep-Q |
| | K | H-y06b21 InvR, H-y04b21 Sub, H-y00b21 IncD, H-d08z16 Rep-F, H-d00z16 Rep-Q |
| Figure S18 | A | L-y10b26 InvD, L-y04b11 Sub, L-y00b11 IncR, L-d06z16 Rep-F, L-d00z16 Rep-Q |
| | B | L-y10b26 InvD, L-y04b16 Sub, L-y00b16 IncR, L-d06z16 Rep-F, L-d00z16 Rep-Q |
| | C | L-y10b26 InvD, L-y04b21 Sub, L-y00b21 IncR, L-d06z16 Rep-F, L-d00z16 Rep-Q |

| | | |
|---|---|---|
| | D | L-y10b26 InvD, L-y04b26 Sub, L-y00b26 IncR, L-d06z16 Rep-F, L-d00z16 Rep-Q |
| | E | L-y10b26 InvD, L-y05b26 Sub, L-y00b26 IncR, L-d06z16 Rep-F, L-d00z16 Rep-Q |
| | F | L-y10b26 InvD, L-y06b26 Sub, L-y00b26 IncR, L-d06z16 Rep-F, L-d00z16 Rep-Q |
| | G | L-y10b26 InvD, L-y07b26 Sub, L-y00b26 IncR, L-d06z16 Rep-F, L-d00z16 Rep-Q |
| | H | L-y10b26 InvD, L-y10b26 Sub, L-y00b26 IncR, L-d06z16 Rep-F, L-d00z16 Rep-Q |
| Figure S19 | A | H-y00b11 IncR, H-y04b11 Sub, H-d06z16 Rep-F, H-d00z16 Rep-Q |
| | B | H-y00b16 IncR, H-y04b16 Sub, H-d06z16 Rep-F, H-d00z16 Rep-Q |
| | C | H-y00b21 IncR, H-y04b21 Sub, H-d06z16 Rep-F, H-d00z16 Rep-Q |
| Figure S20 | A | H-y00b11 IncR, H-y04b11 Sub, H-d06z16 Rep-F, H-d00z16 Rep-Q |
| | B | H-y00b16 IncR, H-y04b16 Sub, H-d06z16 Rep-F, H-d00z16 Rep-Q |
| | C | H-y00b21 IncR, H-y04b21 Sub, H-d06z16 Rep-F, H-d00z16 Rep-Q |
| Figure S21 | A, B, C | H-y06b21 InvD, H-y06b21 InvR, H-y04b21 Sub, H-y04b16 Sub, H-y04b11 Sub, H-y00b21 IncD, H-y00b21 IncR, H-y00b16 IncD, H-y00b16 IncR, H-y00b11 IncD, H-y00b11 IncR, H-d06z16 Rep-F, H-d08z16 Rep-F, H-d00z16 Rep-Q |

**Table S3. Strands required to produce figures.** Combinations of strands involved in reactions to generate each figure in both the main text and supplementary information.

**Supplementary note 6. Fluorescence calibration curves**

*Supplementary note 6.1. Experimental set-up for reporter characterisations to generate calibration curves*

All experiments were performed at 1 M NaCl in 1X TAE buffer. All strands were ordered at 100 µM in LabReady format in 1X IDTE (pH = 8.0) from IDT. All experiments were performed using nuclease-free buffers and reagents to minimise unwanted degradation of strands. We diluted all strands in 1X IDTE buffer to approximately 1 µM stock concentration. For all strands the concentration of this dilution was estimated by taking the average of 3 readings on a Nanodrop 1000c and the derived concentration estimate was used for all further calculations. We define this derived concentration of a DNA or RNA strand as the expected concentration e.g. $[x]_{exp}$. The concentration of each strand as indicated by IDT and from nanodrop measurements were not entirely trustworthy and did not allow for good repeatability between experiments. We therefore assume that the concentration of the reporter complex (*FQ*) in the calibration experiments is correct and we use this assumption to guide estimation of concentrations for all other strands across all experiments by fitting to the fluorescence observed. We use $[x]_{est}$ to refer to concentration of strand *x* as estimated based on the calibration experiments.

We generated a number of calibration curves to correlate the fluorescence readings to the concentration of fluorescent product, [*FInc*]. These calibration curves allowed for consistent conversion of fluorescence readings to concentration of *FInc* in nM for all measurements with the same gain and focal height settings. We selected a focal height of 5.9 mm across all experiments as this provided the optimal detection of fluorescence for the reaction volumes used in this work. Moreover, this focal height resulted in minimal differences in fluorescence for dilution of reaction volumes between 150 µL and 220 µL. Consequently, the fluorescent readings actually relate to the molecular count of each species in each well over time. For convenience, however, we refer to these inferred molecular counts in terms of their putative concentration at 200 µL (the volume at which reaction kinetics were measured).

To generate the calibration curves, 100 µM stocks of H-d06z16 Rep-F, H-d08z16 Rep-F and H-d00z16 Rep-Q for the high RNA purine content design and L-d06z16 Rep-F and L-d00z16 Rep-Q for the low RNA purine content design were diluted in 1X IDTE (pH = 8.0) to make two 1 µM stocks. *Inc* strands used to generate the calibration curves (L-y00b26 IncD, L-y00b26 IncR, H-y00b21 IncD and H-y00b21 IncR) were each also diluted into two stocks of approximately 1 µM. L-d06z16 Rep-FQ was combined with L-y00b26 IncD or L-y00b26 IncR to generate fluorescence calibration curves for the low RNA purine content design. H-d08z16 Rep-FQ and H-d06z16 Rep-FQ were combined with H-y00b21 IncD and H-y00b21 IncR, respectively, to generate fluorescence calibration curves for the high RNA purine content design. Each calibration curve was generated from either 3 or 4 replicates of a reporter characterisation reaction. We ensured that a unique combination of the two *FQ* and *Inc* 1 µM stocks were used for each replicate to account for any pipetting error within the calibration curve.

We added 10 µL of 300 nM annealed *FQ* to 140 µL of 1 M NaCl + 1X TAE for a final concentration of 15 nM in 200 µL. For the low RNA purine content design we made four 500 µL stocks with expected concentrations of 100 nM L-y00b26 IncD or L-y00b26 IncR. The 100 nM *Inc* strand stock was injected automatically using automatic injectors of the BMG CLARIOstar® microplate reader. For each replicate we injected a different 100 nM *Inc* stock with varying volumes of 1 M NaCl + 1X TAE in 50 µL. Injection of the 100 nM *Inc* stock results in final, expected incumbent strand concentrations ($[Inc]_{exp}$) of 0 nM, 2 nM, 4 nM, 6 nM, 8 nM, 10 nM, 12 nM, 14 nM and 20 nM in 200 µL (Table S4). For these experiments a total of 4 replicates were performed. For the high purine content design we made four 500 µL stocks of 200 nM H-y00b21 IncD or H-y00b21 IncR. In this case we injected the 200 nM *Inc* stock with varying volumes of 1 M NaCl + 1X TAE in 50 µL for final, expected incumbent strand concentrations ($[Inc]_{exp}$) of 0 nM, 4 nM, 8 nM, 12 nM, 16 nM, 20 nM, 22 nM, 24 nM and 30 nM (Table S5). For these experiments a total of 3 (for H-y00b21 IncD) or 4 replicates were performed.

Finally, for both high and low RNA purine content experiments when the reaction kinetics had reached completion, we added 10 µL of 400 nM of the corresponding *Inc* for an expected, additional concentration of 20 nM.

For each replicate we included a positive control with expected *F* concentration of 15 nM and an excess (20 nM) of *Inc* in order to form the final fluorescent product *FInc* in 1 M NaCl + 1X TAE. The positive control allowed for corrections in fluorescence as a result of fluctuations in temperature and reaction volume. We had a buffer-only negative control with 1 M NaCl + 1X TAE. We also had a second negative control with an expected *FQ* concentration of 15 nM and to which 0 nM *Inc* was injected. The volume of the controls was adjusted in line with the main assay at each step by injecting corresponding volumes of 1 M NaCl + 1X TAE.

For all reporter characterisation reactions we captured the reaction kinetics and estimated the initial concentration of reactants with a total of 4 individual measurements. Note that these experiments were also used to determine the kinetics of reporter reactions, explaining the multi-step procedure.

<u>Measurement 1:</u> Measurement 1 captured the baseline for the quenched reporter (*FQ*) fluorescence.

<u>Measurement 2:</u> Measurement 2 monitored the reaction kinetics following injection of *Inc*. Most reporter reactions we measured exhibited rapid kinetics and as such we exploited the 'well mode' of the BMG CLARIOstar® microplate reader which reads each well individually for the entire duration of the measurement before progressing to the next well. In this way 'well mode' minimises the time between reads. For the calibration reporter reactions with slower kinetics (H-y00b21 IncD and H-y00b21 IncR) we employed the 'plate mode' which cycles through all wells for a single read before returning to the first well and repeating for each read.

<u>Measurement 3:</u> Measurement 3 quantified the steady state fluorescence when all *Inc* strands had reacted with *FQ*, facilitating estimation of $[Inc]$ at time t = 0 s ($[Inc]_0$).

Measurement 4: Measurement 4 monitored fluorescence after an excess of *Inc* strand has been injected to trigger dissociation of all *FQ* complexes and determine the maximum unquenched fluorescence.

| Steps | Total reaction volume (μL) | Addition | Expected concentration in 200 μL (nM) | Purpose |
|---|---|---|---|---|
| Measurement 1 | 150 | Reaction: 140 μL of 1 M NaCl + 1X TAE + 10 μL $[FQ]_{exp}$ = 300 nM | $[FQ]_{exp}$ = 15 | Estimate residual fluorescence baseline |
| | | Positive control: 3 μL of $[F]_{exp}$ = 1 μM + 10 μL of $[Inc]_{exp}$ = 400 nM + 137 μL of 1 M NaCl + 1X TAE | $[FInc]_{PCexp}$ = 15 | Reference fluorescence value |
| | | Negative control: 140 μL of 1 M NaCl + 1X TAE + 10 μL $[FQ]_{exp}$ = 300 nM | $[FQ]_{NCexp}$ = 15 | |
| | | Buffer-only control: 150 μL of 1 M NaCl + 1X TAE | | |
| Measurement 2 | 200 | Reaction: 4-40 μL of $[Inc]_{exp}$ = 100 nM + 46-10 μL of 1 M NaCl + 1X TAE | Evolve with time | |
| | | Positive control: 50 μL of 1 M NaCl + 1X TAE | $[FInc]_{PCexp}$ = 15 | |
| | | Negative control: 50 μL of 1 M NaCl + 1X TAE | $[FQ]_{NCexp}$ = 15 | |
| | | Buffer-only control: | | |

| | | | | |
|---|---|---|---|---|
| | | 50 µL of 1 M NaCl + 1X TAE | | |
| Measurement 3 | 200 | Reaction: - | $[FQ]_{exp}$ = 13-0; $[FInc]_{exp}$ = 2-15 | Estimate $[Inc]_0$ |
| | | Positive control: - | $[FInc]_{PCexp}$ = 15 | |
| | | Negative control: - | $[FQ]_{NCexp}$ = 15 | |
| | | Buffer-only control: - | | |
| Measurement 4 | 210 | 10 µL of $[Inc]_{exp}$ = 400 nM | $[FInc]_{exp}$ = 15 | |
| | | Positive control: 10 µL of 1 M NaCl + 1X TAE | $[FInc]_{PCexp}$ = 15 | |
| | | Negative control: 10 µL of $[Inc]_{exp}$ = 400 nM | $[FInc]_{NCexp}$ = 15 | |
| | | Buffer-only control: 10 µL of 1 M NaCl + 1X TAE | | |

**Table S4. Detailed experimental protocol for reporter characterisation reactions to generate calibration curves for the low RNA purine content design.** Summary of volumes of reactants added at each measurement for reporter characterisation to generate calibration curves (afu to nM) for the low RNA purine content design. *FQ* is added at an expected concentration of 15 nM for all reactions and a series of different expected *Inc* concentrations are added to trigger initiation of the reaction. The expected concentration of reactants and products in 200 µL are given for each measurement. The purpose of each measurement for the further normalisation and post-processing are outlined.

| Steps | Total reaction volume (μL) | Addition | Expected concentration in 200 μL (nM) | Purpose |
|---|---|---|---|---|
| Measurement 1 | 150 | Reaction: 140 μL of 1 M NaCl + 1X TAE + 10 μL $[FQ]_{exp}$ = 300 nM | $[FQ]_{exp}$ = 15 | Estimate residual fluorescence baseline |
| | | Positive control: 3 μL of $[F]_{exp}$ = 1 μM + 10 μL of $[Inc]_{exp}$ = 400 nM + 137 μL of 1 M NaCl + 1X TAE | $[FInc]_{PCexp}$ = 15 | Reference fluorescence value |
| | | Negative control: 140 μL of 1 M NaCl + 1X TAE + 10 μL $[FQ]_{exp}$ = 300 nM | $[FQ]_{NCexp}$ = 15 | |
| | | Buffer-only control: 150 μL of 1 M NaCl + 1X TAE | | |
| Measurement 2 | 200 | Reaction: 4-30 μL of $[Inc]_{exp}$ = 200 nM + 46-20 μL of 1 M NaCl + 1X TAE | Evolve with time | |
| | | Positive control: 50 μL of 1 M NaCl + 1X TAE | $[FInc]_{PCexp}$ = 15 | |
| | | Negative control: 50 μL of 1 M NaCl + 1X TAE | $[FQ]_{NCexp}$ = 15 | |
| | | Buffer-only control: 50 μL of 1 M NaCl + 1X TAE | | |
| Measurement 3 | 200 | Reaction: | $[FQ]_{exp}$ = 11-0; | Estimate $[Inc]_0$ |

| | | | |
|---|---|---|---|
| | | - | $[FInc]_{exp}$ = 4-15 |
| | | Positive control: - | $[FInc]_{PCexp}$ = 15 |
| | | Negative control: - | $[FQ]_{NCexp}$ = 15 |
| | | Buffer-only control: - | |
| Measurement 4 | 210 | Reaction: 10 µL of $[Inc]_{exp}$ = 400 nM | $[FInc]_{exp}$ = 15 |
| | | Positive control: 10 µL of 1 M NaCl + 1X TAE | $[FInc]_{PCexp}$ = 15 |
| | | Negative control: 10 µL of $[Inc]_{exp}$ = 400 nM | $[FInc]_{NCexp}$ = 15 |
| | | Buffer-only control: 10 µL of 1 M NaCl + 1X TAE | |

**Table S5. Detailed experimental protocol for reporter characterisation reactions to generate calibration curves for the high RNA purine content design.** Summary of volumes of reactants added at each measurement for reporter characterisation to generate calibration curves (afu to nM) for the high RNA purine content design. *FQ* is added at an expected concentration of 15 nM for all reactions and a series of different expected *Inc* concentrations are added to trigger initiation of the reaction. The expected concentration of reactants and products in 200 µL are given for each measurement. The purpose of each measurement for the further normalisation and post-processing are outlined.

*Supplementary note 6.2. Normalisation and data-processing to generate calibration curves for fluorescence scaling*

<u>Step 1: Removing background fluorescence produced by experimental buffer.</u>

The first step in processing all data is to remove the background autofluorescence resulting from the 1 M NaCl + 1X TAE buffer. At each time point the fluorescence of the buffer-only negative control was subtracted from the raw fluorescence values for all other wells (experimental reactions, positive and negative controls).

<u>Step 2: Correction based on the positive control fluorescence</u>

Fluctuations in the fluorescence can occur without a change in the yield of fluorescent species. We use the change in the positive control fluorescence to correct for these fluctuations. We define the mean fluorescence during measurement 1 as the reference fluorescence value to which all other fluorescence is compared. We can assume that during the first measurement any photobleaching is negligible and all wells should be at the experimental temperature because plates were left for 30 minutes at the experimental temperature prior to the first measurement. For reporter reactions with slower kinetics we calculate the ratio between the reference fluorescence value and the fluorescence of the positive control at each time point. For all wells the measured fluorescence is multiplied by this ratio at each time point in order to correct for any changes in the fluorescence of the positive control.

For systems with rapid kinetics, for which the 'well mode' was employed, we took a slightly different approach for this normalisation step. Well mode appeared to be accompanied with greater noise in the fluorescence of the positive control as well as signal artefacts such as jumps and drops in fluorescence. In order to avoid carrying these artefacts through to all wells instead we normalised by multiplying all data points by the ratio between the reference fluorescence value and the mean of the positive control fluorescence at steady state (measurement 3).

*Supplementary note 6.3. Linear regression for fluorescence calibration curves*

After performing steps 1 and 2 on this reporter characterisation data, we next generated a calibration curve to convert fluorescence in arbitrary fluorescence units measured by the microplate reader (afu) to [*Inc*] in nM. For each replicate and each initial incumbent concentration, we took a mean of the fluorescence at steady state (measurement 3) and plotted this value against the expected concentration of *Inc* in nM.

For all calibration curves we included at least one *Inc* at an expected concentration higher than the expected concentration of *FQ* (15 nM). However, for the purpose of fitting a calibration curve, we only included data points for which the fluorescence was still increasing linearly and had not yet started to plateau, suggesting that [*Inc*] was equal to or greater than [*FQ*]. For L-y00b26 IncD and L-y00b26 IncR with d06z16 Rep-FQ we used fluorescence datapoints for $[Inc]_{exp}$ = 0-14 nM to fit the calibration curve (Figure S2A-B). For H-y00b21 IncD with d08z16 Rep-FQ we used fluorescence datapoints for $[Inc]_{exp}$ = 0-16nM to fit calibration curve (Figure S2C-D).

This calibration curve was fit using weighted linear regression to a model of the form f(x) = ax + b. The linear regression was inverse weighted with fluorescence in order to account for the Poisson distribution of the photomultiplier counts (Figure S2).

This fit was then used to deduce 2 parameters to convert subsequent fluorescent measurements of reaction kinetics to concentration in nM: $\alpha$ and $\beta$. We define $\alpha$ as the conversion factor which maps the fluorescence in afu to concentration in nM for the quenched reporter (*FQ*), while $\beta$ maps the fluorescence in afu to concentration in nM for the unquenched reporter (*FInc*). Mathematically, we assume

$$\phi = \alpha[FQ] + \beta[FInc], \qquad (S29)$$

where $\phi$ is the fluorescence measurements after performing steps 1 and 2 of normalisation. Under the constraint

$$[F_\mathrm{T}] = [FQ] + [FInc], \quad (S30)$$

where $[F_\mathrm{T}]$ is the total concentration of fluorophore-labelled strand. We can rearrange equations (*S29*) and (*S30*) to obtain

$$\phi = \alpha([F_\mathrm{T}] - [FInc]) + \beta[FInc], \quad (S31)$$

$$\phi = \alpha[F_T] + (\beta - \alpha)[FInc], \quad (S32)$$

$$[FInc] = \frac{\phi - \alpha[F_T]}{\beta - \alpha}. \quad (S33)$$

We assume that all concentrations are given in nM, and therefore $\alpha$ and $\beta$ are dimensionless constants.

We use equation (*S33*) to infer the concentrations of *FInc* in subsequent measurements of reaction kinetics. During the calibration procedure, we determine $\alpha$ and $\beta$ from the form of the linear fit. To do so, however, it is necessary to handle the fact that, due to uncertainties in concentration, the increase in fluorescent signal does not saturate at exactly $[Inc]_{\mathrm{exp}}$ = 15 nM. To be consistent with our assumption that the expected concentration of *FQ* is correct during calibration, we define $\gamma$ as the correction factor such that the injected concentration of incumbent is given by

$$[Inc] = \gamma \cdot [Inc]_{\mathrm{exp}}, \quad (S34)$$

with $\gamma$ chosen so that the plateau in fluorescent signal occurs at exactly $[Inc] = \gamma \cdot [Inc]_{\mathrm{exp}}$ = 15 nM. Having fixed $\gamma$, we can then extract values of $\alpha$ and $\beta$ from the linear fit, using the intercept ($\alpha$[15]) and the gradient ($(\beta - \alpha)\gamma$).

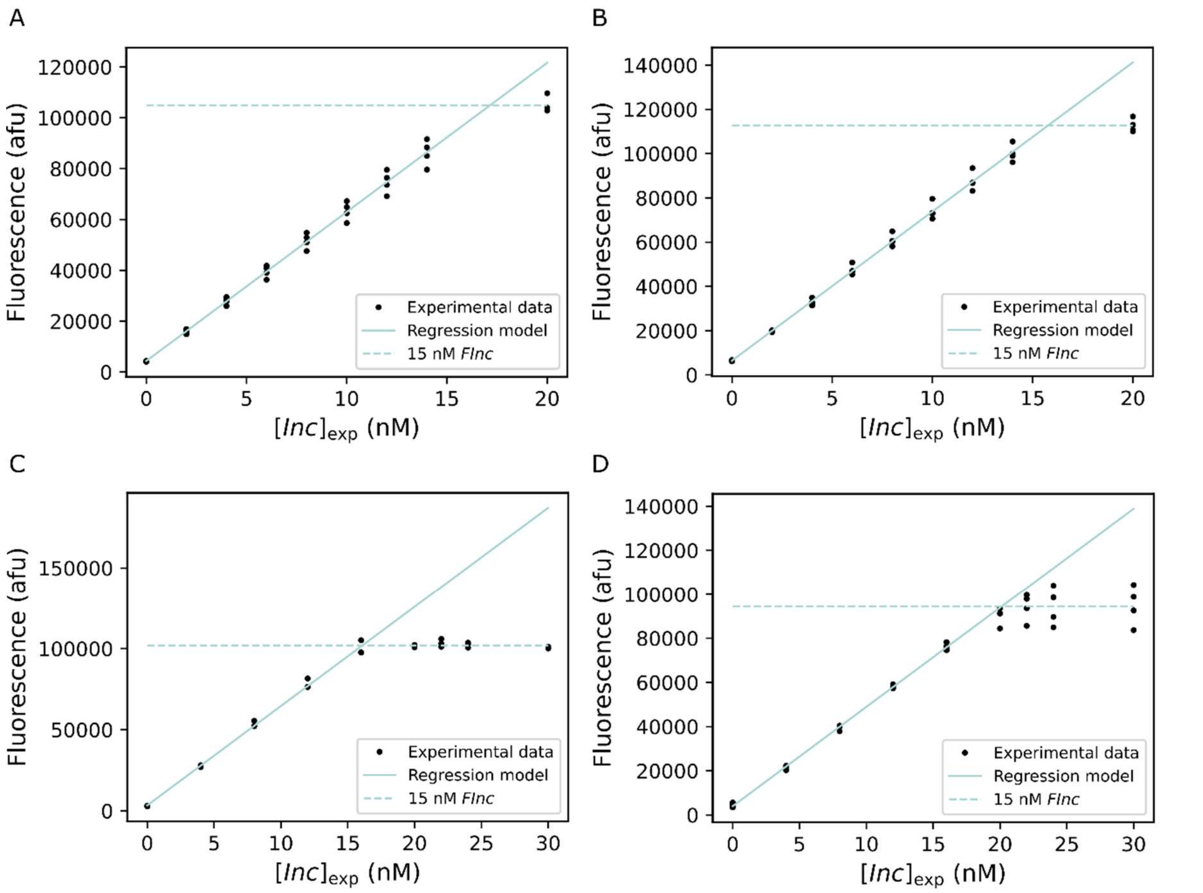

**Figure S2. Calibration curves for conversion of fluorescence readings to concentration of *Inc* in nM.** Calibration curves were fit using weighted linear regression to a model of the form f(x) = ax+b, including only the points that were visually consistent with a linear fit. Calibration curves were generated by combining 15 nM of reporter complex (*FQ*) with a series of different expected *Inc* concentrations ($[Inc]_{\mathrm{exp}}$). Either 3 or 4 replicates were performed for each reporter reaction. The gradient, y-intercept and plateau of these curve are used to derive values of $\alpha$, $\beta$ and $\gamma$, required for consistent conversion of fluorescence readings to concentration in nM. A) Calibration curve for low RNA purine content system 0-20 nM L-y00b26 IncD displacement of 15 nM L-d06z16 Rep-FQ. Gain = 1800, focal height = 5.9 mm, a = 5865, b = 4350, $\alpha$ = 290.0, $\beta$ = 6988, $\gamma$ = 0.8756, R-square = 0.9900. B) Calibration curve for low RNA purine content system 0-20 nM L-y00b26 IncR displacement of 15 nM L-d06z16 Rep-FQ. Gain = 1800, focal height = 5.9 mm, a = 6741, b = 6435, $\alpha$ = 429.0, $\beta$ = 7510, $\gamma$ = 0.9520, R-square = 0.9935. C) Calibration curve for high RNA purine content system 0-30 nM H-y00b21 IncD displacement of 15 nM H-d08z16 Rep-FQ. Gain = 1800, focal height = 5.9, a = 6127, b = 3376, $\alpha$ = 225.0, $\beta$ = 6799, $\gamma$ = 0.9319, R-square = 0.9957. D) Calibration curve for high RNA purine content system 0-30 nM y00b21 IncR displacement of 15 nM d06z16 Rep-FQ. Gain = 1700, focal height = 5.9 mm, a = 4496, b = 3963, $\alpha$ = 264.2, $\beta$ = 6295, $\gamma$ = 0.7455, R-square = 0.9983.

### Supplementary note 7. Reporter characterisation reactions

*Supplementary note 7.1. Experimental set-up for reporter characterisation reactions to estimate* $k_{\mathrm{rep}}$

We characterised all reporter reactions to estimate the reporter rate constant, $k_{\mathrm{rep}}$. As in supplementary note 6.1 these reactions are captured using 4 individual measurements (see illustration in Figure S3). For these experiments we injected 50 μL of 100 nM IncD/IncR with varying volumes of 1 M NaCl + 1X TAE. for final expected IncD/IncR concentrations of 0 nM, 4 nM, 8 nM and 12 nM in 200 μL (detailed protocol in Table S6). We performed 3 or 4 replicates for each of these reporter characterisation reactions.

| Steps | Total reaction volume (μL) | Addition | Expected concentration in 200 μL (nM) | Purpose |
|---|---|---|---|---|
| Measurement 1 | 150 | Reaction: 140 μL of 1 M NaCl + 1X TAE + 10 μL $[FQ]_{\mathrm{exp}}$ = 300 nM | $[FQ]_{\mathrm{exp}}$ = 15 | Estimate residual fluorescence baseline |
| | | Positive control: 3 μL of $[F]_{\mathrm{exp}}$ = 1 μM + 10 μL of $[Inc]_{\mathrm{exp}}$ = 400 nM + 137 μL of 1 M NaCl + 1X TAE | $[FInc]_{\mathrm{PCexp}}$ = 15 | Reference fluorescence value |
| | | Negative control: 140 μL of 1 M NaCl + 1X TAE + 10 μL $[FQ]_{\mathrm{exp}}$ = 300 nM | $[FQ]_{\mathrm{NCexp}}$ = 15 | |
| | | Buffer-only control: 150 μL of 1 M NaCl + 1X TAE | | |
| Measurement 2 | 200 | Reaction: 8-24 μL of $[Inc]_{\mathrm{exp}}$ = 100 nM + 42-26 μL of 1 M NaCl + 1X TAE | Evolve with time | Reaction kinetics |
| | | Positive control: 50 μL of 1 M NaCl + 1X TAE | $[FInc]_{\mathrm{PCexp}}$ = 15 | |

| | | | | |
|---|---|---|---|---|
| | | Negative control: 50 μL of 1 M NaCl + 1X TAE | $[FQ]_{NCexp}$ = 15 | |
| | | Buffer-only control: 50 μL of 1 M NaCl + 1X TAE | | |
| Measurement 3 | 200 | Reaction: - | $[FQ]_{exp}$ = 11-3; $[FInc]_{exp}$ = 4-12 | Estimate $[Inc]_0$ |
| | | Positive control: - | $[FInc]_{PCexp}$ = 15 | |
| | | Negative control: - | $[FQ]_{NCexp}$ = 15 | |
| | | Buffer-only control: - | | |
| Measurement 4 | 210 | Reaction: 10 μL of $[Inc]_{exp}$ = 400 nM | $[SInv]_{exp}$ = 10; $[FInc]_{exp}$ = 15 | Estimate $[FQ]_0$ |
| | | Positive control: 10 μL of 1 M NaCl + 1X TAE | $[FInc]_{PCexp}$ = 15 | |
| | | Negative control: 10 μL of $[Inc]_{exp}$ = 400 nM | $[SInv]_{NCexp}$ = 10; $[FInc]_{NCexp}$ = 15 | Estimate $[FQ]_{NC0}$ |
| | | Buffer-only control: 10 μL of 1 M NaCl + 1X TAE | | |

**Table S6. Detailed experimental protocol for reporter characterisation reactions to estimate $\boldsymbol{k}_{\mathbf{rep}}$.** Summary of volumes of reactants added at each measurement for reporter characterisations to estimate of $k_{\text{rep}}$. *FQ* is added at an expected concentration of 15 nM for all reactions and a series of different expected *Inc* concentrations are added to trigger initiation of the reaction. The expected concentration of reactants and products in 200 μL are given for each measurement. The purpose of each measurement for the further normalisation and post-processing are outlined.

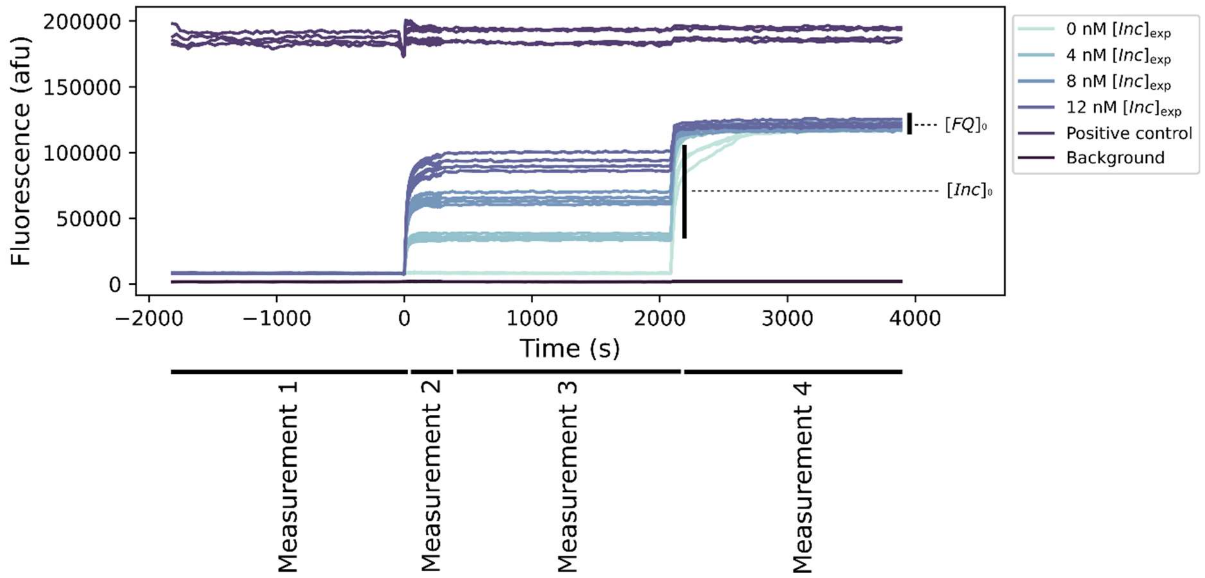

**Figure S3. Example of raw fluorescent traces for reporter characterisation experiment.** Raw fluorescence data from reporter characterisation reaction (15 nM L-z16d06 Rep-FQ + 0-12 nM L-y00b26 IncR) to illustrate the fluorescence measurements and controls for each experiment. Measurement 1 captures the initial reporter (*FQ*) fluorescence. Measurement 2 captures the reaction kinetics after injection of *Inc*. Measurement 3 monitors the steady state fluorescence. Measurement 4 measures the fluorescence after an excess of *Inc* (20 nM) is added. Each reaction is performed in triplicate or quadruplicate. The positive control is composed of 15 nM *FInc* (AlexaFluor488-*Inc*) and the background negative control is 1 M NaCl + 1X TAE.

For the low RNA purine content system, we initially tested L-y00b26 IncD. We confirmed that the reporter reaction was sufficiently rapid to act as an effective readout for the reactions of interest. For this system we estimated a reporter rate constant, $k_{\mathrm{rep}}$, of $3.586 \cdot 10^6$ $M^{-1}$ $s^{-1}$ (Table S9, Figure S6A). We assumed that $k_{\mathrm{rep}}$ was not significantly different for L-y00b21 IncD, L-y00b16 IncD, and L-y00b11 IncD. For DNA>RNA strand displacement systems the reporter reaction involves an RNA incumbent strand displacing the DNA quencher-associated strand. We therefore tested the $k_{\mathrm{rep}}$ for this reporter displacement reaction with L-y00b26 IncR. We estimated a $k_{\mathrm{rep}}$ of $3.675 \cdot 10^6$ $M^{-1}$ $s^{-1}$ for this reaction, confirming the suitability of this reporter system (Figure S6B). Again, we assumed that $k_{\mathrm{rep}}$ was not significantly different for L-y00b21 IncR, L-y00b16 IncR, and L-y00b11 IncR.

For the high purine content design, we were forced to adapt the reporter reaction slightly. The high purine and specifically the high guanine content in the both *Inv* and *Inc* strands complicated the design of this system due increased probability of G-quadruplex formation (5). These secondary structures appeared to decrease $k_{\mathrm{rep}}$ within various designs that we tested which limited their suitability as a readout for the reactions of interest. Ultimately, to overcome these difficulties we selected sequences from previous literature which achieved the high purine restriction while maintaining a relatively fast reporter displacement (6). Having estimated $k_{\mathrm{rep}}$ for reporter H-d06z16 Rep-FQ displacement by H-y00b21 IncR we decided to increase the length of reporter toehold to 8nt in an attempt to maximise $k_{\mathrm{rep}}$ for these systems. Having decided to use known sequences from the literature we were limited in the length of the incumbent strand and as such we only investigated 3 branch migration domain lengths up to 21nt. For H-y00b21 IncD we estimated a $k_{\mathrm{rep}}$ of $1.467 \cdot 10^5$ $M^{-1}$ $s^{-1}$ (Figure S6G). In this case we found significant differences in $k_{\mathrm{rep}}$ between incumbent strands with different branch migration domain lengths. For H-y00b11 IncD and H-y00b16 IncD we estimated $k_{\mathrm{rep}}$ values of $1.790 \cdot 10^6$ $M^{-1}$ $s^{-1}$ and $1.458 \cdot 10^6$ $M^{-1}$ $s^{-1}$, respectively (Figures S6C & S6E). We suggest that the additional nucleotides in the longest incumbent strand may encourage secondary structure which is not present for the shorter incumbent strands, explaining the lower rate constant for this reporter reaction. Finally, we estimated $k_{\mathrm{rep}}$ for the reporter reactions triggered by the RNA incumbent strands of the 3 different branch migration domain lengths. For these reactions we actually used a reporter toehold of 6nt to limit an undesired leak reaction between *SInc* and *FQ*. We estimated $k_{\mathrm{rep}}$ values of $2.033 \cdot 10^5$ $M^{-1}$ $s^{-1}$, $1.637 \cdot 10^5$ $M^{-1}$ $s^{-1}$ and $9.287 \cdot 10^4$ $M^{-1}$ $s^{-1}$ for H-y00b11 IncR, H-y00b16 IncR and H-y00b21 IncR, respectively (Figure S6D, S6F & S6H).

*Supplementary note 7.2. Normalisation and data-processing of reporter characterisation experiments for $k_{\mathrm{rep}}$ estimation*

Step 1: Removing background fluorescence produced by experimental buffer.

The first step in processing all data is to remove the background autofluorescence resulting from the 1 M NaCl + 1X TAE buffer. At each time point the fluorescence of the buffer-only negative control was subtracted from the raw fluorescence values for all other wells (experimental reactions, positive and negative controls).

Step 2: Correction based on the positive control fluorescence

As in Supplementary Note 6, we use the positive control to correct for fluctuations in fluorescence resulting from variable environmental conditions. We define the mean fluorescence of the positive control during measurement 1 as the reference fluorescence value to which all other fluorescence is compared. We can assume that during the first measurement any photobleaching is negligible and all wells should be at the experimental temperature because plates were left for 30 minutes at the experimental temperature prior to the first measurement. We correct for these fluctuations using the same approach as in Step 2 of Supplementary note 6.2, with the calculation dependent on whether 'well mode' or 'plate mode' is used to measure the reaction kinetics.

Step 3: Estimation of $[FQ]_0$

We next calculate the initial (t = 0 s) concentration of *FQ*. We calculate the mean fluorescence corresponding to complete dissociation of *FQ* for each reaction (measurement 4 in Table S6). In this case we assume that the fluorescence of measurement 4 corresponds to 100% unquenched *F* where all *FQ* has been converted into fluorescent product *FInc*. Under this assumption we can determine the initial concentration of *FQ* ($[FQ]_{\mathrm{est0}}$) by dividing the mean fluorescence of measurement 4 by $\beta$.

Step 4: Conversion of fluorescence data to concentrations using $\alpha$ and $\beta$

We employ the values of $\alpha$ and $\beta$ from the corresponding calibration curve in order to convert all data points in the reaction dynamics into inferred concentrations according to equation (*S33*). Here, $[F_T]$ is the total concentration of $[FQ]_0$ as estimated in step 3 (Table S6). The normalised fluorescent traces for all reporter characterisation experiments are given in Figure S4A-H). Applying equation (*S33*) to steady state (measurement 3) we obtain an estimate of $[Inc]_0$ ($[Inc]_{est0}$).

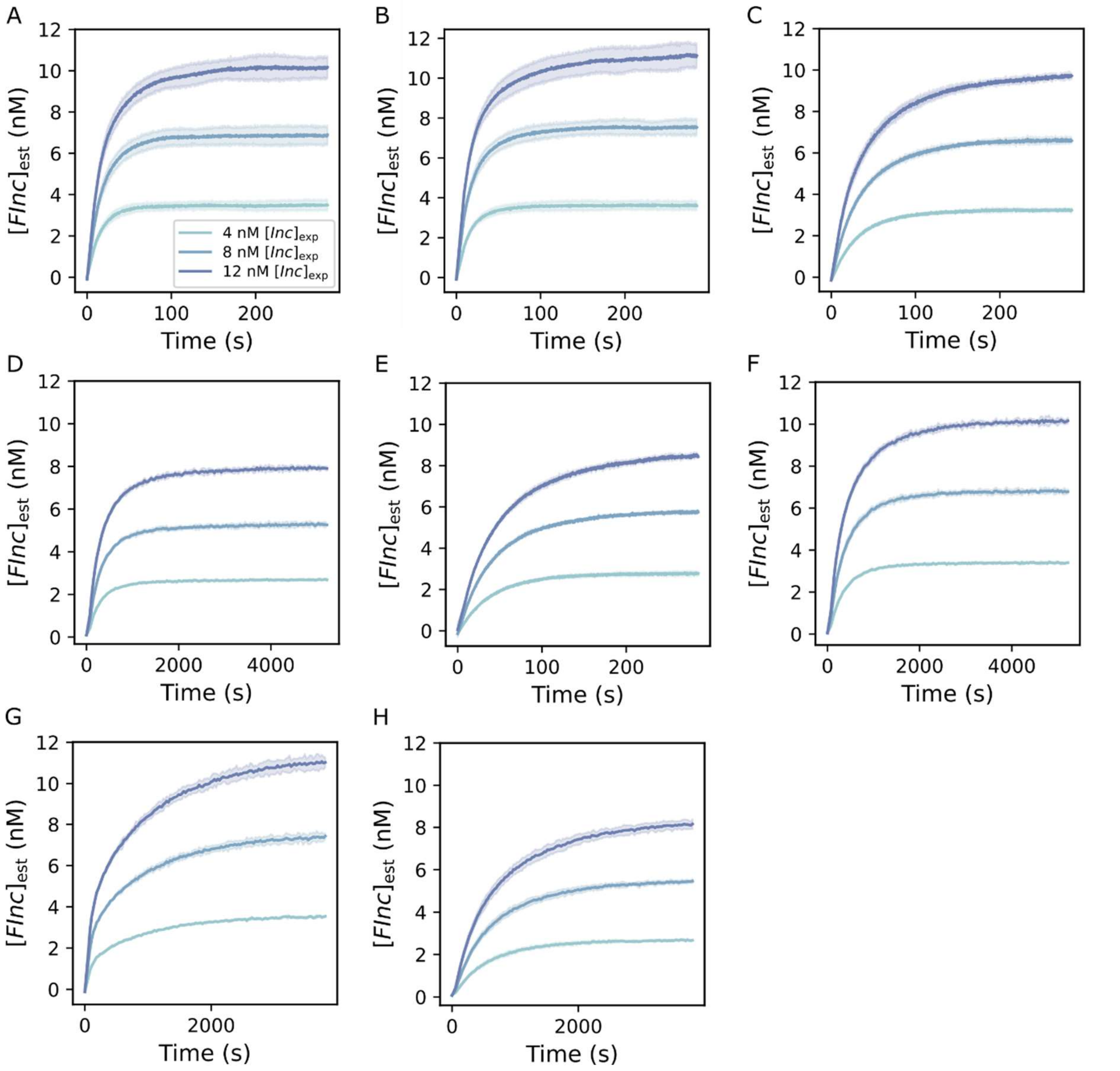

**Figure S4. Processed fluorescent traces for all reporter characterisation reactions.** Processed fluorescent outputs for reporter reactions in which 15 nM reporter *FQ* is combined with 4-12 nM of an incumbent strand. Each reaction is performed in triplicate or quadruplicate. Data is processed to convert fluorescence readings to an estimated concentration of the fluorescent product, *FInc*, in nM and to account for fluctuations due to changes in temperature and reaction volume. For the low RNA purine content design 1 branch migration domain length was characterised (26nt). For the high RNA purine content 3 different branch migration domain lengths were tested (11-21nt). This normalised fluorescence data is used to estimate $k_{\mathrm{rep}}$ for each reporter reaction. A) L-y00b26 IncD displacement of L-d06z16 Rep-FQ. B) L-y00b26 IncR displacement of L-d06z16 Rep-FQ. C) H-y00b11 IncD displacement of H-d08z16 Rep-FQ. D) H-y00b11 IncR displacement of H-d06z16 Rep-FQ. E) H-y00b16 IncD displacement of H-d08z16 Rep-FQ. F) H-y00b16 IncR displacement of H-d06z16 Rep-FQ. G) H-y00b21 IncD displacement of H-d08z16 Rep-FQ. H)  H-y00b21 IncR displacement of H-d06z16 Rep-FQ. The legend given in A corresponds to all subplots.

*Supplementary note 7.3. Fitting ordinary differential equations (ODEs) to reporter characterisation experiments for $k_{\mathrm{rep}}$ estimation*

We present two alternative approaches to fitting $k_{\mathrm{rep}}$: individual and global fits. Individual fitting obtains fits and estimates $k_{\mathrm{rep}}$ for each kinetic trace for a given set of strands, while global fitting yields a single $k_{\mathrm{rep}}$ estimate across all replicates and $[Inc]_{\mathrm{exp}}$ for a given set of strands. In the main text we report only the $k_{\mathrm{rep}}$ estimates from global fits, however we present both here as evidence that our $k_{\mathrm{rep}}$ estimates are robust to the fitting approach.

For individual fits, we employ the python package scipy.integrate.odeint to integrate the ODE model for reporter characterisation reactions (equation (*2*)). This package accepts an ODE system, initial reactant and product concentrations and the timesteps over which the reaction is measured as inputs. We provide fixed, initial concentrations of $[FQ]_{\mathrm{est0}}$ estimated from measurement 4 (Table S6) and 0 nM for both [*FInc*] and [*Q*]. We set the integration step size (hmax) at 20. For each kinetic trace we fit $k_{\mathrm{rep}}$, along with the initial incumbent concentration $[Inc]_0$. Notably, we log-transform $k_{\mathrm{rep}}$ for the purpose of fitting given that $k_{\mathrm{rep}}$ can vary over several orders of magnitude. We feed this ODE solver into a second function which fits the solution of the numerical integration to the experimental fluorescent data by varying $k_{\mathrm{rep}}$ and $[Inc]_0$. For this purpose we employ the scipy.optimize.curve_fit function. We define the estimated concentration outputs of this fitting procedure as $[Inc]_{\mathrm{fit0}}$. This function accepts an initial prediction of $[Inc]_{\mathrm{fit0}}$ and $k_{\mathrm{rep}}$ as inputs as well as upper and lower bounds for these systems. For all reporter characterisations we set an initial $k_{\mathrm{rep}}$ prediction of $10^6$ $M^{-1}$ $s^{-1}$, with lower and upper bounds of $10^1$ $M^{-1}$ $s^{-1}$ and $10^8$ $M^{-1}$ $s^{-1}$, respectively. For $[Inc]_{\mathrm{fit0}}$ we set the initial prediction as the value of *Inc* estimated as the mean from the steady state measurement 3 ($[Inc]_{\mathrm{est0}}$) and we set lower and upper bounds of $[Inc]_{\mathrm{est0}}/2$ and $2[Inc]_{\mathrm{est0}}$. The output of this function are optimised estimates of $k_{\mathrm{rep}}$ and $[Inc]_{\mathrm{fit0}}$ for each individual kinetic trace. We obtain an estimate for the mean $k_{\mathrm{rep}}$ from the inverse log-transformed $k_{\mathrm{rep}}$ values across all traces for each set of strands and calculate the standard error by considering the variability in these individual estimates (Table S7, with fits illustrated in Figure S5).

| Reporter system | Mean $k_{\mathrm{rep}}$ ($M^{-1}$ $s^{-1}$) | Standard error ($M^{-1}$ $s^{-1}$) |
|---|---|---|
| L-y00b26 IncD + L-d06z16 Rep-FQ | $4.141 \cdot 10^{6}$ | $0.327 \cdot 10^{6}$ |
| L-y00b26 IncR + L-d06z16 Rep-FQ | $4.283 \cdot 10^{6}$ | $0.150 \cdot 10^{6}$ |
| H-y00b21 IncD + H-d08z16 Rep-FQ | $1.569 \cdot 10^{5}$ | $0.145 \cdot 10^{5}$ |
| H-y00b21 IncR + H-d06z16 Rep-FQ | $1.178 \cdot 10^{5}$ | $0.055 \cdot 10^{5}$ |
| H- y00b16 IncD + H-d08z16 Rep-FQ | $1.705 \cdot 10^{6}$ | $0.083 \cdot 10^{6}$ |
| H-y00b16 IncR + H-d06z16 Rep-FQ | $1.752 \cdot 10^{5}$ | $0.088 \cdot 10^{5}$ |
| H-y00b11 IncD + H-d08z16 Rep-FQ | $2.085 \cdot 10^{6}$ | $0.093 \cdot 10^{6}$ |
| H-y00b11 IncR + H-d06z16 Rep-FQ | $2.172 \cdot 10^{5}$ | $0.032 \cdot 10^{5}$ |

**Table S7. $k_{\mathrm{rep}}$ estimates from averaging over individual reporter characterisation reactions.** Mean reporter rate constant, $k_{\mathrm{rep}}$, estimate and standard error from individual fits of 3 or 4 replicates at 3 different expected *Inc* ($[Inc]_{\mathrm{exp}}$ = 4-12 nM) concentrations. $k_{\mathrm{rep}}$ estimates were obtained by fitting an ODE model of reporter strand displacement to each individual fluorescence curve.

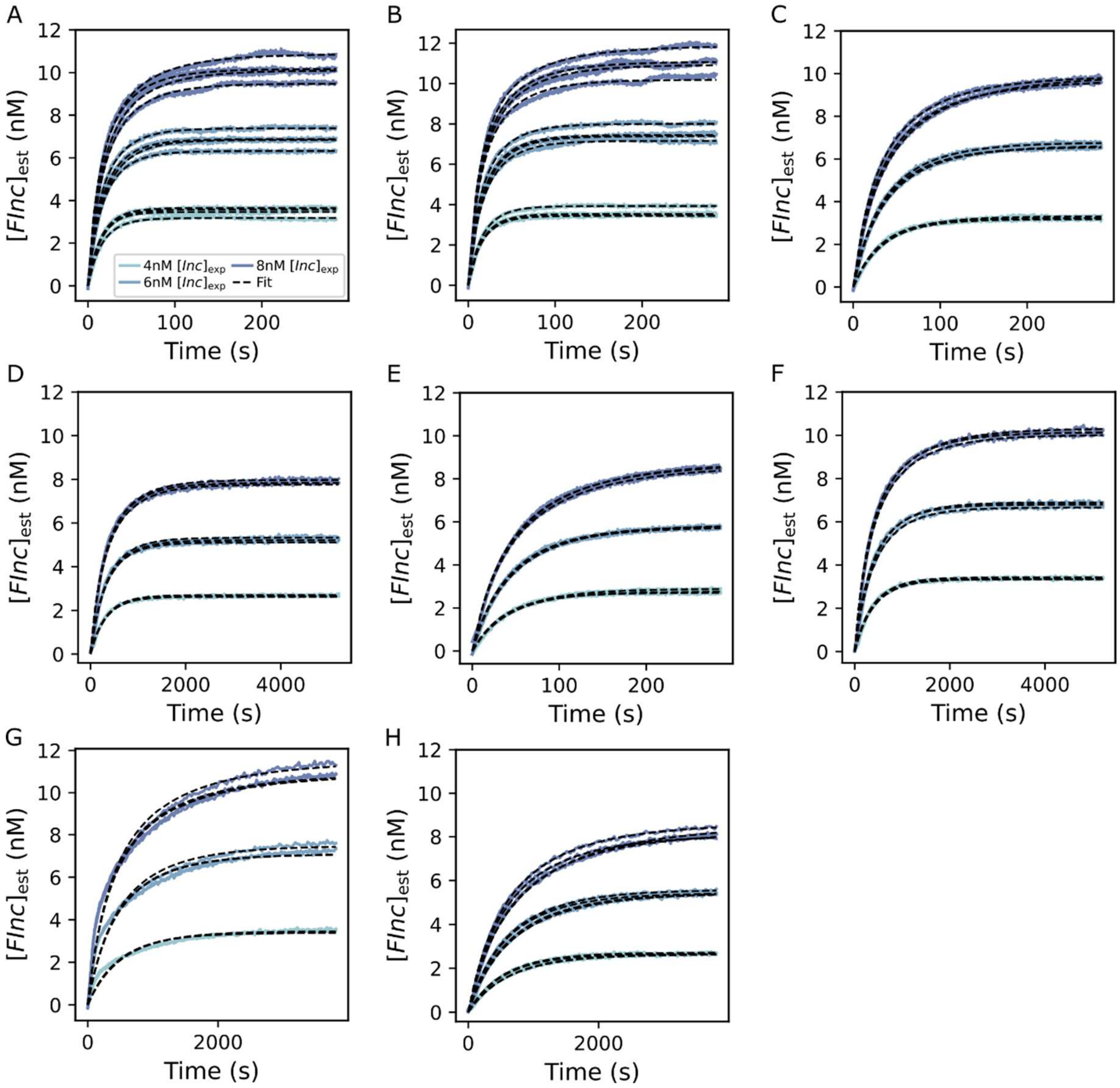

**Figure S5. Individual fits to fluorescent traces in reporter characterisation reactions.** Individual fits of an ODE model to processed fluorescent data from each reporter characterisation experiment. Each experiment had either 3 or 4 replicates and were performed across 3 different expected *Inc* ($[\mathrm{Inc}]_{\mathrm{exp}}$ = 4-12 nM) concentrations. An expected reporter concentration ($[FQ]_{\mathrm{exp}}$) of 15 nM was used for all experiments. Black, dotted lines represent fits of the ODE model and solid, coloured lines represent the normalised experimental fluorescent data. A) L-y00b26 IncD displacement of L-d06z16 Rep-FQ. B) L-y00b26 IncR displacement of L-d06z16 Rep-FQ. C) H-y00b11 IncD displacement of H-d08z16 Rep-FQ. D) H-y00b11 IncR displacement of H-d06z16 Rep-FQ. E) H-y00b16 IncD displacement of H-d08z16 Rep-FQ. F) H-y00b16 IncR displacement of H-d06z16 Rep-FQ. G) H-y00b21 IncD displacement of H-d08z16 Rep-FQ. H) H-y00b21 IncR displacement of H-d06z16 Rep-FQ. The legend given in A corresponds to all subplots.

We also determined a global estimate for $k_{\mathrm{rep}}$ across all replicates and $[Inc]_{\mathrm{exp}}$ concentrations for each set of strands (Figure S6). In this case we use a similar function to scipy.integrate.odeint known as jax.experimental.ode.odeint. Notably, this function allows for improved parallelisation as compared to scipy.integrate.odeint and reduces the computational power required. As for the individual fits, we provide initial concentrations of $[FQ]_{\mathrm{est0}}$ estimated from measurement 4 (Table S6) and 0 nM for both $[FInc]$ and $[Q]$. We next employed an alternative function to fit the solution of the numerical integration to the experimental data. We utilised scipy.optimize.curve_fit to fit the solution of the integration to the experimental data and output an optimised $k_{\mathrm{rep}}$ value fitted to all fluorescence curves for the same strands simultaneously. To fit a single, global estimate of $k_{\mathrm{rep}}$ for each set of strands we fixed $[Inc]_0$ at $[Inc]_{\mathrm{est0}}$. Finally, in order to obtain a sensible error for this $k_{\mathrm{rep}}$ estimate we used a jackknife (leave-one-out) approach (Table S8). We estimated the global $k_{\mathrm{rep}}$ each time leaving out a different curve ($i$) within each experiment, yielding a total of $n$ jackknife replicates ($k_{\mathrm{rep}_{(i)}}$), where $n$ is the number of curves. These jackknife replicates were then inverse log transformed and the final jackknife estimate of $\overline{k_{\mathrm{rep}}}_{\mathrm{jack}}$ is given by

$$\overline{k_{\mathrm{rep}}}_{\mathrm{jack}} = n \cdot k_{\mathrm{rep}} - (n-1) \cdot \frac{1}{n} \sum_{i=1}^{n} k_{\mathrm{rep}_{(i)}}, \qquad (S35)$$

which removes bias from the initial global $k_{\mathrm{rep}}$ estimate (7).

We calculated the jackknife standard error associated with this $\overline{k_{\mathrm{rep}}}_{jack}$ estimate according to

$$SE\left(\overline{k_{\mathrm{rep}}}_{\mathrm{jack}}\right) = \sqrt{\frac{n-1}{n} \sum_{i=1}^{n} \left(k_{\mathrm{rep}_{(i)}} - k_{\mathrm{rep}}\right)^2}. \qquad (S36)$$

Plots of fits using a single, global $k_{\mathrm{rep}}$ for each set of strands are given in Figure S6. We used the global jackknife estimates of $k_{\mathrm{rep}}$ as fixed values for fitting of full toehold exchange reactions.

| Reporter system | Jackknife global $k_{\mathrm{rep}}$ (M$^{-1}$ s$^{-1}$) | Jackknife standard error (M$^{-1}$ s$^{-1}$) |
|---|---|---|
| Low purine y00b26 IncD + d06z16 Rep-FQ | $3.586 \cdot 10^6$ | $0.071 \cdot 10^6$ |
| Low purine y00b26 IncR + d06z16 Rep-FQ | $3.675 \cdot 10^6$ | $0.106 \cdot 10^6$ |
| High purine y00b21 IncD + d08z16 Rep-FQ | $1.467 \cdot 10^5$ | $0.067 \cdot 10^5$ |
| High purine y00b21 IncR + d06z16 Rep-FQ | $9.287 \cdot 10^4$ | $0.357 \cdot 10^4$ |
| High purine y00b16 IncD + d08z16 Rep-FQ | $1.458 \cdot 10^6$ | $0.026 \cdot 10^6$ |
| High purine y00b16 IncR + d06z16 Rep-FQ | $1.637 \cdot 10^5$ | $0.037 \cdot 10^5$ |
| High purine y00b11 IncD + d08z16 Rep-FQ | $1.790 \cdot 10^6$ | $0.043 \cdot 10^6$ |
| High purine y00b11 IncR + d06z16 Rep-FQ | $2.033 \cdot 10^5$ | $0.023 \cdot 10^5$ |

**Table S8. Global $k_{\mathrm{rep}}$ estimates across reporter characterisation reactions.** Global $k_{\mathrm{rep}}$ estimates and standard errors across 3 or 4 replicates at3 different expected *Inc* ($[Inc]_{\mathrm{exp}}$ = 4-12 nM) concentrations. $k_{\mathrm{rep}}$ estimates were obtained by fitting an ODE model of the reporter strand displacement reaction to all fluorescent curves simultaneously for each experiment. $k_{\mathrm{rep}}$ estimates and standard errors were calculated using a jackknife (leave-one-out) approach.

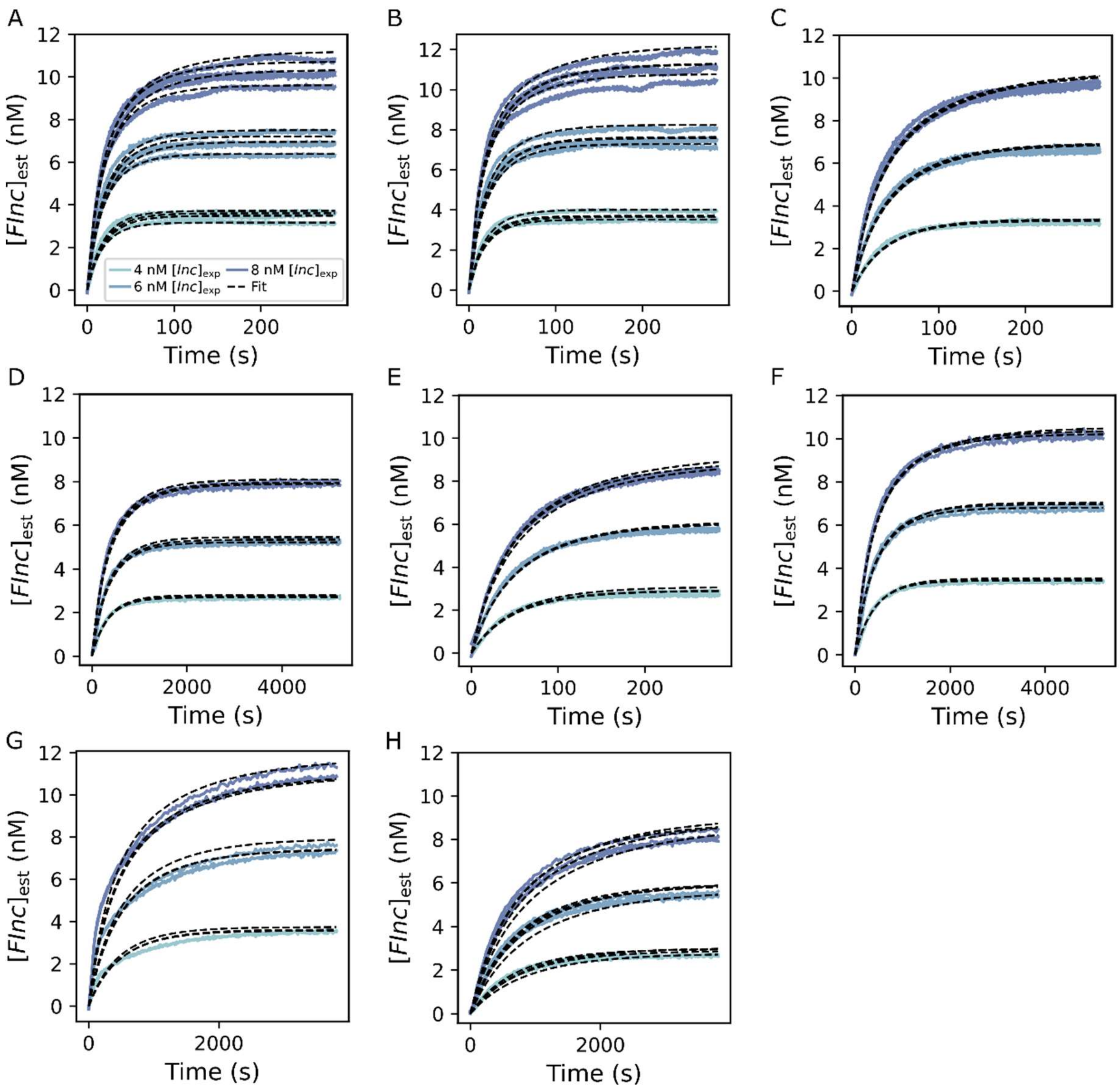

**Figure S6. Global fits to fluorescent traces in reporter characterisation reactions.** Global fits of ODE model to processed fluorescent data from each reporter characterisation experiment. Global fits for each set of strands were derived from 3 or 4 replicates at 3 different, expected *Inc* ($[Inc]_{exp}$ = 4-12 nM) concentrations. An expected reporter concentration ($[FQ]_{exp}$) of 15 nM was used for all experiments. $k_{\mathrm{rep}}$ estimates and standard errors were calculated using a jackknife (leave-one-out) approach. Black, dotted lines represent fit of ODE model and solid, coloured lines represent the normalised experimental fluorescent data. A) L-y00b26 IncD displacement of L-d06z16 Rep-FQ. B) L-y00b26 IncR displacement of L-d06z16 Rep-FQ. C) H-y00b11 IncD displacement of H-d08z16 Rep-FQ. D) H-y00b11 IncR displacement of H-d06z16 Rep-FQ. E) H-y00b16 IncD displacement of H-d08z16 Rep-FQ. F) H-y00b16 IncR displacement of H-d06z16 Rep-FQ. G) H-y00b21 IncD displacement of H-d08z16 Rep-FQ. H) H-y00b21 IncR displacement of H-d06z16 Rep-FQ. The legend given in A corresponds to all subplots.

### Supplementary note 8. Full strand displacement reactions

*Supplementary note 8.1. Experimental set-up for full strand displacement reactions*

RNA>DNA strand displacement reactions and DNA>DNA strand displacement reactions were performed in the same manner. Each experiment was performed in triplicate. 10 µL of reporter complex *FQ* (300 nM) was added to 130 µL of 1 M NaCl + 1X TAE for a final expected concentration of $[FQ]_{exp}$ = 15 nM in 200 µL. Different volumes of 100 nM *Inv* were added in a total of 50 µL for a final expected concentration of $[Inv]_{exp}$ = 4-8 nM in 200 µL. In this way we could confirm that *Inv* and *FQ* were orthogonal. Next 10 µL of *SInc* complex (200 nM) was automatically injected for a final expected concentration of $[SInc]_{exp}$ = 10 nM in 200 µL. When the system had reached steady state, we next added 10 µL of excess *Inv* (400 nM) for an expected additional concentration of 20 nM to trigger dissociation of all *SInc* complexes. We then added 10 µL of 400 nM *Inc* for an expected additional concentration of 20 nM to trigger all *FQ* complexes to dissociate (full protocols in Table S9, illustrative fluorescent data in Figure S7). As with the reporter characterisations (supplementary note 7.1), each replicate included a buffer-only negative control (1 M NaCl + 1X TAE) and a positive control composed of $[F]_{exp}$ = 15 nM and an excess of *Inc* ($[Inc]_{exp}$ = 20 nM) to account for any changes in the fluorescence as a result of temperature or reaction volume. There was a second negative control composed of $[FQ]_{exp}$ = 15 nM and $[SInc]_{exp}$ = 10 nM but with $[Inv]_{exp}$ = 0 nM. The volume of each control was consistent with the main assay for each measurement by injecting corresponding volumes of 1 M NaCl + 1X TAE.

For all RNA>DNA and DNA>DNA toehold exchange reactions we captured the reaction kinetics and estimated the initial concentration of reactants with a total of 5 individual measurements.

<u>Measurement 1:</u> Measurement 1 captured the residual background fluorescence of the quenched *FQ* in the presence of *Inv* in 190 µL.

<u>Measurement 2:</u> Measurement 2 captured fluorescence changes after automatic injection of 10 µL of *SInc* complex in 200 µL. The 20% excess of *Inc* in the *SInc* complex gave an initial increase in the fluorescence corresponding to [*exInc*]. The actual reaction kinetics then follows over the rest of the measurement. Again, for systems with rapid kinetics ($\gamma$ > 6nt) we used the 'well mode' of the microplate reader to ensure that we collected sufficient datapoints for these rapid reactions. For the rest of the reactions with slower kinetics we used the 'plate mode'.

<u>Measurement 3:</u> Measurement 3 monitored the fluorescence once the reaction had reached steady state, allowing for estimation of the initial *Inv* concentration ($[Inv]_0$) for each replicate.

<u>Measurement 4:</u> Measurement 4 captured the fluorescence corresponding to complete dissociation of *SInc* complexes and allows for calculation of $[SInc]_0$.

<u>Measurement 5:</u> The fluorescence resulting from complete dissociation of *FQ* is captured by measurement 5 and gives an estimate of $[FQ]_0$.

| Steps | Total reaction volume (μL) | Addition | Expected concentration in 200 μL (nM) | Purpose |
|---|---|---|---|---|
| Measurement 1 | 190 | Reaction: 10 μL $[FQ]_{exp}$ = 300 nM + 8-16 μL of $[Inv]_{exp}$ = 100 nM + 172-164 μL of 1 M NaCl + 1X TAE | $[FQ]_{exp}$ = 15; $[Inv]_{exp}$ = 4-8 | Estimate residual fluorescence baseline |
| | | Positive control: 3 μL $[F]_{exp}$ = 1 μM + 10 μL $[Inc]_{exp}$ = 400 nM + 187 μL of 1 M NaCl + 1X TAE | $[FInc]_{PCexp}$ = 15 | Reference fluorescence value |
| | | Negative control: 10 μL $[FQ]_{exp}$ = 300 nM + 190 μL of 1 M NaCl + 1X TAE | $[FQ]_{NCexp}$ = 15 | |
| | | Buffer-only control: 190 μL of 1 M NaCl + 1X TAE | | |
| Measurement 2 | 200 | Reaction: 10 μL of $[SInc]_{exp}$ = 200 nM | Evolve with time | Reaction kinetics |
| | | Positive control: 10 μL of 1 M NaCl + 1X TAE | $[FInc]_{PCexp}$ = 15 | |
| | | Negative control: 10 μL of $[SInc]_{exp}$ = 200 nM | $[FQ]_{NCexp}$ = 13; $[SInc]_{NCexp}$ = 10; $[FInc]_{NCexp}$ = 2 | |
| | | Buffer-only control: 10 μL of 1M NaCl + 1X TAE | | |
| Measurement 3 | 200 | Reaction: - | $[FQ]_{exp}$ = 9-5; $[SInv]_{exp}$ = 4-8; $[FInc]_{exp}$ = 6-10; | Estimate $[Inv]_0$ |
| | | Positive control: - | $[FInc]_{PCexp}$ = 15 | |
| | | Negative control: - | $[FQ]_{NCexp}$ = 13; $[SInc]_{NCexp}$ = 10; $[FInc]_{NCexp}$ = 2 | Estimate $[exInc]_{NC0}$ |
| | | Buffer-only control: - | | |
| Measurement 4 | 210 | Reaction: | $[FQ]_{exp}$ = 3; $[SInv]_{exp}$ = 10; | Estimate $[SInc]_0$ |

| | | | | |
|---|---|---|---|---|
| | | 10 µL of $[Inv]_{exp}$ =<br>400 nM | $[FInc]_{exp}$ = 12; | |
| | | Positive control:<br>10 µL of 1 M NaCl<br>+ 1X TAE | $[FInc]_{PCexp}$ = 15 | |
| | | Negative control:<br>10 µL of $[Inv]_{exp}$ =<br>400 nM | $[FQ]_{NCexp}$ = 3;<br>$[SInv]_{NCexp}$ = 10;<br>$[FInc]_{NCexp}$ = 12 | |
| | | Buffer-only control:<br>10 µl of 1 M NaCl<br>+ 1X TAE | | |
| Measurement 5 | 220 | Reaction:<br>10 µL of $[Inc]_{exp}$ =<br>400 nM | $[SInv]_{exp}$ = 10;<br>$[FInc]_{exp}$ = 15 | Estimate $[FQ]_0$ |
| | | Positive control:<br>10 µL of 1 M NaCl<br>+ 1X TAE | $[FInc]_{PCexp}$ = 15 | |
| | | Negative control:<br>10 µL of $[Inc]_{exp}$ =<br>400 nM | $[SInv]_{NCexp}$ = 10;<br>$[FInc]_{NCexp}$ = 15 | Estimate $[FQ]_{NC0}$ |
| | | Buffer-only control:<br>10 µL of 1 M NaCl<br>+ 1X TAE | | |

**Table S9. Detailed experimental protocol for DNA>DNA and RNA>DNA strand displacement reactions to estimate $k_{\mathrm{eff}}$.** Summary of volumes of reactants added at each measurement for full RNA>DNA and DNA>DNA strand displacement reactions for both high and low RNA purine content designs. *FQ* is added at an expected concentration of 15 nM and *SInc* is added at an expected concentration of 10 nM for all reactions and a series of different expected *Inv* concentrations are added to trigger initiation of the reaction. The expected concentration of reactants and products in 200 µL are given for each measurement. The purpose of each measurement for the further normalisation and post-processing are outlined.

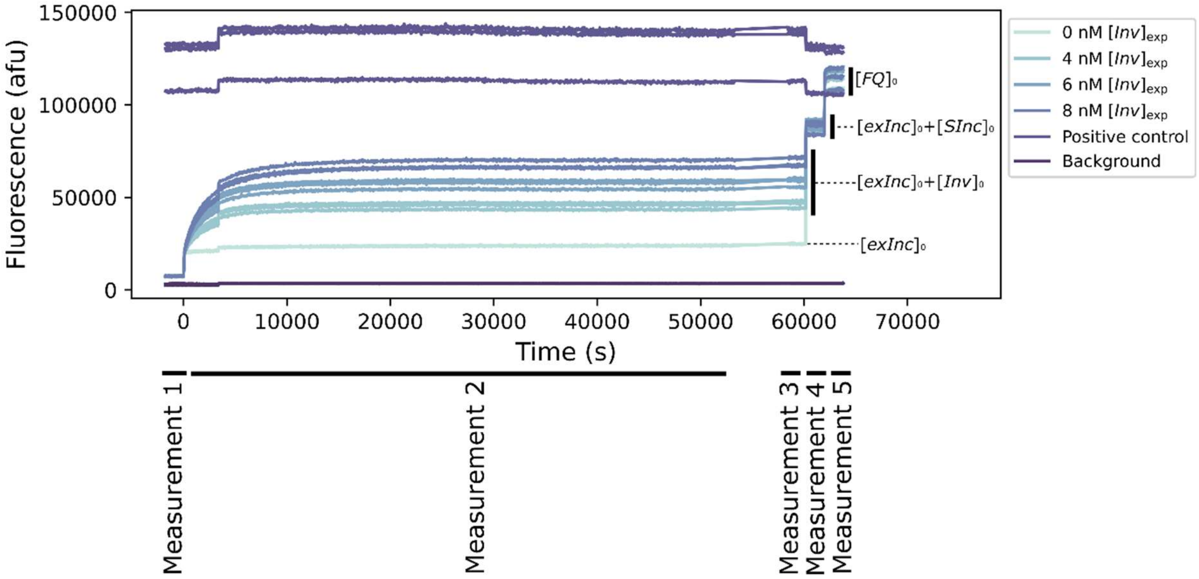

**Figure S7. Example of raw fluorescent traces for DNA>DNA and RNA>DNA strand displacement experiment.** Raw fluorescence data from $DNA_{py}$>$DNA_{py}$ strand displacement reaction (15 nM L-d06z16 Rep-FQ + 10 nM L-y04b11 SIncD with 20% excess L-y00b11 IncD + 0-8 nM L-y10b26 InvD) to illustrate the fluorescence measurements and controls for each experiment. The same measurement approach was used for $DNA_{pu}$>$DNA_{pu}$ and all RNA>DNA strand displacement experiments. Measurement 1 captured the initial reporter *FQ* fluorescence in the presence of *Inv*. Measurement 2 captured the reaction kinetics after injection of *SInc*. Measurement 3 monitored the steady state fluorescence. Measurement 4 measured the fluorescence after an excess of *Inv* (~20 nM) is added. Measurement 5 monitored the fluorescence after an excess of *Inc* (~20 nM) is added. Each reaction is performed in triplicate. The positive control is composed of 15 nM *FInc* (AlexaFluor488-*Inc*) and the background negative control is 1 M NaCl + 1X TAE.

For $DNA_{py}$>$RNA_{py}$ strand displacement reactions we adapted this experimental protocol in order to minimise the volume of RNA lost during injection (full protocol in Table S10). In this case the DNA-RNA *SInc* complex was added manually prior to measurement 2 and the DNA *Inv* was injected via the automated injector prior to measurement 3. The additions for the reactions of interest and the controls are specified alongside an example raw fluorescence trace in Figure S8.

| Steps | Total reaction volume (µL) | Addition | Expected concentration in 200 µL (nM) | Purpose |
|---|---|---|---|---|
| Measurement 1 | 140 | Reaction: 10 µL [*FQ*]$_{exp}$ = 300 nM + 130 µL of 1 M NaCl + 1X TAE | [*FQ*]$_{exp}$ = 15 | Estimate residual fluorescence baseline |
| | | Positive control: 3 µL [*F*]$_{exp}$ = 1 µM + 10 µL [Inc]$_{exp}$ = 400 nM + 127 µL 1 M NaCl + 1X TAE | [*FInc*]$_{Pcexp}$ = 15 | Reference fluorescence value |
| | | Negative control: 10 µL [FQ]$_{exp}$ = 300 nM + 130 µL of 1 M NaCl + 1X TAE | [*FQ*]$_{Ncexp}$ = 15; [*SInc*]$_{Ncexp}$ = 10 | |
| | | Buffer-only control: 140 µL of 1 M NaCl + 1X TAE | | |
| Measurement 2 | 150 | Reaction: 10 µL of [*SInc*]$_{exp}$ = 200 nM | [*FQ*]$_{exp}$ = 13; [*SInc*]$_{exp}$ = 10; [*FInc*]$_{exp}$ = 2 | - |
| | | Positive control: 10 µL of 1 M NaCl + 1X TAE | [*FInc*]$_{Pcexp}$ = 15 | |
| | | Negative control: 10 µL of [*Sinc*]$_{exp}$ = 200 nM | [*FQ*]$_{Ncexp}$ = 13; [*SInc*]$_{Ncexp}$ = 10; [*FInc*]$_{Ncexp}$ = 2 | |
| | | Buffer-only control: 10 µL of 1 M NaCl + 1X TAE | | |
| Measurement 3 | 200 | Reaction: 8-16 µL of [*Inv*]$_{exp}$ = 100 nM + 42-34 µL | Evolve with time | Reaction kinetics |

| | | | | |
|---|---|---|---|---|
| | | of 1 M NaCl + 1X TAE | | |
| | | Positive control: 50 µL of 1 M NaCl + 1X TAE | $[\mathit{FInc}]_{Pcexp}$ = 15 | |
| | | Negative control: 50 µL of 1 M NaCl + 1X TAE | $[\mathit{FQ}]_{Ncexp}$ = 13; $[\mathit{SInc}]_{Ncexp}$ = 10; $[\mathit{FInc}]_{Ncexp}$ = 2 | |
| | | Buffer-only control: 50 µL of 1 M NaCl + 1X TAE | | |
| Measurement 4 | 200 | Reaction: - | $[\mathit{FQ}]_{exp}$ = 9-5; $[\mathit{SInv}]_{exp}$ = 4-8; $[\mathit{FInc}]_{exp}$ = 6-10 | Estimate $[\mathit{Inv}]_0$ |
| | | Positive control: - | $[\mathit{FInc}]_{Pcexp}$ = 15 | |
| | | Negative control: - | $[\mathit{FQ}]_{Ncexp}$ = 13; $[\mathit{SInc}]_{Ncexp}$ = 10; $[\mathit{FInc}]_{Ncexp}$ = 2 | Estimate $[\mathit{exInc}]_{NC0}$ |
| | | Buffer-only control: - | | |
| Measurement 5 | 210 | Reaction: 10 µL of $[\mathit{Inv}]_{exp}$ = 400 nM | $[\mathit{FQ}]_{exp}$ = 3; $[\mathit{SInv}]_{exp}$ = 10; $[\mathit{FInc}]_{exp}$ = 12 | Estimate $[\mathit{SInc}]_0$ |
| | | Positive control: 10 µL of 1 M NaCl + 1X TAE | $[\mathit{FInc}]_{Pcexp}$ = 15 | |
| | | Negative control: 10 µL of $[\mathit{Inv}]_{exp}$ = 400 nM | $[\mathit{FQ}]_{Ncexp}$ = 3; $[\mathit{SInv}]_{Ncexp}$ = 10; $[\mathit{FInc}]_{Ncexp}$ = 12 | |
| | | Buffer-only control: 10 µL of 1 M NaCl + 1X TAE | | |
| Measurement 6 | 220 | Reaction: 10 µL of $[\mathit{Inc}]_{exp}$ = 400 nM | $[\mathit{SInv}]_{exp}$ = 10; $[\mathit{FInc}]_{exp}$ = 15 | Estimate $[\mathit{FQ}]_0$ |

| | | | |
|---|---|---|---|
| | Positive control: 10 µL of 1 M NaCl + 1X TAE | $[FInc]_{\text{Pcexp}}$ = 15 | |
| | Negative control: 10 µL of $[Inc]_{\text{exp}}$ = 400 nM | $[SInv]_{\text{Ncexp}}$ = 10; $[FInc]_{\text{Ncexp}}$ = 15 | Estimate $[FQ]_{\text{NC0}}$ |
| | Buffer-only control: 10 µL of 1M NaCl + 1X TAE | | |

**Table S10. Detailed experimental protocol for $DNA_{py}$>$RNA_{py}$ strand displacement reactions to estimate $k_{\text{eff}}$.** Summary of volumes of reactants added at each measurement for full DNA>RNA strand displacement reactions. *FQ* is added at an expected concentration of 15 nM, *SInc* is added at an expected concentration of 10 nM for all reactions, and a series of different expected *Inv* (4-8 nM) concentrations are added to trigger initiation of the reaction. The expected concentration of reactants and products in 200 µL are given for each measurement. The purpose of each measurement for the further normalisation and post-processing are outlined.

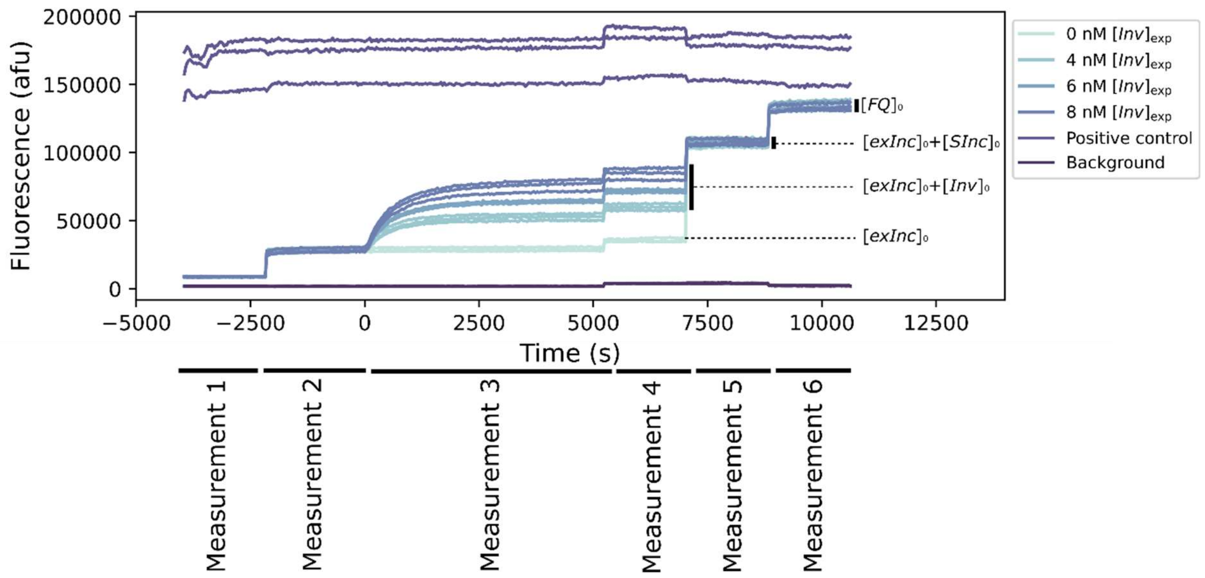

**Figure S8. Example of raw fluorescent traces for $DNA_{py}$>$RNA_{py}$ strand displacement experiment.** Raw fluorescence data from $DNA_{py}$>$RNA_{py}$ strand displacement reaction (15 nM L-d06z16 Rep-FQ + 10 nM L-y04b11 SIncR with 20% excess L-y00b11 IncR + 0-8 nM L-y10b26 InvD) to illustrate the fluorescence measurements and controls for each experiment. Measurement 1 captured the initial reporter *FQ* fluorescence. Measurement 2 monitored the fluorescence after injection of *SInc*. Measurement 3 captured the reaction kinetics after injection of *Inv*. Measurement 4 monitored the steady state fluorescence. Measurement 5 measured the fluorescence after an excess of *Inv* (~20 nM) is added. Measurement 6 monitored the fluorescence after an excess of *Inc* (~20 nM) is added. Each reaction is performed in triplicate. The positive control is composed of 15 nM *FInc* (AlexaFluor488-*Inc*) and the background negative control is 1 M NaCl + 1X TAE.

We experienced some difficulties in characterising the $DNA_{pu}$>$RNA_{pu}$ strand displacement reactions due to an undesired interaction between the *FQ* and *SInc* complexes which resulted in a significant leak reaction (Figures S19-20). This leak reaction appeared to be exaggerated by the longer 8nt reporter toehold. Consequently, for these $DNA_{pu}$>$RNA_{pu}$ strand displacement reactions we employed a reporter complex with a shorter 6nt reporter toehold. Moreover, by using higher concentrations of *Inv* we were able to better differentiate between the undesired leak reactions and the reactions of interest. As such, we used $[Inv]_{exp}$ = 40 nM, 60 nM and 80 nM for these reactions (full details in Table S11). This increased the rate of the desired strand displacement reactions which facilitated better estimation of the rate for these systems. However, these higher *Inv* concentrations meant that we were unable to explicitly infer the concentration of *Inv* by saturating the system (example traces shown in Figure S9).

| Steps | Total reaction volume (μL) | Addition | Expected concentration in 200 μL (nM) | Purpose |
|---|---|---|---|---|
| Measurement 1 | 140 | Reaction: 10 μL $[FQ]_{exp}$ = 300 nM + 130 μL of 1 M NaCl + 1X TAE | $[FQ]_{exp}$ = 15 | Estimate residual fluorescence baseline |
| | | Positive control: 3 μL $[F]_{exp}$ = 1 μM + 10 μL $[\mathrm{Inc}]_{exp}$ = 400 nM + 127 μL 1 M NaCl + 1X TAE | $[FInc]_{PCexp}$ = 15 | Reference fluorescence value |
| | | Negative control: 10 μL $[\mathrm{FQ}]_{exp}$ = 300 nM + 130 μL of 1 M NaCl + 1X TAE | $[FQ]_{NCexp}$ = 15; $[SInc]_{NCexp}$ = 10 | |
| | | Buffer-only control: 140 μL of 1 M NaCl + 1X TAE | | |
| Measurement 2 | 150 | Reaction: 10 μL of $[SInc]_{exp}$ = 200 nM | $[FQ]_{exp}$ = 13; $[SInc]_{exp}$ = 10; $[FInc]_{exp}$ = 2 | Estimate $[exInc]_0$ |
| | | Positive control: 10 μL of 1 M NaCl + 1X TAE | $[FInc]_{PCexp}$ = 15 | |
| | | Negative control: 10 μL of $[SInc]_{exp}$ = 200 nM | $[FQ]_{NCexp}$ = 13; $[SInc]_{NCexp}$ = 10; $[FInc]_{NCexp}$ = 2 | Estimate $[exInc]_{NC0}$ |
| | | Buffer-only control: 10 μL of 1 M NaCl + 1X TAE | | |
| Measurement 3 | 200 | Reaction: 16-32 μL of $[Inv]_{exp}$ = 500 nM + 34-18 μL | Evolve with time | Reaction kinetics |

| | | | | | |
|---|---|---|---|---|---|
| | | | of 1 M NaCl + 1X TAE | | |
| | | | Positive control:<br>50 µL of 1 M NaCl + 1X TAE | $[\textit{FInc}]_{PCexp}$ = 15 | |
| | | | Negative control:<br>50 µL of 1 M NaCl + 1X TAE | $[\textit{FQ}]_{NCexp}$ = 13;<br>$[\textit{SInc}]_{NCexp}$ = 10;<br>$[\textit{FInc}]_{NCexp}$ = 2 | |
| | | | Buffer-only control:<br>50 µL of 1 M NaCl + 1X TAE | | |
| Measurement 4 | | 200 | Reaction:<br>- | $[\textit{FQ}]_{exp}$ = 3;<br>$[\textit{SInv}]_{exp}$ = 10;<br>$[\textit{exInv}]_{exp}$ = 30-70;<br>$[\textit{FInc}]_{exp}$ = 12 | |
| | | | Positive control:<br>- | $[\textit{FInc}]_{PCexp}$ = 15 | |
| | | | Negative control:<br>- | $[\textit{FQ}]_{NCexp}$ = 13;<br>$[\textit{SInc}]_{NCexp}$ = 10;<br>$[\textit{FInc}]_{NCexp}$ = 2 | |
| | | | Buffer-only control:<br>- | | |
| Measurement 5 | 210 | | Reaction:<br>10 µL of $[\textit{Inv}]_{exp}$ = 400 nM | $[\textit{FQ}]_{exp}$ = 3;<br>$[\textit{SInv}]_{exp}$ = 10;<br>$[\textit{FInc}]_{exp}$ = 12 | Estimate $[\textit{SInc}]_0$ |
| | | | Positive control:<br>10 µL of 1 M NaCl + 1X TAE | $[\textit{FInc}]_{PCexp}$ = 15 | |
| | | | Negative control:<br>10 µL of $[\textit{Inv}]_{exp}$ = 400 nM | $[\textit{FQ}]_{NCexp}$ = 3;<br>$[\textit{SInv}]_{NCexp}$ = 10;<br>$[\textit{FInc}]_{NCexp}$ = 12 | |
| | | | Buffer-only control:<br>10 µL of 1 M NaCl + 1X TAE | | |
| Measurement 6 | 220 | | Reaction: | $[\textit{SInv}]_{exp}$ = 10; | Estimate $[\textit{FQ}]_0$ |

| | | | |
|---|---|---|---|
| | 10 μL of $[Inc]_{exp}$ = 400 nM | $[FInc]_{exp}$ = 15 | |
| | Positive control: 10 μL of 1 M NaCl + 1X TAE | $[FInc]_{PCexp}$ = 15 | |
| | Negative control: 10 μL of $[Inc]_{exp}$ = 400 nM | $[SInv]_{NCexp}$ = 10; $[FInc]_{NCexp}$ = 15 | Estimate $[FQ]_{NC0}$ |
| | Buffer-only control: 10 μL of 1M NaCl + 1X TAE | | |

**Table S11. Detailed experimental protocol for $DNA_{pu}$>$RNA_{pu}$ strand displacement reactions to estimate $k_{\mathbf{eff}}$.** Summary of volumes of reactants added at each measurement for full $DNA_{pu}$>$RNA_{pu}$ strand displacement reactions. *FQ* is added at an expected concentration of 15 nM and *SInc* is added at an expected concentration of 10 nM for all reactions and a series of different expected *Inv* (40-80 nM) concentrations are added to trigger initiation of the reaction. The expected concentration of reactants and products in 200 μL are given for each measurement. These expected concentrations are under the assumption that the leak reaction is negligible. The purpose of each measurement for the further normalisation and post-processing are outlined.

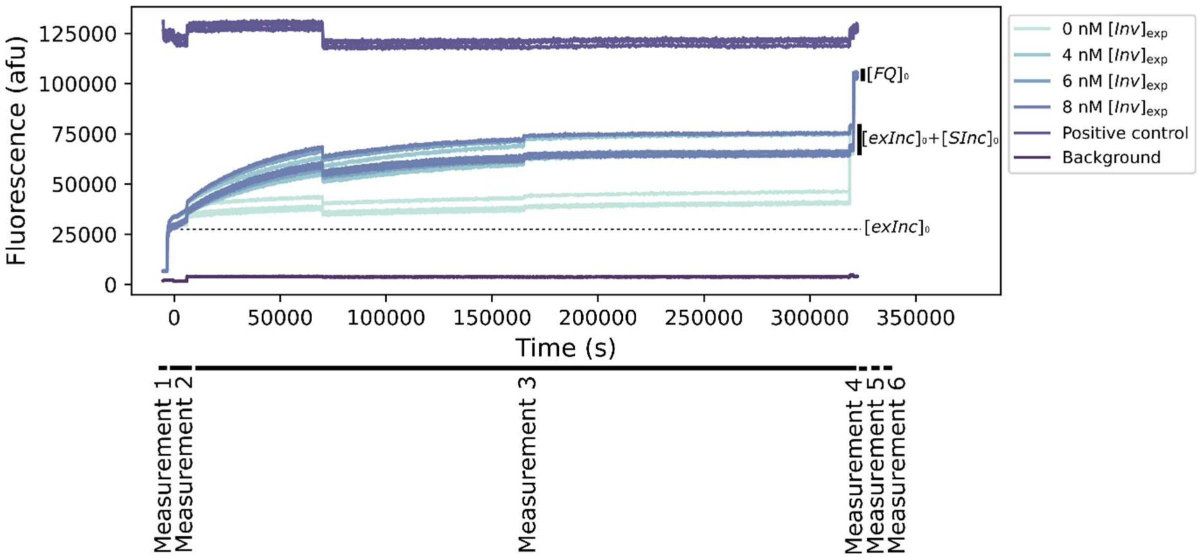

**Figure S9. Example of raw fluorescent traces for $DNA_{pu}$>$RNA_{pu}$ strand displacement experiment.** Raw fluorescence data from $DNA_{pu}$>$RNA_{pu}$ strand displacement reaction (15 nM H-d06z16 Rep-FQ + 10 nM H-y04b11 SIncR with 20% excess H-y00b11 IncR + 0-80 nM H-y06b21 InvD) to illustrate the fluorescence measurements and controls for each experiment. Measurement 1 captured the initial reporter *FQ* fluorescence. Measurement 2 monitored the fluorescence after injection of *SInc*. Measurement 3 captured the reaction kinetics after injection of *Inv*. We were unable to explicitly estimate the concentration of *Inv* added as *Inv* was in excess. Measurement 5 measured the fluorescence after a further excess of *Inv* (~20 nM) is added. Measurement 6 monitored the fluorescence after an excess of *Inc* (~20 nM) is added. Each reaction is performed in triplicate. The positive control is composed of 15 nM *FInc* (AlexaFluor488-*Inc*) and the background negative control is 1 M NaCl + 1X TAE.

*Supplementary note 8.2. Normalisation and data-processing of-full strand displacement experiments for $k_{\mathrm{eff}}$ estimation*

<u>Step 1: Removing background fluorescence produced by experimental buffer.</u>

Background autofluorescence is removed exactly as in Step 1 of Supplementary note 7.2.

<u>Step 2: Correction based on the positive control fluorescence</u>

The positive control is used to correct for environmentally-induced fluctuations as in Step 2 of Supplementary note 7.2 for RNA>DNA and DNA>DNA strand displacement reactions. For DNA>RNA strand displacement reactions with rapid kinetics we normalised by multiplying all data points by the ratio between the reference fluorescence value and the mean of the positive control fluorescence at steady state (measurement 4).

<u>Step 3: Estimation of $[FQ]_0$</u>

We calculate the mean fluorescence corresponding to complete dissociation of *FQ* for each reaction (measurement 5 for RNA>DNA and DNA>DNA strand displacement reactions ($[FInc]_{est(meas5)}$) and measurement 6 for DNA>RNA strand displacement ($[FInc]_{est(meas6)}$)). This value corresponds to the maximum fluorescence of the system where all *FQ* has been converted to the fluorescent product *FInc*. We estimate the initial concentration of *FQ* ($[FQ]_{est0}$) by dividing this mean fluorescence by $\beta$.

<u>Step 4: Conversion of fluorescence data to concentrations using $\alpha$ and $\beta$</u>

We employ the values of $\alpha$ and $\beta$ from the corresponding calibration curve in order to convert all data points according to equation (*S33*).

<u>Step 5: Obtain final reaction dynamics by removing $[exInc]_{NCest}$</u>

To obtain the final reaction dynamics we subtract the contribution from the excess *Inc* at each time point. We assume that the reacted excess *Inc* at each time point (t) in the negative control reaction ($[FInc]_{NCest(t)}$) is equivalent to that in the reactions of interest. We apply the same approach to convert the negative control signal to inferred concentrations. In this case we use the mean fluorescence corresponding to the complete dissociation of *FQ* ($[FQ]_{NCest0}$) for the negative control reaction (measurement 5 for RNA>DNA and DNA>DNA (Table S9) and measurement 6 for DNA>RNA (Table S10-S11)). We then apply equation (*S33*) as above. In this way we can determine the concentration of excess *Inc* that has reacted with the reporter over time.

To obtain our fully normalised displacement dynamics we subtract estimated concentration of excess *Inc* of the negative control ($[FInc]_{NCest(t)}$ at each timepoint (t) from the estimated concentration values for the reactions of interest at the corresponding timepoints. This subtraction relies on the assumption that the reaction between *exInc* and *FQ* is effectively instantaneous compared to the reaction between *Inv* and *SInc*. We perform this subtraction for all timepoints during both the reactions kinetics and the steady state (measurement 2 and measurement 3 for DNA>DNA and RNA>DNA strand displacement and measurement 3 and measurement 4 for DNA>RNA strand displacement) Mathematically,

$$[FInc]_{\mathrm{estrct(t)}} = [FInc]_{\mathrm{est(t)}} - [FInc]_{\mathrm{NCest(t)}}, \qquad (S37)$$

where $[FInc]_{estrct(t)}$ is the estimated concentration in nM of [*FInc*] that arises from the displacement dynamics of interest, rather than the reaction of the excess *Inc* at each time point (t) ($[FInc]_{NCest(t)}$). Using this strategy we obtained the processed displacement dynamics in Figures S10, S11 and S12A-H.

<u>Step 6: Estimation of $[exInc]_{NCest0}$, $[Inv]_{est0}$, $[SInc]_{est0}$ and $[FQ]_{rctest0}$</u>

We can calculate the estimated initial (t = 0 s) and reactive concentrations of *exInc*, *SInc* and *FQ* complexes that are present in each reaction in order to explicitly account for these concentrations in the fitting protocol. We take a similar approach to step 5 in order to obtain $[exInc]_{NCest0}$. However, rather than calculating $[FInc]_{NCest(t)}$ at each timepoint (t) we obtain a mean fluorescence over steady state (measurement 3 for DNA>DNA and RNA>DNA strand displacement ($[FInc]_{NCest(meas3)}$) (Table S9) and measurement 4 for DNA>RNA strand displacement ($[FInc]_{NCest(meas4)}$) (Table S10-S11)) for the negative control. We then plug this mean fluorescence into equation (*S33*) using the relevant value of $[FQ]_{est0}$ to obtain $[FInc]_{NCest}$ during this measurement, which corresponds to $[exInc]_{NCest0}$.

To estimate the initial concentration of *Inv* we subtract $[exInc]_{NCest0}$ from the mean fluorescence at steady state (measurement 3 for DNA>DNA and RNA>DNA strand displacement ($[FInc]_{est(meas3)}$) and measurement 4 for DNA>RNA strand displacement ($[FInc]_{est(meas4)}$) according to

$$[Inv]_{\mathrm{est0}} = [FInc]_{\mathrm{est(meas3|meas4)}} - [exInc]_{\mathrm{NCest}} \ . \qquad (S38)$$

For *SInc* we calculate the mean fluorescence of measurement 4 for DNA>DNA and RNA>DNA strand displacement ($[FInc]_{est(meas4)}$) and measurement 5 for DNA>RNA strand displacement ($[FInc]_{est(meas5)}$) for each reaction of interest and again apply equation (*S33*). Finally, we subtract $[exInc]_{NCest0}$ in order to determine the estimated concentration *SInc* according to

$$[SInc]_{\mathrm{est0}} = [FInc]_{\mathrm{est(meas4|meas5)}} - [exInc]_{\mathrm{NCes}} \ . \qquad (S39)$$

To determine the estimated concentration of *FQ* that was actually available for reaction with *Inv* at the start of each reaction ($[FQ]_{rctest0}$), we subtract $[exInc]_{NCest0}$ from the $[FQ]_{est0}$ calculated in step 3. This subtraction relies on the assumption that *exInc* reacts effectively instantaneously with *FQ* compared to the reaction between *Inv* and *SInc*. We calculate $[FQ]_{rctest0}$ according to

$$[FQ]_{\mathrm{estrct0}} = [FQ]_{\mathrm{est0}} - [exInc]_{\mathrm{NCest}} \ . \qquad (S40)$$

We had to adapt this normalisation and post-processing strategy slightly for $DNA_{pu}$>$RNA_{pu}$ strand displacement reactions due to the undesired leak reaction between *SInc* and *FQ* complexes. Due to the increase in the fluorescence over time resulting from an off-target interaction between *SInc* and *FQ*, we were not able to simply subtract $[FInc]_{NCest}$ at each time point in order to obtain the reaction dynamics as in equation (*S37*) (Figures S12I-K). Instead we used the mean fluorescence of the negative control in measurement 2 to estimate $[exInc]_{NCest0}$ for these reactions and subtracted the fluorescence due to $[exInc]_{NCest0}$ from the inferred concentrations at each time point. We then accounted for both the desired strand displacement and undesired leak reactions in the ODE fitting (supplementary note 8.4). Our estimate of $[exInc]_{est0}$ for these reactions allowed us to calculate the concentrations *SInc* and *FQ* that were present to react with *Inv* ($[SInc]_{est0}$ and $[FQ]_{rctest0}$). We were unable to calculate $[Inv]_{est0}$ as an excess of *Inv* was used. As such, we set $[Inv]_{est0}$ at 40, 60 and 80 nM.

Processed data are shown in Figures S10-S12.

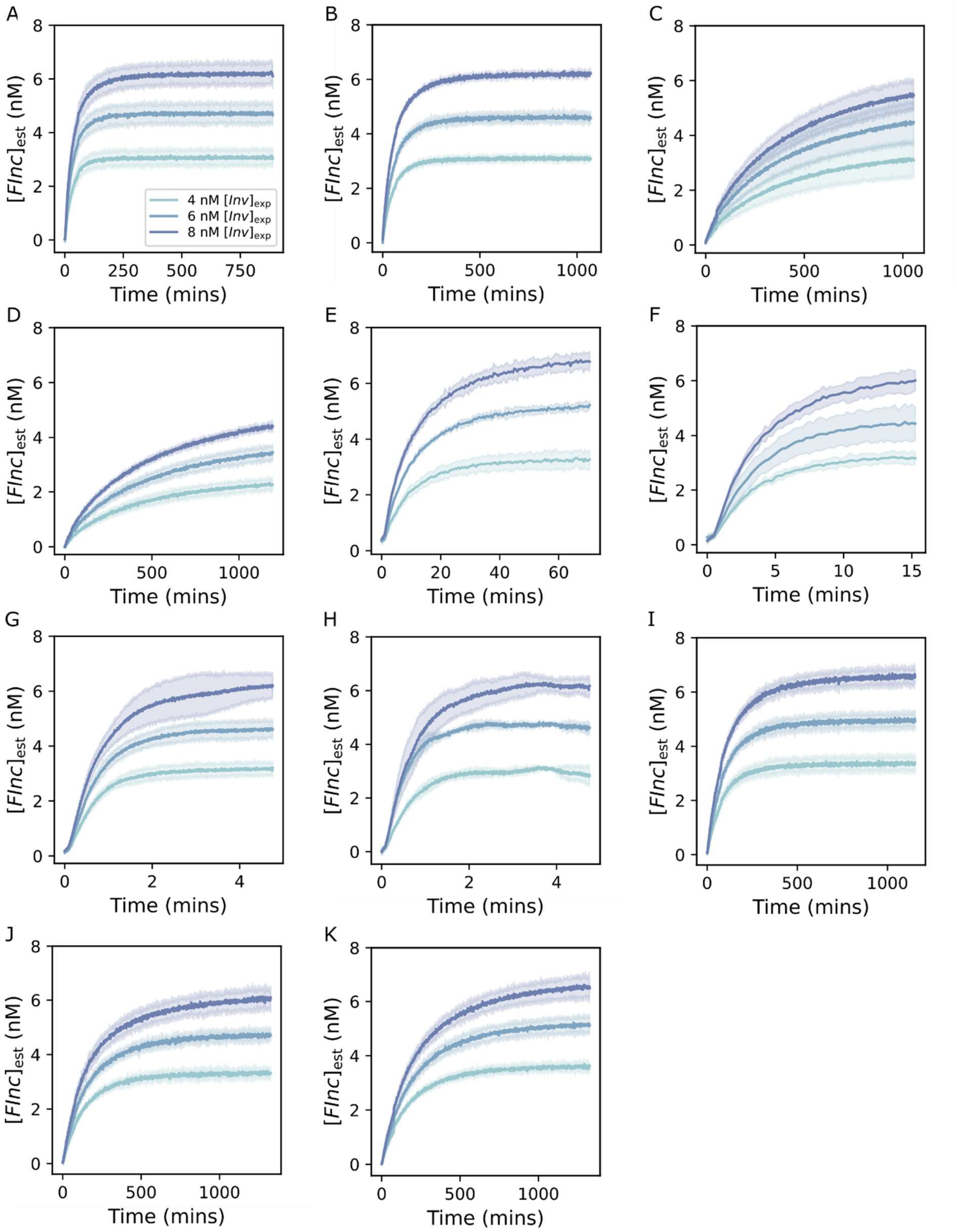

**Figure S10. Processed fluorescent traces for full DNA>DNA strand displacement reactions for $k_{\text{eff}}$ estimation. Processed** fluorescent outputs for DNA>DNA strand displacement reactions in which reporter ($[FQ]_{\text{exp}}$ = 15 nM) is combined with of incumbent-substrate complex ($[SInc]_{\text{exp}}$ = 10 nM) and invader strand ($[Inv]_{\text{exp}}$ = 4-8 nM). Each reaction is performed in triplicate. Data is processed to convert fluorescence readings to an estimated concentration of the fluorescent product, *FInc*, in nM and to account for fluctuations due to changes in temperature and reaction volume. For $DNA_{py}$>$DNA_{py}$ 4 different branch migration domain lengths (11-26nt) and 5 different invader toehold lengths (4-10nt)

were tested. For $DNA_{pu}$>$DNA_{pu}$ 3 different branch migration domain lengths (11-21nt) were tested. This normalised fluorescence data is used to estimate $k_{\text{eff}}$ for each DNA>DNA toehold exchange reaction. A) L-y10b26 InvD displacement of L-y04b11 SIncD with L-d06z16 Rep-FQ. B) L-y10b26 InvD displacement of L-y04b16 SIncD with L-d06z16 Rep-FQ. C) L-y10b26 InvD displacement of L-y04b21 SIncD with L-d06z16 Rep-FQ. D) L-y10b26 InvD displacement of L-y04b26 SIncD with L-d06z16 Rep-FQ. E) L-y10b26 InvD displacement of L-y05b26 SIncD with L-d06z16 Rep-FQ. F) L-y10b26 InvD displacement of L-y06b26 SIncD with L-d06z16 Rep-FQ. G) L-y10b26 InvD displacement of L-y07b26 SIncD with L-d06z16 Rep-FQ. H) L-y10b26 InvD displacement of L-y10b26 SIncD with L-d06z16 Rep-FQ. I) H-y06b21 InvD displacement of H-y04b11 SIncD with H-d08z16 Rep-FQ. J) H-y06b21 InvD displacement of H-y04b16 SIncD with H-d08z16 Rep-FQ. K) H-y06b21 InvD displacement of H-y04b21 SIncD with H-d08z16 Rep-FQ. The legend given in A corresponds to all subplots.

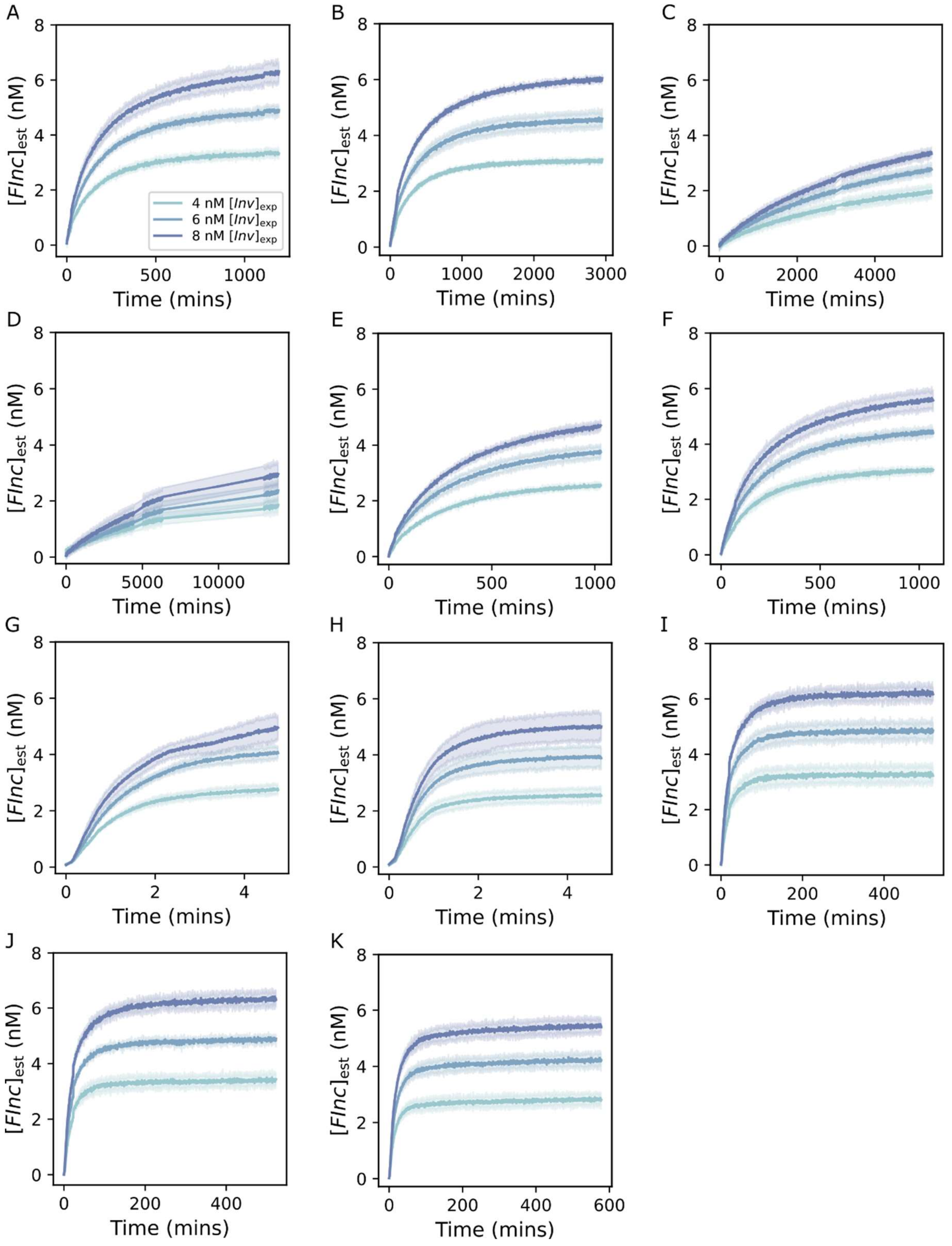

**Figure S11. Processed fluorescent traces for full RNA>DNA strand displacement reactions for $k_{\mathrm{eff}}$ estimation.** Processed fluorescent outputs for RNA>DNA strand displacement reactions in which reporter ($[FQ]_{exp}$ = 15 nM) is combined with of incumbent-substrate complex ($[SInc]_{exp}$ = 10 nM) and invader strand ($[Inv]_{exp}$ = 4-8 nM). Each reaction is performed in triplicate. Data is processed to convert fluorescence readings to an estimated concentration of the fluorescent product, *FInc*, in nM and to account for fluctuations due to changes in temperature and reaction volume. For $RNA_{py}$>$DNA_{py}$ 4 different branch migration domain lengths (11-26nt) and 5 different invader toehold lengths (4-10nt)

were tested. For $RNA_{pu}$>$DNA_{pu}$ 3 different branch migration domain lengths (11-21nt) were tested. This normalised fluorescence data is used to estimate $k_{\mathrm{eff}}$ for each RNA>DNA toehold exchange reaction. A) L-y10b26 InvR displacement of L-y04b11 SIncD with L-d06z16 Rep-FQ. B) L-y10b26 InvR displacement of L-y04b16 SIncD with L-d06z16 Rep-FQ. C) L-y10b26 InvR displacement of L-y04b21 SIncD with L-d06z16 Rep-FQ. D) L-y10b26 InvR displacement of L-y04b26 SIncD with L-d06z16 Rep-FQ. E) L-y10b26 InvR displacement of L-y05b26 SIncD with L-d06z16 Rep-FQ. F) L-y10b26 InvR displacement of L-y06b26 SIncD with L-d06z16 Rep-FQ. G) L-y10b26 InvR displacement of L-y07b26 SIncD with L-d06z16 Rep-FQ. H) L-y10b26 InvR displacement of L-y10b26 SIncD with L-d06z16 Rep-FQ. I) H-y06b21 InvR displacement of H-y04b11 SIncD with H-d08z16 Rep-FQ. J) H-y06b21 InvR displacement of H-y04b16 SIncD with H-d08z16 Rep-FQ. K) H-y06b21 InvR displacement of H-y04b21 SIncD with H-d08z16 Rep-FQ. The legend given in A corresponds to all subplots.

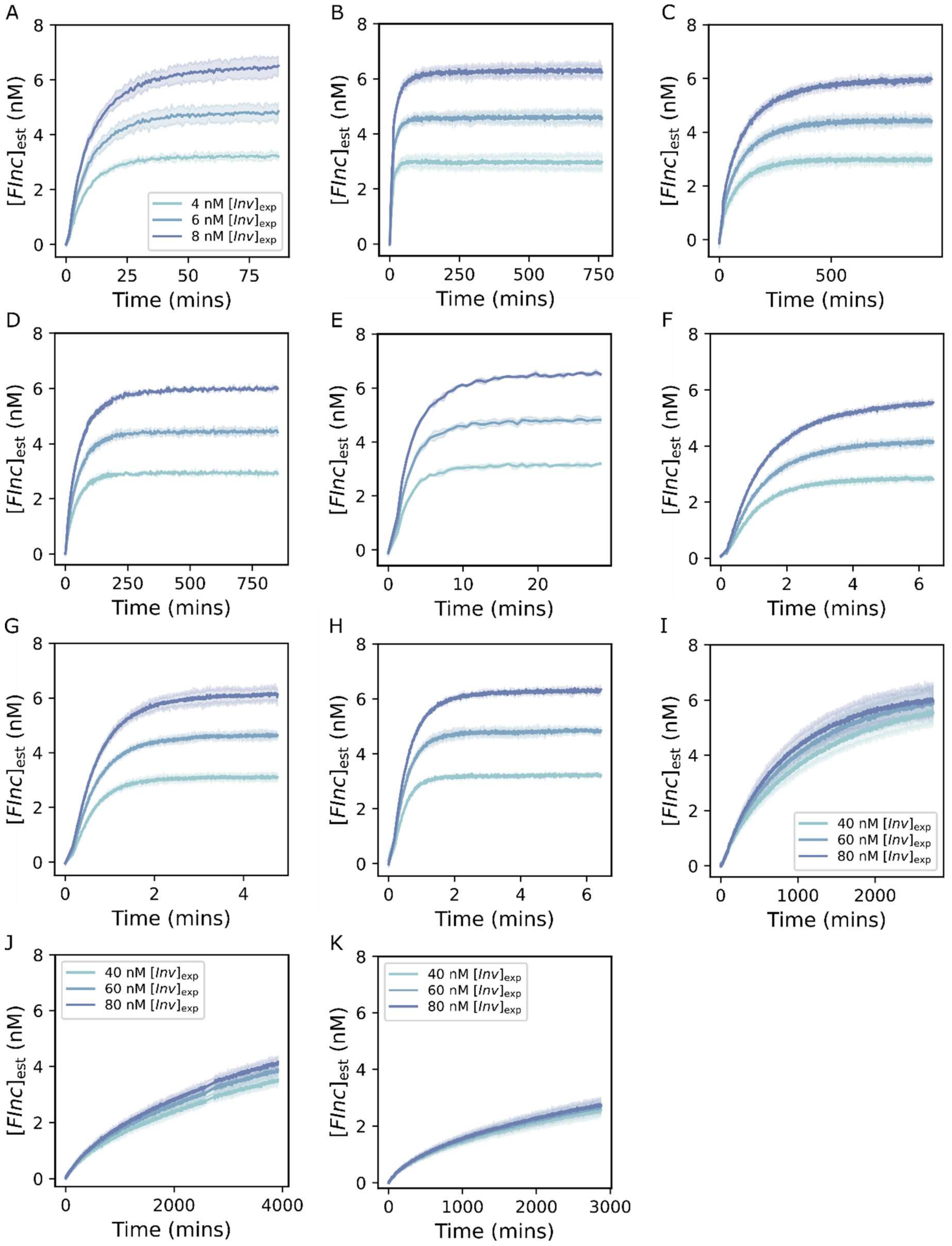

**Figure S12. Processed fluorescent traces for full DNA>RNA strand displacement reactions for $k_{\mathrm{eff}}$ estimation.** Processed fluorescent outputs for DNA>RNA strand displacement reactions in which reporter ($[FQ]_{\mathrm{exp}}$ = 15 nM) is combined with incumbent-substrate complex ($[SInc]_{\mathrm{exp}}$ = 10 nM) and an invader strand ($[Inv]_{\mathrm{exp}}$ = 4-8 nM (or 40-80 nM for $\mathrm{DNA_{pu}}$>$\mathrm{RNA_{pu}}$)). Each reaction is performed in triplicate. Data is processed to convert fluorescence readings to an estimated concentration of the fluorescent product, *FInc*, in nM and to account for fluctuations due to changes in temperature and reaction volume. For $\mathrm{DNA_{py}}$>$\mathrm{RNA_{py}}$ 4 different branch migration domain lengths (11-26nt) and 5 different invader toehold lengths (4-10nt) were tested. For $DNA_{pu}$>$RNA_{pu}$ 3 different branch migration domain lengths (11-21nt) were tested. This normalised fluorescence data is used to estimate $k_{\mathrm{eff}}$ for each DNA>RNA toehold exchange reaction. A) L-y10b26 InvD displacement of L-y04b11 SIncR with L-d06z16 Rep-FQ. B) L-y10b26 InvD displacement of L-y04b16 SIncR with L-d06z16 Rep-FQ. C) L-y10b26 InvD displacement of L-y04b21 SIncR with L-d06z16 Rep-FQ. D) L-y10b26 InvD displacement of L-y04b26 SIncR with L-d06z16 Rep-FQ. E) L-y10b26 InvD displacement of L-y05b26 SIncR with L-d06z16 Rep-FQ. F) L-y10b26 InvD displacement of L-y06b26 SIncR with L-d06z16 Rep-FQ. G) L-y10b26 InvD displacement of L-y07b26 SIncR with L-d06z16 Rep-FQ. H) L-y10b26 InvD displacement of L-y10b26 SIncR with L-d06z16 Rep-FQ. I) H-y06b21 InvD displacement of H-y04b11 SIncR with H-d06z16 Rep-FQ. J) H-y06b21 InvD displacement of H-y04b16 SIncR with H-d06z16 Rep-FQ. K) H-y06b21 InvD displacement of H-y04b21 SIncR with H-d06z16 Rep-FQ. The legend given in A corresponds to subplots B-H.

*Supplementary note 8.3. Fitting ODEs to full strand displacement experiments for $k_{\mathrm{eff}}$ estimation*

We present two alternative approaches to fitting $k_{\mathrm{eff}}$: individual and global fits. Individual fitting obtains fits and estimates $k_{\mathrm{eff}}$ for each kinetic trace for a given set of strands, while global fitting yields a single $k_{\mathrm{eff}}$ estimate across all replicates and $[Inv]_{\mathrm{exp}}$ for a given set of strands. In the main text we report only the $k_{\mathrm{eff}}$ estimates from global fits (with the exception of $DNA_{pu}$>$RNA_{pu}$ reactions in which we use the individual fits), however we present both here as evidence that our $k_{\mathrm{eff}}$ estimates are robust to the fitting approach.

We used a similar approach to the reporter characterisations (supplementary note 7.3) to integrate the system of ODEs for full strand displacement reactions (equations (*4-6*)). In this case for the individual fits we used jax.experimental.ode.odeint. We set initial, fixed concentrations of 0 nM for [*SInv*], [*Inc*], [*FInc*], [*Q*] and used the estimated concentrations of *SInc* ($[SInc]_{\mathrm{est0}}$) and *FQ* ($[FQ]_{\mathrm{rctest0}}$) as initial concentrations. We included $k_{\mathrm{rep}}$, $k_{\mathrm{eff}}$ and $[Inv]_{\mathrm{fit0}}$ as individual arguments in this function. For both $k_{\mathrm{rep}}$ and $k_{\mathrm{eff}}$ we input a log-transformed value to given that both $k_{\mathrm{rep}}$ and $k_{\mathrm{eff}}$ vary over orders of magnitude. We used a fixed value for $k_{\mathrm{rep}}$ from the corresponding reporter characterisation (global jackknife estimate) while we fitted $k_{\mathrm{eff}}$ and $[Inv]_0$. We next apply scipy.optimize.curve_fit as in supplementary note 7.3 in order to fit the solution of the numerical integration to the experimental data by varying $[Inv]_{\mathrm{fit0}}$ and $k_{\mathrm{eff}}$ for each individual $[Inv]_{\mathrm{exp}}$ and replicate. The inputs to this function are the ODE solver object, the time steps over which the measurement is taken, the normalised fluorescence measurements and the initial predictions and lower and upper bounds for the parameters to be estimated. For $[Inv]_{\mathrm{fit0}}$ we used an initial prediction of the steady state estimates from experimental fluorescence data ($[Inv]_{\mathrm{est0}}$). The lower and upper bound were set to $[Inv]_{\mathrm{est0}}/2$ and $2[Inv]_{\mathrm{est0}}$, respectively. For $k_{\mathrm{eff}}$ the lower and upper limits were set at $10^1$ $M^{-1}$ $s^{-1}$ and $10^8$ $M^{-1}$ $s^{-1}$, respectively. The initial $k_{\mathrm{eff}}$ prediction differed for each system depending on the length of the invader toehold and branch migration domain. We calculated the mean $k_{\mathrm{eff}}$ from the inverse log-transformed $k_{\mathrm{eff}}$ values across all 3 replicates at 3 different $[Inv]_{\mathrm{exp}}$ for each set of strands and estimated a standard error based on the variability in these individual fits (numerical values in Table S12, fits shown in Figure S13, Figure S14 and Figure S15).

| Full strand displacement system | Mean $k_{\text{eff}}$ (M-1 s-1) | Standard error (M-1 s-1) |
| --- | --- | --- |
| L-y10b26 InvD + L-y04b11 SIncD + L-d06z16 Rep-FQ | $5.749 \cdot 10^4$ | $0.105 \cdot 10^4$ |
| L-y10b26 InvD + L-y04b16 SIncD + L-d06z16 Rep-FQ | $2.938 \cdot 10^4$ | $0.075 \cdot 10^4$ |
| L-y10b26 InvD + L-y04b21 SIncD + L-d06z16 Rep-FQ | $5.865 \cdot 10^3$ | $0.119 \cdot 10^3$ |
| L-y10b26 InvD + L-y04b26 SIncD + L-d06z16 Rep-FQ | $4.972 \cdot 10^3$ | $0.162 \cdot 10^3$ |
| L-y10b26 InvD + L-y05b26 SIncD + L-d06z16 Rep-FQ | $1.908 \cdot 10^5$ | $0.064 \cdot 10^5$ |
| L-y10b26 InvD + L-y06b26 SIncD + L-d06z16 Rep-FQ | $6.112 \cdot 10^5$ | $0.581 \cdot 10^5$ |
| L-y10b26 InvD + L-y07b26 SIncD + L-d06z16 Rep-FQ | $4.434 \cdot 10^6$ | $0.369 \cdot 10^6$ |
| L-y10b26 InvD + L-y10b26 SIncD + L-d06z16 Rep-FQ | $7.249 \cdot 10^6$ | $2.573 \cdot 10^6$ |
| L-y10b26 InvR + L-y04b11 SIncD + L-d06z16 Rep-FQ | $1.223 \cdot 10^4$ | $0.041 \cdot 10^4$ |
| L-y10b26 InvR + L-y04b16 SIncD + L-d06z16 Rep-FQ | $6.471 \cdot 10^3$ | $0.237 \cdot 10^3$ |
| L-y10b26 InvR + L-y04b21 SIncD + L-d06z16 Rep-FQ | $5.533 \cdot 10^2$ | $0.891 \cdot 10^2$ |
| L-y10b26 InvR + L-y04b26 SIncD + L-d06z16 Rep-FQ | $3.625 \cdot 10^2$ | $1.368 \cdot 10^2$ |
| L-y10b26 InvR + L-y05b26 SIncD + L-d06z16 Rep-FQ | $7.266 \cdot 10^3$ | $0.242 \cdot 10^3$ |
| L-y10b26 InvR + L-y06b26 SIncD + L-d06z16 Rep-FQ | $1.200 \cdot 10^4$ | $0.075 \cdot 10^4$ |
| L-y10b26 InvR + L-y07b26 SIncD + L-d06z16 Rep-FQ | $2.198 \cdot 10^6$ | $0.285 \cdot 10^6$ |
| L-y10b26 InvR + L-y10b26 SIncD + L-d06z16 Rep-FQ | $4.793 \cdot 10^6$ | $0.541 \cdot 10^6$ |
| L-y10b26 InvD + L-y04b11 SIncR + L-d06z16 Rep-FQ | $2.169 \cdot 10^5$ | $0.048 \cdot 10^5$ |
| L-y10b26 InvD + L-y04b16 SIncR + L-d06z16 Rep-FQ | $1.865 \cdot 10^5$ | $0.223 \cdot 10^5$ |
| L-y10b26 InvD + L-y04b21 SIncR + L-d06z16 Rep-FQ | $2.521 \cdot 10^4$ | $0.178 \cdot 10^4$ |
| L-y10b26 InvD + L-y04b26 SIncR + L-d06z16 Rep-FQ | $4.125 \cdot 10^4$ | $0.154 \cdot 10^4$ |
| L-y10b26 InvD + L-y05b26 SIncR + L-d06z16 Rep-FQ | $7.888 \cdot 10^5$ | $0.296 \cdot 10^5$ |
| L-y10b26 InvD + L-y06b26 SIncR + L-d06z16 Rep-FQ | $3.037 \cdot 10^6$ | $0.070 \cdot 10^6$ |

| | | |
|---|---|---|
| L-y10b26 InvD + L-y07b26 SIncR + L-d06z16 Rep-FQ | $5.073 \cdot 10^6$ | $0.186 \cdot 10^6$ |
| L-y10b26 InvD + L-y10b26 SIncR + L-d06z16 Rep-FQ | $1.359 \cdot 10^7$ | $0.144 \cdot 10^7$ |
| H-y06b21 InvD + H-y04b11 SIncD + H-d08z16 Rep-FQ | $1.932 \cdot 10^4$ | $0.163 \cdot 10^4$ |
| H-y06b21 InvD + H-y04b16 SIncD + H-d08z16 Rep-FQ | $1.772 \cdot 10^4$ | $0.120 \cdot 10^4$ |
| H-y06b21 InvD + H-y04b21 SIncD + H-d08z16 Rep-FQ | $1.157 \cdot 10^4$ | $0.084 \cdot 10^4$ |
| H-y06b21 InvR + H-y04b11 SIncD + H-d08z16 Rep-FQ | $1.021 \cdot 10^5$ | $0.012 \cdot 10^5$ |
| H-y06b21 InvR + H-y04b16 SIncD + H-d08z16 Rep-FQ | $1.176 \cdot 10^5$ | $0.039 \cdot 10^5$ |
| H-y06b21 InvR + H-y04b21 SIncD + H-d08z16 Rep-FQ | $2.633 \cdot 10^5$ | $0.475 \cdot 10^5$ |

**Table S12. $k_{\text{eff}}$ estimates from averaging over individual full strand displacement reactions.** Mean $k_{\text{eff}}$ estimates and standard errors from individual fits of 3 replicates at 3 different expected *Inv* ($[Inv]_{\text{exp}}$ = 4-8 nM) concentrations for RNA>DNA, DNA>DNA and DNA>RNA strand displacement reactions. Expected reporter ($[FQ]_{\text{exp}}$) and incumbent-substrate complex ($[SInc]_{\text{exp}}$) concentrations of 15 nM and 10 nM were used for all experiments, respectively. $k_{\text{eff}}$ estimates were obtained by fitting an ODE model of the full strand displacement reaction to each individual fluorescence curve.

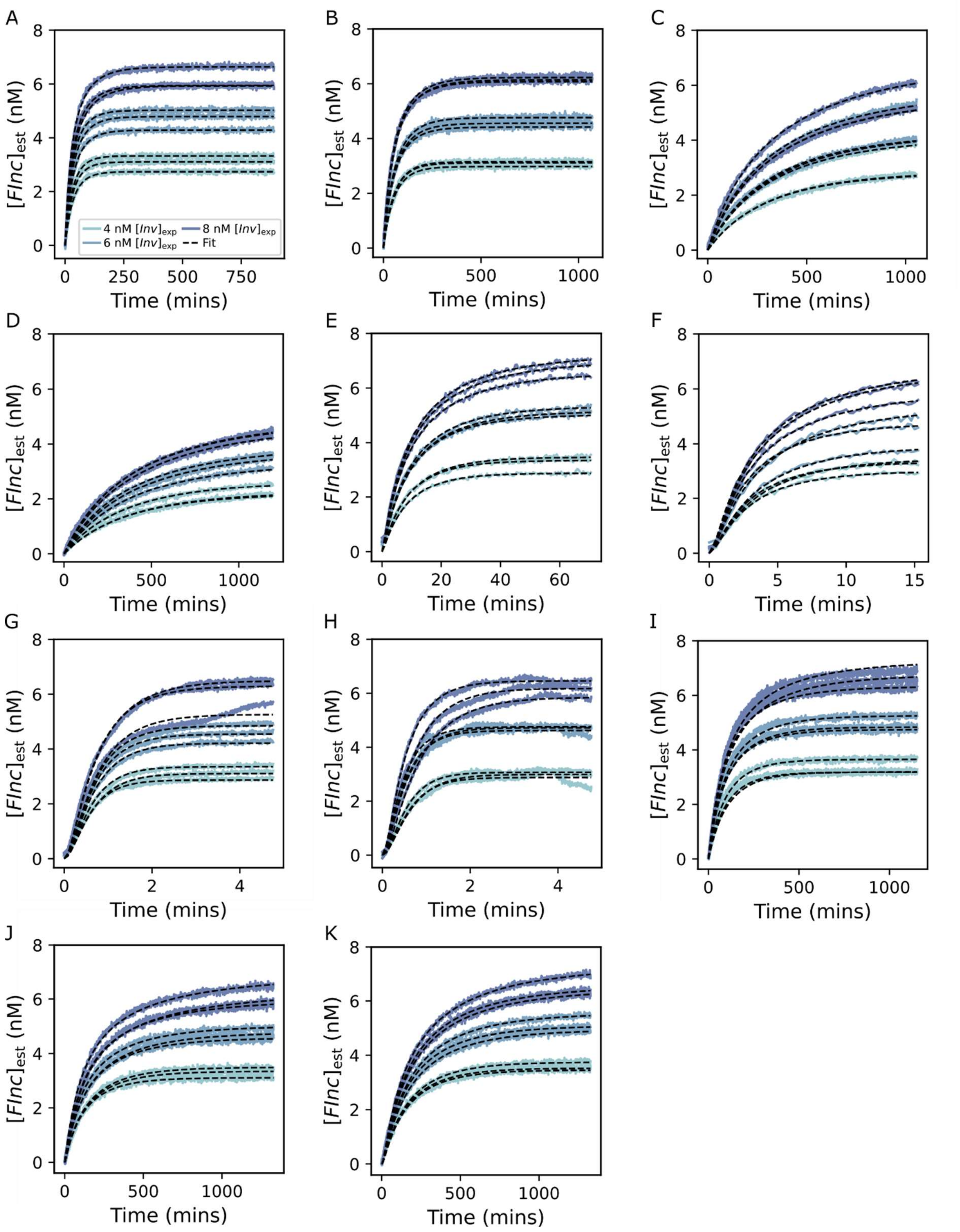

**Figure S13. Individual fits to fluorescent traces in full DNA>DNA strand displacement reactions.** Individual fits of the ODE model to normalised fluorescent data for each DNA>DNA strand displacement experiment. Each experiment was performed in triplicate and across 3 different expected *Inv* ($[Inv]_{exp}$ = 4-8 nM) concentrations. Expected reporter ($[FQ]_{exp}$) and incumbent-substrate complex ($[SInc]_{exp}$) concentrations of 15 nM and 10 nM were used for all experiments, respectively. Black, dotted lines represent fit of ODE model and solid, coloured lines represent the normalised experimental fluorescent data. A) L-y10b26 InvD displacement of L-y04b11 SIncD with L-d06z16 Rep-

FQ. B) L-y10b26 InvD displacement of L-y04b16 SIncD with L-d06z16 Rep-FQ. C) L-y10b26 InvD displacement of L-y04b21 SIncD with L-d06z16 Rep-FQ. D) L-y10b26 InvD displacement of L-y04b26 SIncD with L-d06z16 Rep-FQ. E) L-y10b26 InvD displacement of L-y05b26 SIncD with L-d06z16 Rep-FQ. F) L-y10b26 InvD displacement of L-y06b26 SIncD with L-d06z16 Rep-FQ. G) L-y10b26 InvD displacement of L-y07b26 SIncD with L-d06z16 Rep-FQ. H) L-y10b26 InvD displacement of L-y10b26 SIncD with L-d06z16 Rep-FQ. I) H-y06b21 InvD displacement of H-y04b11 SIncD with H-d08z16 Rep-FQ. J) H-y06b21 InvD displacement of H-y04b16 SIncD with H-d08z16 Rep-FQ. K) H-y06b21 InvD displacement of H-y04b21 SIncD with H-d08z16 Rep-FQ. The legend given in A corresponds to all subplots.

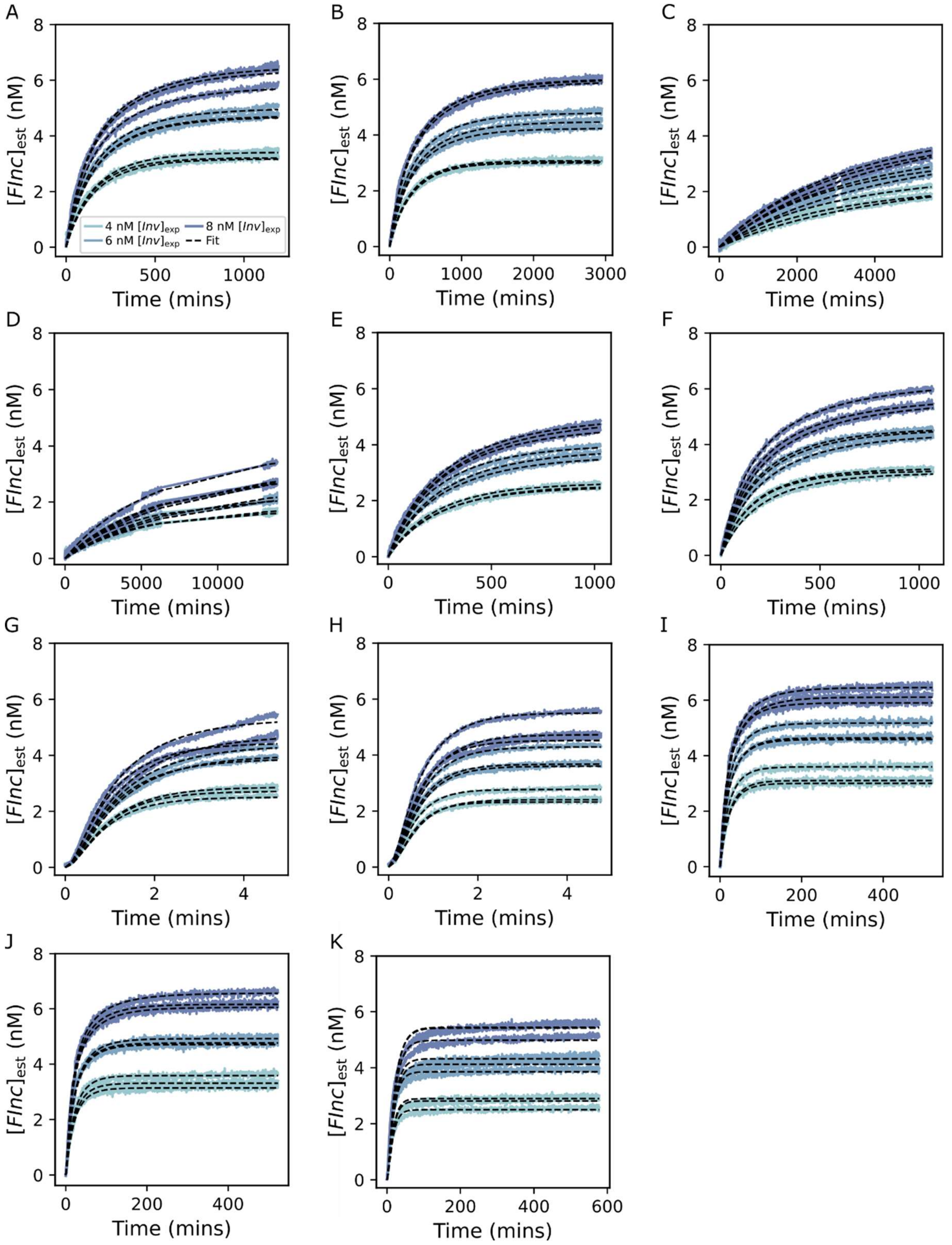

**Figure S14. Individual fits to fluorescent traces in full RNA>DNA strand displacement reactions.** Individual fits of the ODE model to normalised fluorescent data for each RNA>DNA strand displacement experiment. Each experiment was performed in triplicate and across 3 different expected *Inv* ($[Inv]_{exp}$ = 4-8 nM) concentrations. Expected reporter ($[FQ]_{exp}$) and incumbent-substrate complex ($[SInc]_{exp}$) concentrations of 15 nM and 10 nM were used for all experiments, respectively. Black, dotted lines represent fit of ODE model and solid, coloured lines represent the normalised experimental fluorescent data. A) L-y10b26 InvR displacement of L-y04b11 SIncD with L-d06z16 Rep-

FQ. B) L-y10b26 InvR displacement of L-y04b16 SIncD with L-d06z16 Rep-FQ. C) L-y10b26 InvR displacement of L-y04b21 SIncD with L-d06z16 Rep-FQ. D) L-y10b26 InvR displacement of L-y04b26 SIncD with L-d06z16 Rep-FQ. E) L-y10b26 InvR displacement of L-y05b26 SIncD with L-d06z16 Rep-FQ. F) L-y10b26 InvR displacement of L-y06b26 SIncD with L-d06z16 Rep-FQ. G) L-y10b26 InvR displacement of L-y07b26 SIncD with L-d06z16 Rep-FQ. H) L-y10b26 InvR displacement of L-y10b26 SIncD with L-d06z16 Rep-FQ. I) H-y06b21 InvR displacement of H-y04b11 SIncD with H-d08z16 Rep-FQ. J) H-y06b21 InvR displacement of H-y04b16 SIncD with H-d08z16 Rep-FQ. K) H-y06b21 InvR displacement of H-y04b21 SIncD with H-d08z16 Rep-FQ. The legend given in A corresponds to all subplots.

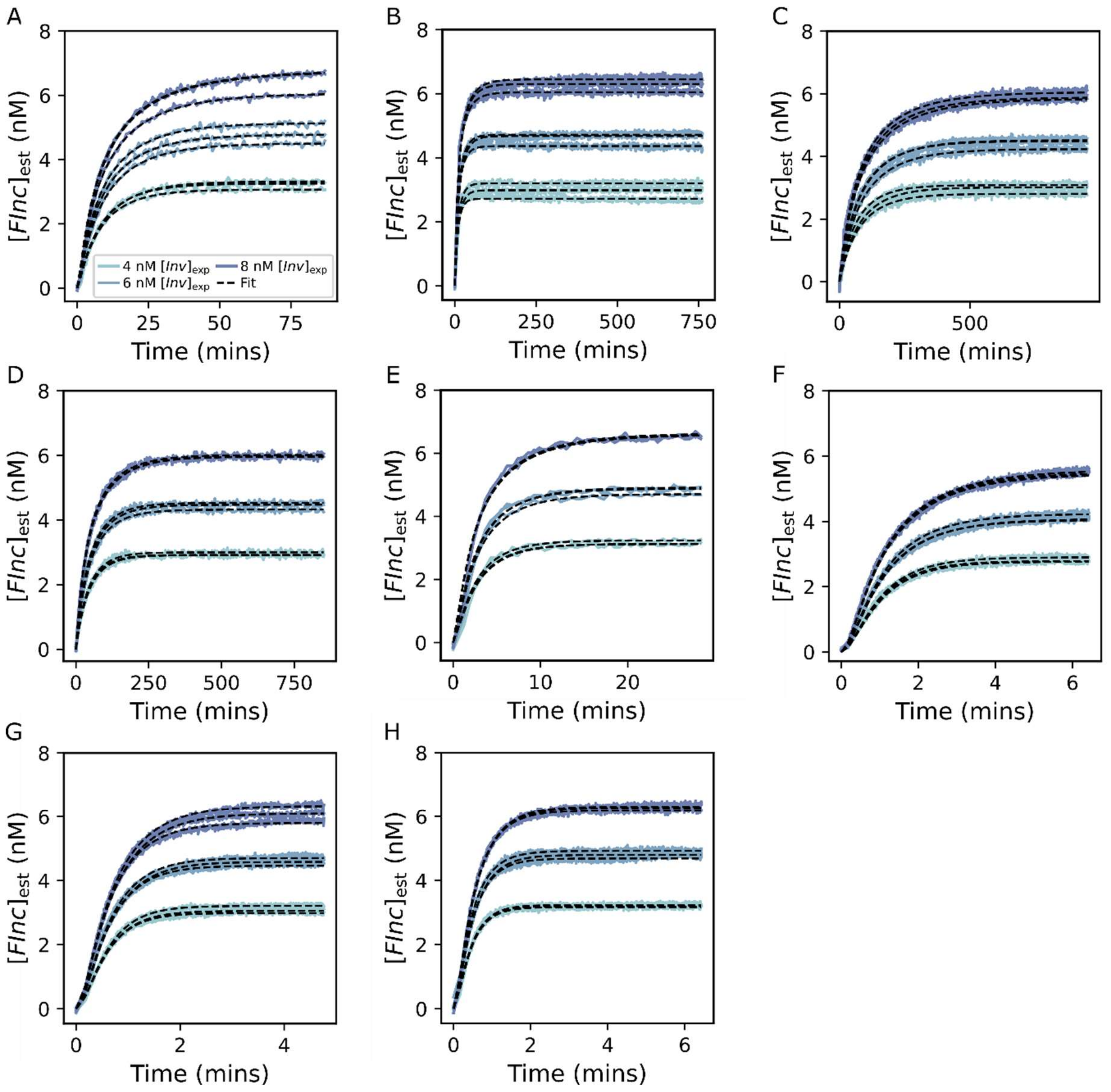

**Figure S15. Individual fits to fluorescent traces in full DNA>RNA strand displacement reactions.** Individual fits of the ODE model to normalised fluorescent data for each DNA>RNA strand displacement experiment. Each experiment was performed in triplicate and across 3 different expected *Inv* ($[Inv]_{exp}$ = 4-8 nM) concentrations. Expected reporter ($[FQ]_{exp}$) and incumbent-substrate complex ($[SInc]_{exp}$) concentrations of 15 nM and 10 nM were used for all experiments, respectively. Black, dotted lines represent fit of ODE model and solid, coloured lines represent the normalised experimental fluorescent data. A) L-y10b26- InvD displacement of L-y04b11 SIncR with L-d06z16 Rep-FQ. B) L-y10b26 InvD displacement of L-y04b16 SIncR with L-d06z16 Rep-FQ. C) L-y10b26 InvD displacement of L-y04b21 SIncR with L-d06z16 Rep-FQ. D) L-y10b26 InvD displacement of L-y04b26 SIncR with L-d06z16 Rep-FQ. E) L-y10b26 InvD displacement of L-y05b26 SIncR with L-d06z16 Rep-FQ. F) L-y10b26 InvD displacement of L-y06b26 SIncR with L-d06z16 Rep-FQ. G) L-y10b26 InvD displacement of L-y07b26 SIncR with L-d06z16 Rep-FQ. H) L-y10b26 InvD displacement of L-y10b26 SIncR with L-d06z16 Rep-FQ. The legend given in A corresponds to all subplots.

As well as fitting each curve individually we also calculated a global fit which estimated a single value for $k_{\text{eff}}$ for all 9 curves for each set of strands (i.e. across 3 replicates and 3 different $[Inv]_{exp}$ concentrations) (Table S13). Global fits were applied to kinetic data from all

DNA>DNA (Figure S16), RNA>DNA (Figure S17) and $DNA_{py}$>$RNA_{py}$ (Figure S18) strand displacement reactions. As above we employed the python function jax.experimental.ode.odeint to integrate the ODE model. We set initial, fixed concentrations of 0 nM for [*SInv*], [*Inc*], [*FInc*], [*Q*] and used the estimated concentrations of *SInc* ($[SInc]_{est0}$) and *FQ* ($[FQ]_{rctest0}$) as initial concentrations. We utilised scipy.optimize.curve_fit as for global fits in supplementary note 7.3. To fit a single, global estimate of $k_{\mathrm{eff}}$ for each set of strands we fixed $k_{\mathrm{rep}}$ at the estimated value from the corresponding reporter characterisation (global jackknife estimate) and fixed $[Inv]_0$ at $[Inv]_{est0}$. We used the same initial $k_{\mathrm{eff}}$ predictions and lower and upper bounds as for the individual fits. Notably, for fast reactions (all systems with $\gamma$ = 7nt or 10nt and DNA>RNA reactions with $\gamma$ = 6nt) we adapted the $[Inv]_0$ inputs. We noted a sudden increase in the fluorescence between the measurement of reaction kinetics (measurement 2 for DNA>DNA and RNA>DNA strand displacement (Table S9) or measurement 3 for DNA>RNA strand displacement (Table S10)) and the steady state measurement (measurement 3 for DNA>DNA and RNA>DNA strand displacement or measurement 4 for DNA>RNA strand displacement) for these fast reactions. This jump in fluorescence was not observed in the background negative control nor the positive control and so was not eliminated during processing. Given that we do not fit $[Inv]_0$ in the global fitting protocol this jump led to poor fits to the kinetic data. Therefore, for these reactions we set $[Inv]_0$ to the fluorescence value of the final timepoint in the reaction kinetics (measurement 2 for DNA>DNA and RNA>DNA strand displacement and measurement 3 for DNA>RNA strand displacement). In order to obtain a sensible error for the $k_{\mathrm{eff}}$ estimates we used a jackknife (leave-one-out) approach. We estimated the global $k_{\mathrm{eff}}$ each time leaving out a different one of the 9 fluorescence curves. These jackknife replicates were then inverse log transformed and the final estimate of $k_{\mathrm{eff}}$ was calculated according to equation (*S35*), replacing $k_{\mathrm{rep}}$ by $k_{\mathrm{eff}}$. Finally, we calculated the jackknife standard error according to equation (*S36*), replacing $k_{\mathrm{rep}}$ with $k_{\mathrm{eff}}$. The global jackknife $k_{\mathrm{eff}}$ estimates for each experiment are used for the further parameterisation of the free-energy landscape model.

Fitted parameters are reported in Table S13 and the fits themselves are plotted in Figures S16-S18.

| Full strand displacement system | Jackknife global $k_{\mathrm{eff}}$ ($M^{-1}$ $s^{-1}$) | Jackknife standard error ($M^{-1}$ $s^{-1}$) |
|---|---|---|
| L-y10b26 InvD + L-y04b11 SIncD + L-d06z16 Rep-FQ | $5.655 \cdot 10^4$ | $0.056 \cdot 10^4$ |
| L-y10b26 InvD + L-y04b16 SIncD + L-d06z16 Rep-FQ | $2.823 \cdot 10^4$ | $0.032 \cdot 10^4$ |
| L-y10b26 InvD + L-y04b21 SIncD + L-d06z16 Rep-FQ | $5.344 \cdot 10^3$ | $0.179 \cdot 10^3$ |
| L-y10b26 InvD + L-y04b26 SIncD + L-d06z16 Rep-FQ | $3.553 \cdot 10^3$ | $0.082 \cdot 10^3$ |
| L-y10b26 InvD + L-y05b26 SIncD + L-d06z16 Rep-FQ | $1.686 \cdot 10^5$ | $0.015 \cdot 10^5$ |
| L-y10b26 InvD + L-y06b26 SIncD + L-d06z16 Rep-FQ | $5.938 \cdot 10^5$ | $0.150 \cdot 10^5$ |
| L-y10b26 InvD + L-y07b26 SIncD + L-d06z16 Rep-FQ | $3.583 \cdot 10^6$ | $1.000 \cdot 10^6$ |
| L-y10b26 InvD + L-y10b26 SIncD + L-d06z16 Rep-FQ | $6.389 \cdot 10^6$ | $2.305 \cdot 10^6$ |
| L-y10b26 InvR + L-y04b11 SIncD + L-d06z16 Rep-FQ | $1.000 \cdot 10^4$ | $0.015 \cdot 10^4$ |
| L-y10b26 InvR + L-y04b16 SIncD + L-d06z16 Rep-FQ | $6.156 \cdot 10^3$ | $0.040 \cdot 10^3$ |
| L-y10b26 InvR + L-y04b21 SIncD + L-d06z16 Rep-FQ | $4.347 \cdot 10^2$ | $0.036 \cdot 10^2$ |
| L-y10b26 InvR + L-y04b26 SIncD + L-d06z16 Rep-FQ | $1.860 \cdot 10^2$ | $0.098 \cdot 10^2$ |
| L-y10b26 InvR + L-y05b26 SIncD + L-d06z16 Rep-FQ | $7.773 \cdot 10^3$ | $0.084 \cdot 10^3$ |
| L-y10b26 InvR + L-y06b26 SIncD + L-d06z16 Rep-FQ | $1.060 \cdot 10^4$ | $0.022 \cdot 10^4$ |
| L-y10b26 InvR + L-y07b26 SIncD + L-d06z16 Rep-FQ | $1.966 \cdot 10^6$ | $0.256 \cdot 10^6$ |
| L-y10b26 InvR + L-y10b26 SIncD + L-d06z16 Rep-FQ | $4.405 \cdot 10^6$ | $0.188 \cdot 10^6$ |
| L-y10b26 InvD + L-y04b11 SIncR + L-d06z16 Rep-FQ | $2.176 \cdot 10^5$ | $0.055 \cdot 10^5$ |
| L-y10b26 InvD + L-y04b16 SIncR + L-d06z16 Rep-FQ | $1.747 \cdot 10^5$ | $0.075 \cdot 10^5$ |
| L-y10b26 InvD + L-y04b21 SIncR + L-d06z16 Rep-FQ | $2.395 \cdot 10^4$ | $0.054 \cdot 10^4$ |
| L-y10b26 InvD + L-y04b26 SIncR + L-d06z16 Rep-FQ | $3.842 \cdot 10^4$ | $0.031 \cdot 10^4$ |
| L-y10b26 InvD + L-y05b26 SIncR + L-d06z16 Rep-FQ | $6.777 \cdot 10^5$ | $0.135 \cdot 10^5$ |

| | | |
|---|---|---|
| L-y10b26 InvD + L-y06b26 SIncR + L-d06z16 Rep-FQ | $2.944 \cdot 10^6$ | $0.045 \cdot 10^6$ |
| L-y10b26 InvD + L-y07b26 SIncR + L-d06z16 Rep-FQ | $4.811 \cdot 10^6$ | $0.479 \cdot 10^6$ |
| L-y10b26 InvD + L-y10b26 SIncR + L-d06z16 Rep-FQ | $1.210 \cdot 10^7$ | $0.063 \cdot 10^7$ |
| H-y06b21 InvD + H-y04b11 SIncD + H-d08z16 Rep-FQ | $1.887 \cdot 10^4$ | $0.058 \cdot 10^4$ |
| H-y06b21 InvD + H-y04b16 SIncD + H-d08z16 Rep-FQ | $1.674 \cdot 10^4$ | $0.037 \cdot 10^4$ |
| H-y06b21 InvD + H-y04b21 SIncD + H-d08z16 Rep-FQ | $1.153 \cdot 10^4$ | $0.010 \cdot 10^4$ |
| H-y06b21 InvR + H-y04b11 SIncD + H-d08z16 Rep-FQ | $9.670 \cdot 10^4$ | $0.044 \cdot 10^4$ |
| H-y06b21 InvR + H-y04b16 SIncD + H-d08z16 Rep-FQ | $1.036 \cdot 10^5$ | $0.016 \cdot 10^5$ |
| H-y06b21 InvR + H-y04b21 SIncD + H-d08z16 Rep-FQ | $1.640 \cdot 10^5$ | $0.063 \cdot 10^5$ |

**Table S13. Global $k_{\mathrm{eff}}$ estimates across full strand displacement reactions.** Global $k_{\mathrm{eff}}$ estimates and standard errors across 3 replicates and 3 different expected *Inv* ($[Inv]_{exp}$ = 4-8 nM) concentrations for RNA>DNA, DNA>DNA and DNA>RNA strand displacement reactions. Expected reporter ($[FQ]_{exp}$) and incumbent-substrate complex ($[SInc]_{exp}$) concentrations of 15 nM and 10 nM were used for all experiments, respectively. $k_{\mathrm{eff}}$ estimates were obtained by fitting an ODE model of the full strand displacement reaction all fluorescence curves simultaneously for each experiment. $k_{\mathrm{eff}}$ estimates and standard errors were calculated using a jackknife (leave-one-out) approach.

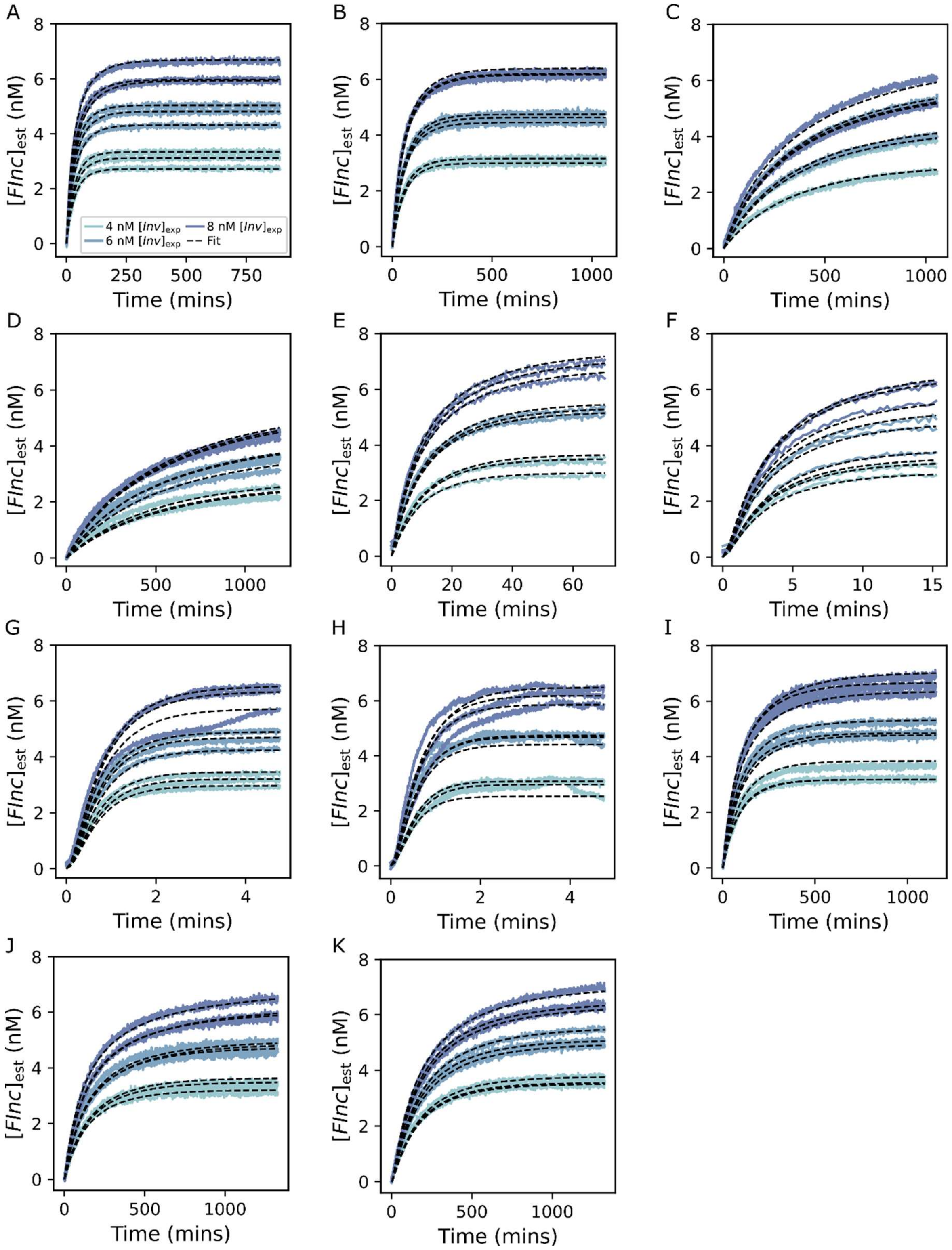

**Figure S16. Global fits to fluorescent traces in DNA>DNA strand displacement reactions.** Global fit of the ODE model to processed fluorescent data for each DNA>DNA set of strands. Global fits for each set of strands were derived from 3 replicates at 3 different expected *Inv* ($[Inv]_{exp}$ = 4-8 nM) concentrations. Expected reporter ($[FQ]_{exp}$) and incumbent-substrate complex ($[SInc]_{exp}$) concentrations of 15 nM and 10 nM were used for all experiments, respectively. For $DNA_{py}$>$DNA_{py}$ 4 different branch migration domain lengths (11-26nt) and 5 different invader toehold lengths (4-10nt) were tested. For $DNA_{pu}$>$DNA_{pu}$ 3 different branch migration domain lengths (11-21nt) were tested. $k_{eff}$ estimates and standard errors were calculated using a jackknife (leave-one-out) approach. Black, dotted lines represent fit of ODE model and solid, coloured lines represent the normalised experimental fluorescent data. A) L-y10b26 InvD displacement of L-y04b11 SIncD with L-d06z16 Rep-FQ. B) L-y10b26 InvD displacement of L-y04b16 SIncD with L-d06z16 Rep-FQ. C) L-y10b26 InvD displacement of L-y04b21 SIncD with L-d06z16 Rep-FQ. D) L-y10b26 InvD displacement of L-y04b26 SIncD with L-d06z16 Rep-FQ. E) L-y10b26 InvD displacement of L-y05b26 SIncD with L-d06z16 Rep-FQ. F) L-y10b26 InvD displacement of L-y06b26 SIncD with L-d06z16 Rep-FQ. G) L-y10b26 InvD displacement of L-y07b26 SIncD with L-d06z16 Rep-FQ. H) L-y10b26 InvD displacement of L-y10b26 SIncD with L-d06z16 Rep-FQ. I) H-y06b21 InvD displacement of H-y04b11 SIncD with H-d08z16 Rep-FQ. J) H-y06b21 InvD displacement of H-y04b16 SIncD with H-d08z16 Rep-FQ. K) H-y06b21 InvD displacement of H-y04b21 SIncD with H-d08z16 Rep-FQ. The legend given in A corresponds to all subplots.

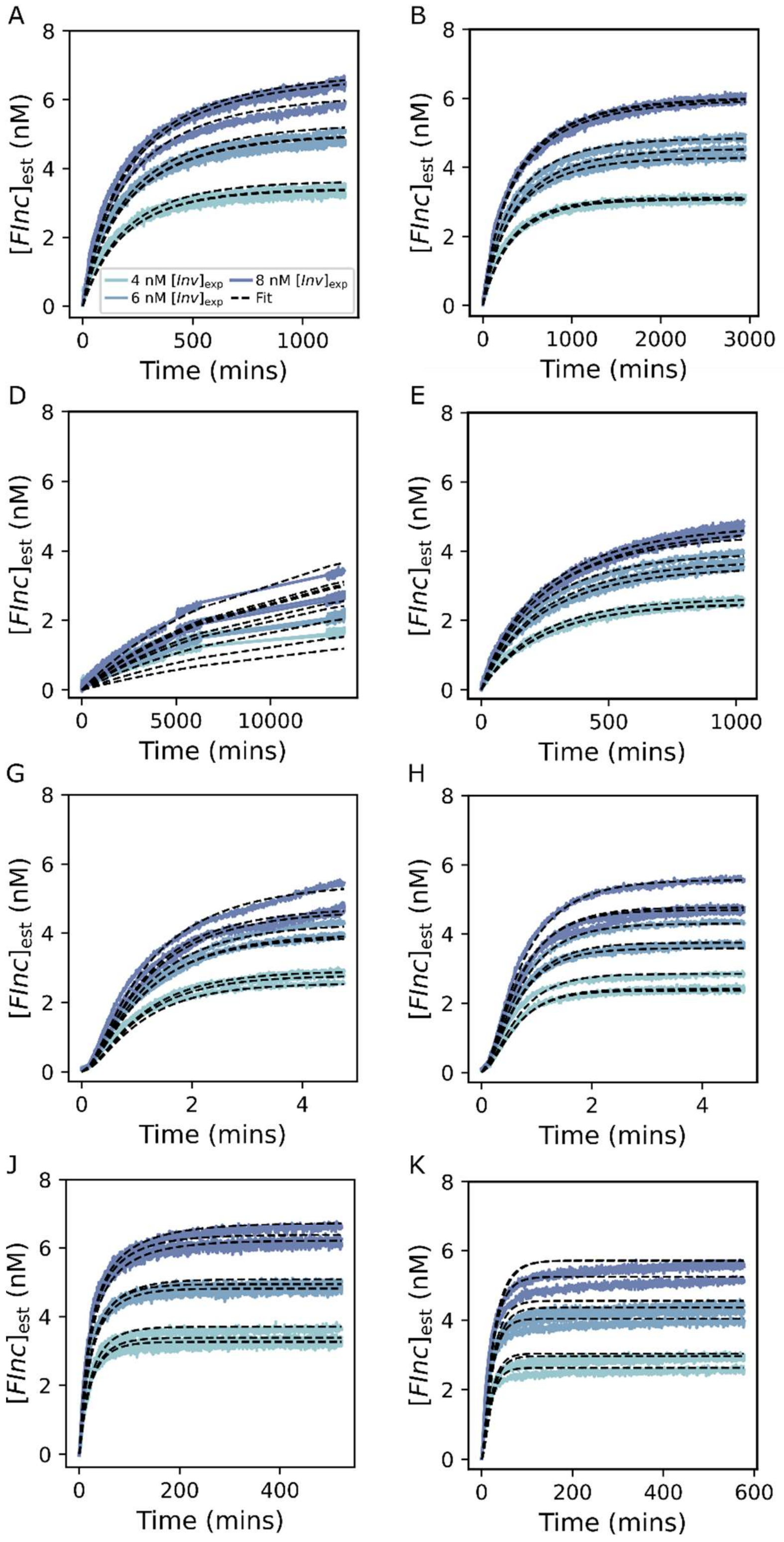

**Figure S17. Global fits to fluorescent traces in RNA>DNA strand displacement reactions.** Global fit of the ODE model to processed fluorescent data for each RNA>DNA set of strands. Global fits for each set of strands were derived from 3 replicates at 3 different expected *Inv* ($[Inv]_{exp}$ = 4-8 nM) concentrations. Expected reporter ($[FQ]_{exp}$) and incumbent-substrate complex ($[SInc]_{exp}$) concentrations of 15 nM and 10 nM were used for all experiments, respectively. For $RNA_{py}$>$DNA_{py}$ 4 different branch migration domain lengths (11-26nt) and 5 different invader toehold lengths (4-10nt) were tested. For $RNA_{pu}$>$DNA_{pu}$ 3 different branch migration domain lengths (11-21nt) were tested. $k_{eff}$ estimates and standard errors were calculated using a jackknife (leave-one-out) approach. Black, dotted lines represent fit of ODE model and solid, coloured lines represent the normalised experimental fluorescent data. A) L-y10b26 InvR displacement of L-y04b11 SIncD with L-d06z16 Rep-FQ. B) L-y10b26 InvR displacement of L-y04b16 SIncD with L-d06z16 Rep-FQ. C) L-y10b26 InvR displacement of L-y04b21 SIncD with L-d06z16 Rep-FQ. D) L-y10b26 InvR displacement of L-y04b26 SIncD with L-d06z16 Rep-FQ. E) L-y10b26 InvR displacement of L-y05b26 SIncD with L-d06z16 Rep-FQ. F) L-y10b26 InvR displacement of L-y06b26 SIncD with L-d06z16 Rep-FQ. G) L-y10b26 InvR displacement of L-y07b26 SIncD with L-d06z16 Rep-FQ. H) L-y10b26 InvR displacement of L-y10b26 SIncD with L-d06z16 Rep-FQ. I) H-y06b21 InvR displacement of H-y04b11 SIncD with H-d08z16 Rep-FQ. J) H-y06b21 InvR displacement of H-y04b16 SIncD with H-d08z16 Rep-FQ. K) H-y06b21 InvR displacement of H-y04b21 SIncD with H-d08z16 Rep-FQ. The legend given in A corresponds to all subplots.

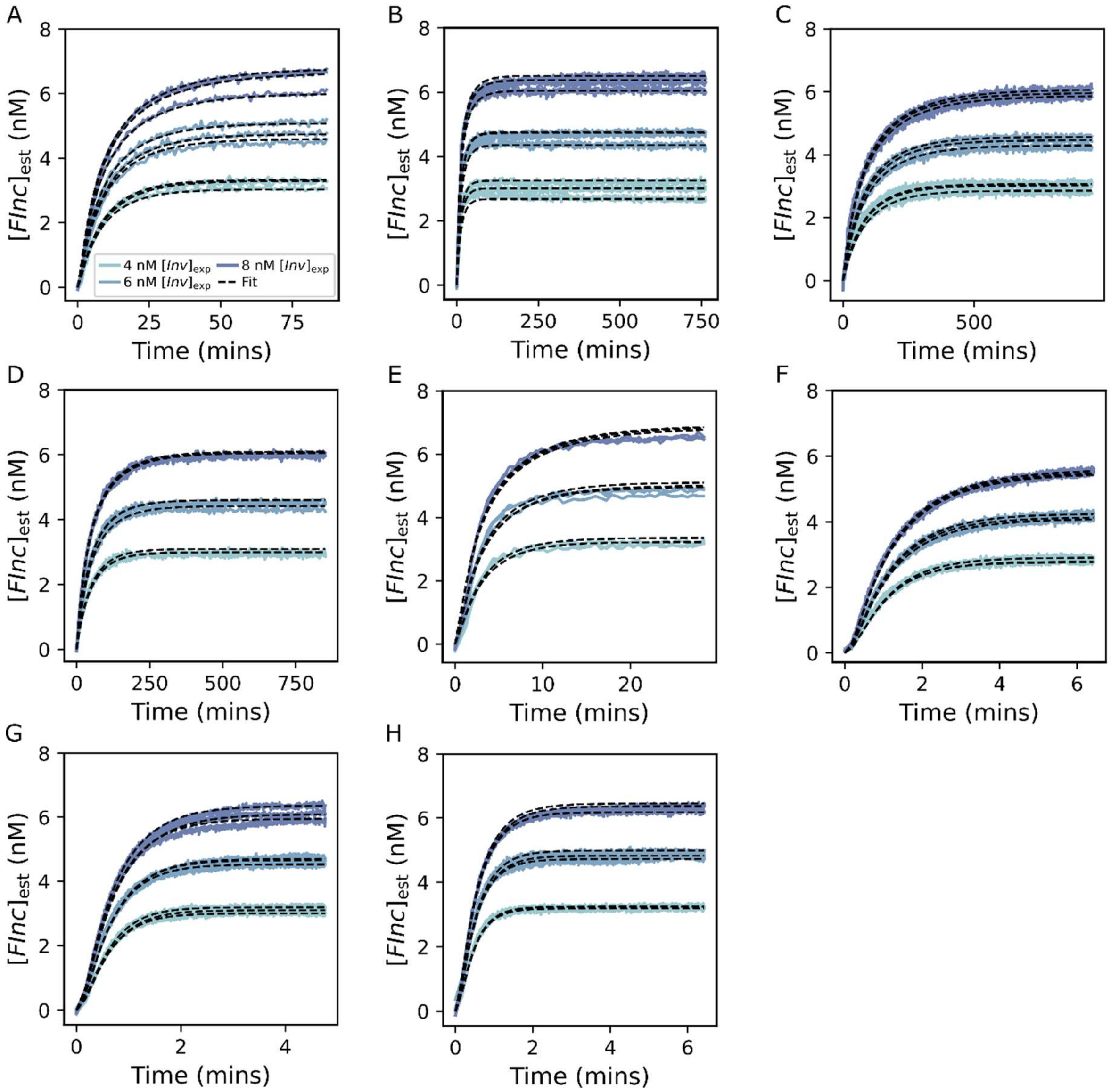

**Figure S18. Global fits to fluorescent traces in DNA>RNA strand displacement reactions.** Global fit of the ODE model to processed fluorescent data for each DNA>RNA set of strands. Global fits for each set of strands were derived from 3 replicates at 3 different expected *Inv* ($[Inv]_{exp}$ = 4-8 nM) concentrations. Expected reporter ($[FQ]_{exp}$) and incumbent-substrate complex ($[SInc]_{exp}$) concentrations of 15 nM and 10 nM were used for all experiments, respectively. For $DNA_{py}$>$RNA_{py}$ 4 different branch migration domain lengths (11-26nt) and 5 different invader toehold lengths (4-10nt) were tested. $k_{eff}$ estimates and standard errors were calculated using a jackknife (leave-one-out) approach. Black, dotted lines represent fit of ODE model and solid, coloured lines represent the normalised experimental fluorescent data. A) L-y10b26 InvD displacement of L-y04b11 SIncR with L-d06z16 Rep-FQ. B) L-y10b26 InvD displacement of L-y04b16 SIncR with L-d06z16 Rep-FQ. C) L-y10b26 InvD displacement of L-y04b21 SIncR with L-d06z16 Rep-FQ. D) L-y10b26 InvD displacement of L-y04b26 SIncR with L-d06z16 Rep-FQ. E) L-y10b26 InvD displacement of L-y05b26 SIncR with L-d06z16 Rep-FQ. F) L-y10b26 InvD displacement of L-y06b26 SIncR with L-d06z16 Rep-FQ. G) L-y10b26 InvD displacement of L-y07b26 SIncR with L-d06z16 Rep-FQ. H) L-y10b26 InvD displacement of L-y10b26 SIncR with L-d06z16 Rep-FQ. The legend given in A corresponds to all subplots.

*Supplementary note 8.4. Leak in $DNA_{pu}$>$RNA_{pu}$ strand displacement reactions*

For the DNA>RNA strand displacement reactions which exhibited an undesired leak reaction ($DNA_{pu}$>$RNA_{pu}$) we were forced to adapt the ODE model to capture this leak reaction. We define a system of ODEs to describe the leak reaction within the negative control for which 0 nM *Inv* is added (equations (*8-10*)). For this purpose, we define an additional rate constant, $k_{\mathrm{leak}}$, which is the rate constant for the leak reaction between *SInc* and *FQ*. Using only the fluorescent data from the negative control we were able to obtain an estimate for $k_{\mathrm{leak}}$ (Figure S19). As in supplementary note 7.3, we used scipy.integrate.odeint and scipy.optimize.curve_fit to fit the ODE model to the experimental data, however in this case we fitted $k_{\mathrm{leak}}$ while keeping $k_{\mathrm{rep}}$ as a fixed constant. We set initial concentrations of 0 nM for [*FInc*] and [*Q*] and used the estimated concentrations of *SInc* ($[SInc]_{\mathrm{est0}}$) and *FQ* ($[FQ]_{\mathrm{est0}}$) as initial concentrations. We used an initial estimate of $10^2$ $M^{-1}$ $s^{-1}$ for $k_{\mathrm{leak}}$ with lower and upper bounds of $10^0$ $M^{-1}$ $s^{-1}$ and $10^8$ $M^{-1}$ $s^{-1}$, respectively. We also allow $[exInc]_{\mathrm{fit0}}$ to be a flexible parameter, with an initial estimate of $[exInc]_{\mathrm{est0}}$ from the fluorescence of measurement 2 and lower and upper bounds of $[exInc]_{\mathrm{est0}}/2$ and $2[exInc]_{\mathrm{est0}}$, respectively (Table S11). For each system we calculated the mean $k_{\mathrm{leak}}$ value across the 3 negative control replicates (Table S14). Fits are shown in Figure S19.

| Leak reaction system | Mean $k_{\mathrm{leak}}$ ($M^{-1}$ $s^{-1}$) | Standard error ($M^{-1}$ $s^{-1}$) |
|---|---|---|
| High purine y04b11 SIncR + d06z16 Rep-FQ | $1.270 \cdot 10^2$ | $0.106 \cdot 10^2$ |
| High purine y04b16 SIncR + d06z16 Rep-FQ | $6.855 \cdot 10^1$ | $0.041 \cdot 10^2$ |
| High purine y04b21 SIncR + d06z16 Rep-FQ | $1.245 \cdot 10^2$ | $0.042 \cdot 10^2$ |

**Table S14. $k_{\mathrm{leak}}$ estimates from averaging over individual leak characterisation reactions.** Mean $k_{\mathrm{leak}}$ estimates and standard errors from individual fits of 3 replicates for $DNA_{pu}$>$RNA_{pu}$ strand displacement reactions. $k_{\mathrm{leak}}$ estimates were obtained by fitting an ODE model of the leak reaction to each individual fluorescence curve.

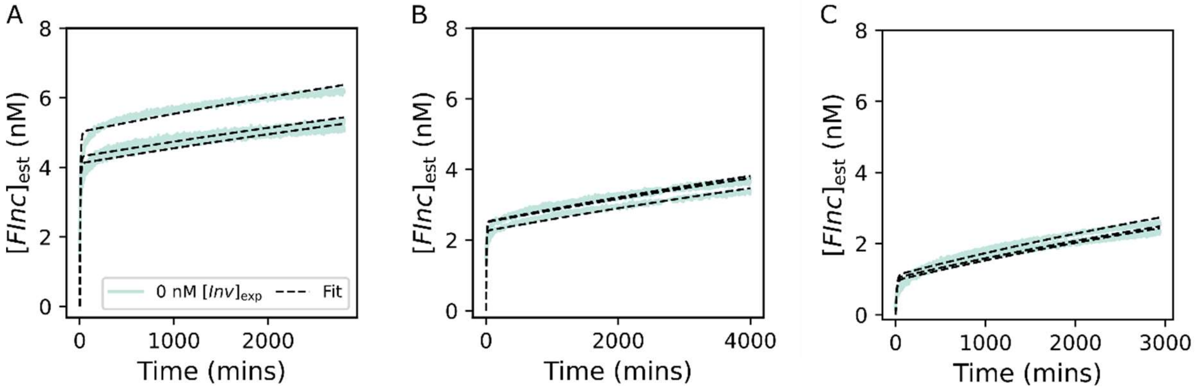

**Figure S19. Individual fits to fluorescent traces in leak characterisation reactions.** Individual fits of ODE model to normalised negative control fluorescent data from each leak reaction characterisation experiment. Expected reporter ($[FQ]_{exp}$) and incumbent-substrate complex ($[SInc]_{exp}$) concentrations of 15 nM and 10 nM were used for all experiments, respectively. An excess *Inc* was added in all experiments ($[exInc]_{exp}$ = 2 nM). 3 different branch migration domain lengths (11-21nt) were tested. Each experiment was performed in triplicate. Black, dotted lines represent fit of ODE model and solid, coloured lines represent the normalised experimental fluorescent data. A) H-y04b11 SIncR and excess H-y04b11 IncR leak interaction with H-d06z16 Rep-FQ. B) H-y04b16 SIncR and excess H-y04b16 IncR leak interaction with H-d06z16 Rep-FQ and C) H-y04b21 SIncR and excess H-y04b21 IncR leak interaction with H-d06z16 Rep-FQ.

We then calculated individual fits for the $DNA_{pu}$>$RNA_{pu}$ reactions of interest, taking this leak into account. In this case we used an adapted ODE model (equation (*11-14*)) which captures the leak reaction (Figure S20). We fixed $k_{\mathrm{leak}}$ at the estimated values from the corresponding negative control reactions (Table S14). We also fixed $k_{\mathrm{rep}}$ at the values estimated from the corresponding reporter characterisation reactions (global jackknife estimate). We set initial, fixed concentrations of 0 nM for [*SInv*], [*Inc*], [*FInc*], [*Q*] and used $[SInc]_{est0}$ and $[FQ]_{rctest0}$ as fixed reactant inputs into the ODE integrator function. Unlike the other strand displacement systems we were unable to explicitly estimate $[Inv]_{est0}$ from the steady state fluorescence as we used an excess of $[Inv]_0$ for these reactions. We therefore used initial estimates of $[Inv]_{est0}$ values equal to $[Inv]_{exp}$ (40 nM, 60 nM and 80 nM). For the individual fits we allowed $[Inv]_0$ to be a flexible parameter ($[Inv]_{fit0}$) with bounds of $[Inv]_{est0}/2$ and $2[Inv]_{est0}$, and fitted it alongside $k_{\mathrm{eff}}$. The mean $k_{\mathrm{eff}}$ estimate and standard error obtained are given in Table S15, and the fits are shown in Figure S20. We were unable to effectively obtain a global estimate of $k_{\mathrm{eff}}$ for these systems as we could not effectively estimate $[Inv]_{est0}$ which is a critical input for global estimation.

| Full strand displacement system | Mean $k_{\mathrm{eff}}$ (M-1 s-1) | Standard error (M-1 s-1) |
|---|---|---|
| H-y04b11 SIncR + H-d06z16 Rep-FQ | $5.821 \cdot 10^2$ | $1.026 \cdot 10^2$ |
| H-y04b16 SIncR + H-d06z16 Rep-FQ | $1.217 \cdot 10^2$ | $0.249 \cdot 10^2$ |
| H-y04b21 SIncR + H-d06z16 Rep-FQ | $8.137 \cdot 10^1$ | $0.216 \cdot 10^2$ |

**Table S15. $k_{\mathrm{eff}}$ estimates from averaging over individual $DNA_{pu}$>$DNA_{pu}$ strand displacement reactions with leak.** Mean $k_{\mathrm{leak}}$ estimates and standard errors from individual fits of 3 replicates for $DNA_{pu}$>$RNA_{pu}$ strand displacement reactions. $k_{\mathrm{eff}}$ estimates were obtained by fitting an ODE model of the full $DNA_{pu}$>$RNA_{pu}$ strand displacement reaction to each individual fluorescence curve.

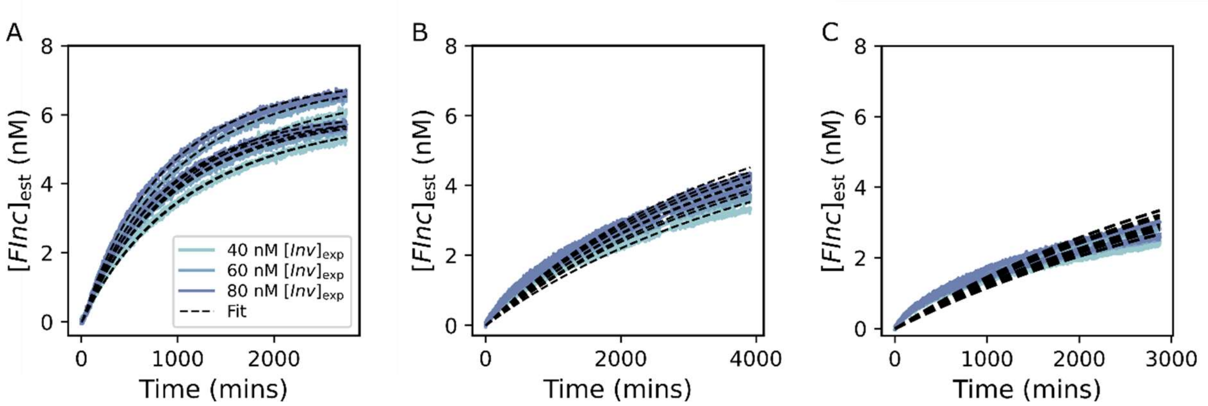

**Figure S20. Individual fits to fluorescent traces in full $DNA_{pu}$>$RNA_{pu}$ strand displacement reactions with leak.** Individual fits of ODE model to normalised fluorescent data for each $DNA_{pu}$>$RNA_{pu}$ strand displacement experiment. Each experiment was performed in triplicate and across 3 different *Inv* ($[Inv]_{exp}$ = 40-80 nM) concentrations. Expected reporter ($[FQ]_{exp}$) and incumbent-substrate complex ($[SInc]_{exp}$) concentrations of 15 nM and 10 nM were used for all experiments, respectively. 3 different branch migration domain lengths (11-21nt) were tested. Black, dotted lines represent fit of ODE model and solid, coloured lines represent the normalised experimental fluorescent data. A) H-y06b21 InvD displacement of H-y04b11 SIncR with H-d06z16 Rep-FQ. B) H-y06b21 InvD displacement of H-y04b16 SIncR with H-d06z16 Rep-FQ. C) H-y06b21 InvD displacement of H-y04b21 SIncR with H-d06z16 Rep-FQ.

### Supplementary note 9. Parameterisation of the free-energy landscape model

*Supplementary note 9.1. Details of parameterisation method*

We define 3 separate functions, one for each type of strand displacement reaction we investigated: DNA>DNA; RNA>DNA and DNA>RNA. These functions take the following inputs: branch migration domain length, invader toehold length, incumbent toehold length, initial invader concentration, temperature (in K), Boltzmann constant, 5 free-energy parameters ($\Delta G_{\mathrm{assoc}}$, $\Delta G_{\mathrm{bp}}$, $\Delta G_{\mathrm{p}}$, $\Delta G_{\mathrm{bm}}$, $\Delta G_{\mathrm{rd}}$) and a rate parameter ($k_{\mathrm{bp}}$). With these inputs the function can fully specify the free-energy landscape model and determine the first passage time for the system and output an effective rate constant ($k_{\mathrm{eff}}$). Notably, the output of these functions is a log-transformed $k_{\mathrm{eff}}$ value in order to facilitate comparison with the experimentally derived rate constants.

We define an additional function that computes the residuals ($R$) between experimentally-derived $k_{\mathrm{eff}}$ values (log-transformed) and the model predicted values ($\widehat{k_{\mathrm{eff}}}$) according to the following equation

$$R_{(i)} = \frac{\left(\widehat{k_{\mathrm{eff}(i)}} - k_{\mathrm{eff}(i)}\right)}{SE\left(k_{\mathrm{eff}(i)}\right)}. \qquad (S41)$$

This function takes the global jackknife estimates of $k_{\mathrm{eff}}$ for each experimental system and the standard error of estimated $k_{\mathrm{eff}}$ across individual curves as inputs ($SE(k_{\mathrm{eff}})$). Notably, for $DNA_{pu}$>$RNA_{pu}$ strand displacement systems we used the mean of the individual $k_{\mathrm{eff}}$ estimates (Table S15) as we were unable to obtain suitable global $k_{\mathrm{eff}}$ estimates. Further, we used the standard error across these individual fits as $SE(k_{\mathrm{eff}})$ for these strand displacement reactions. Finally, we employed the python function scipy.optimize.least_squares to identify the optimal parameter values which minimise the squared residuals between the model-predicted $k_{\mathrm{eff}}$ and the experimental $k_{\mathrm{eff}}$ values. We provide initial parameter estimates as well as lower and upper bounds for each flexible parameter.

*Supplementary note 9.2. Calculation of error in model-predicted $k_{eff}$ values*

Introducing each estimated parameter value back into the free-energy landscape model we extracted the model predicted $k_{\mathrm{eff}}$ values. We then calculated the weighted residual sum of squares ($WRSS$) between the experimentally derived $k_{\mathrm{eff}}$ values and the model-predicted $\widehat{k_{\mathrm{eff}}}$ values for each observation $i$ according to

$$WRSS = \sum_{i=1}^{n} \left(k_{\mathrm{eff}(i)} - \widehat{k_{\mathrm{eff}(i)}}\right)^2. \qquad (S42)$$

We calculated the 95% confidence interval for each model-predicted $\widehat{k_{\mathrm{eff}}}$ value. We calculated updated $k_{\mathrm{eff}}$ values based on the $WRSS$ and the degrees of freedom ($df$), calculated as the number of parameters subtracted from number of observations

$$k_{\mathrm{eff}(i)} = \widehat{k_{\mathrm{eff}(i)}} + \sqrt{\frac{WRSS}{df}} \cdot X \sim N(0, 1), \qquad (S43)$$

where $X$ is a random variable sampled from a normal distribution with a mean of 0 and a variance of 1. We introduce the new weighted values of $k_{\mathrm{eff}}$ as arguments into scipy.optimize.least_squares which produces a distribution of new parameter values, from which we predict novel $k_{\mathrm{eff}}$ values. Running this simulation 100 times and ordering the outputs of $k_{\mathrm{eff}}$, we identified the upper and lower 95% confidence interval for the model-predicted $\widehat{k_{\mathrm{eff}}}$ values.

*Supplementary note 9.3. Alternative parameterisation of free-energy and rate parameters*

At first we attempted to estimate all of the free-energy and rate parameters in the model from the experimental data for $RNA_{py}$>$DNA_{py}$, $DNA_{py}$>$DNA_{py}$ and $DNA_{py}$>$RNA_{py}$ strand displacement. Initial parameter estimates for scipy.optimize.least_squares were extracted from Irmisch et al. (2020). The final output parameter estimates from this parameterisation were not in good agreement with previous experimental studies and in particular $\Delta G_{\mathrm{p}}$ could not be estimated (Table S16). We noted that the available data didn't strongly constrain all parameters, with similar fits arising when subsets of parameters were varied together. We therefore fixed some parameters at the values from Irmisch et al. (2020) to allow for appropriate parameter estimation. By fixing a subset of the parameters ($\Delta G_{\mathrm{bp}}$, $\Delta G_{\mathrm{assoc}}$, and $\Delta G_{\mathrm{p}}$) we achieved better agreement with prior studies (3, 4), while maintaining a good fit to the experimental data. $k_{\mathrm{bp}}$ was log-transformed as this parameter varies over orders of magnitude. We used initial parameter estimates of 7.4 $k_{\mathrm{B}}T$, $\log_{10}(5.9 \cdot 10^{7})$ $s^{-1}$ and 0.5 $k_{\mathrm{B}}T$ for $\Delta G_{\mathrm{bm}}$, $k_{\mathrm{bp}}$ and $\Delta G_{\mathrm{rd}}$, respectively. While the lower and upper bounds were set at [0, 15]; [0, 15]; and [-5; 5] for $\Delta G_{\mathrm{bm}}$, $k_{\mathrm{bp}}$ and $\Delta G_{\mathrm{rd}}$, respectively.

| **Data set** | $\Delta G_{\mathrm{bp}}$ $(k_{\mathrm{B}}T)$ | $\Delta G_{\mathrm{assoc}}$ $(k_{\mathrm{B}}T)$ | $\Delta G_{\mathrm{p}}$ $(k_{\mathrm{B}}T)$ | $\Delta G_{\mathrm{bm}}$ $(k_{\mathrm{B}}T)$ | $k_{\mathrm{bp}}$ ($s^{-1}$) | $\Delta G_{\mathrm{rd}}$ $(k_{\mathrm{B}}T)$ |
|---|---|---|---|---|---|---|
| This study | $-2.22 \pm 0.88$ | $0.61 \pm 31.1$ | - | $12.1 \pm 34.3$ | $1.5 \cdot 10^{7}$* | $0.15 \pm 0.10$ |
| Irmisch | $-2.52$ | $2.5 \pm 0.2$ | $3.5 \pm 0.2$ | $7.4 \pm 0.2$ | $(5.9 \pm 1.1) \cdot 10^{7}$ | - |
| Machinek | $-2.51$ | $4.0 \pm 0.2$ | $5.4 \pm 0.4$ | $8.5 \pm 0.3$ | $(20.6 \pm 0.6) \cdot 10^{7}$ | - |

**Table S16. Parameter estimates from alternative parameterisation of IEL model using low RNA purine data.** Estimates of free-energy and rate parameters that describe the free-energy landscape (IEL) model when all parameters are allowed to be flexible parameters. The IEL model was parameterised based on $RNA_{py}$>$DNA_{py}$, $DNA_{py}$>$DNA_{py}$ and $DNA_{py}$>$RNA_{py}$ strand displacement experimental kinetic data. *The 95% confidence interval for the estimated $k_{\mathrm{bp}}$ parameter was greater than $10^{10}$ $s^{-1}$ so is not explicitly quantified here. $\Delta G_{\mathrm{p}}$ could not be reliably estimated in this parameterisation. Previous studies of DNA>DNA strand displacement estimated significantly different parameter values for most parameters (3, 4).

For $RNA_{pu}$>$DNA_{pu}$, $DNA_{pu}$>$DNA_{pu}$, $DNA_{pu}$>$RNA_{pu}$ strand displacement reactions we tried various combinations of fixed vs. flexible parameters to evaluate the effect on the model-predicted $k_{\mathrm{eff}}$ values (Table S17 and Figure S21). In all cases $\Delta G_{\mathrm{rd}}$ was less than 0 $k_{\mathrm{B}}T$. The choice of allowing $k_{\mathrm{bp}}$ and $\Delta G_{\mathrm{rd}}$ only to vary allowed a reasonable fit to data while keeping the parameters at sensible values.

| $\Delta G_{\mathrm{bp}}$ ($k_{\mathrm{B}}T$) | $\Delta G_{\mathrm{assoc}}$ ($k_{\mathrm{B}}T$) | $\Delta G_{\mathrm{p}}$ ($k_{\mathrm{B}}T$) | $\Delta G_{\mathrm{bm}}$ ($k_{\mathrm{B}}T$) | $k_{\mathrm{bp}}$ ($s^{-1}$) | $\Delta G_{\mathrm{rd}}$ ($k_{\mathrm{B}}T$) |
|---|---|---|---|---|---|
| $-2.52^{*}$ | $2.5^{*}$ | $3.5^{*}$ | $10.1 \pm 0.4$ | $6.4 \cdot 10^{7*}$ | $-0.25 \pm 0.07$ |
| $-2.52^{*}$ | $2.5^{*}$ | $3.5^{*}$ | $5.8 \pm 0.3$ | $1.8\ (CI{:}\ 1.6;\ 2.0) \cdot 10^{6}$ | $-0.34 \pm 0.03$ |
| $-2.52^{*}$ | $2.5^{*}$ | $3.5^{*}$ | $9.3^{*}$ | $6.4 \cdot 10^{7*}$ | $-0.15 \pm 0.03$ |

**Table S17. Parameter estimates from alternative parameterisation of IEL model using high RNA purine data.** Estimates of free-energy and rate parameters that describe the free-energy landscape (IEL) model when varying flexible parameters and fixed parameters. The IEL model was parameterised using $RNA_{pu}$>$DNA_{pu}$, $DNA_{pu}$>$DNA_{pu}$ and $DNA_{pu}$>$RNA_{pu}$ strand displacement experimental kinetic data. The available data doesn't strongly constrain all parameters, with similar fits arising when subsets of parameters were varied together therefore parameters marked with an asterisk (*) were fixed at the values from parameterisation using low RNA purine content data.

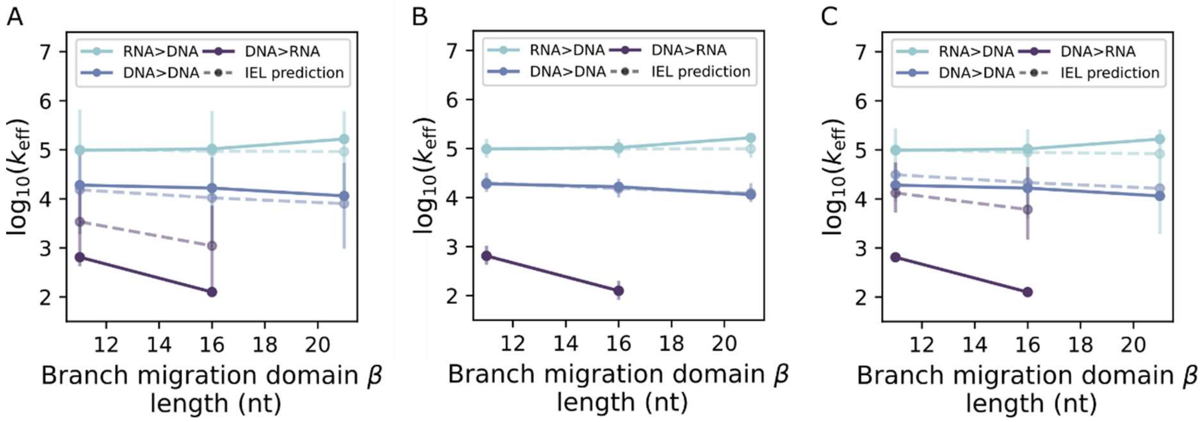

**Figure S21. Alternative IEL model predictions for $RNA_{pu}$>$DNA_{pu}$ and $DNA_{pu}$>$RNA_{pu}$ toehold exchange kinetics.** A) Predicted $k_{\mathrm{eff}}$ for $RNA_{pu}$>$DNA_{pu}$, $DNA_{pu}$>$DNA_{pu}$ and $DNA_{pu}$>$RNA_{pu}$ toehold exchange reactions across a range of branch migration domain lengths (11-21nt) for fixed $\gamma$ = 4nt and $\varepsilon$ = 4nt from parameterised IEL model (dashed lines) compared to experimentally-derived values (solid lines). Error bars show 95% confidence intervals of model predicted $k_{\mathrm{eff}}$ values.

**SUPPLEMENTARY INFORMATION REFERENCES**
